# Reassessing the diversity and distribution of African rope squirrels (*Funisciurus* Trouessart, 1880)

**DOI:** 10.1101/2025.08.25.672055

**Authors:** Pascal Baelo, Léa Fourchault, Nicolas Laurent, Claude Mande, Anne Laudisoit, Nicaise Amundala, Ganiyat Temidayo Saliu, Guy-Crispin Gembu, Joachim Mariën, Sophie Gryseels, Jan Hulselmans, Erik Verheyen

## Abstract

The diversity, taxonomy, distribution and ecology of sub-Saharan tree squirrels remain under-researched. This study aims to elucidate the diversity and distribution of rope squirrels, genus *Funisciurus* Trouessart 1880, in the Congo Basin. We assembled the most comprehensive genetic and morphometric data set to date, on a significant portion of the geographical and taxonomic range of these squirrels (470 specimens from seven provinces of the Democratic Republic of Congo). We identified six main taxa: a previously undescribed *Funisciurus* species (here, Fx), three allopatric taxa of *Funisciurus anerythrus* (Fc, FaRB and FaLB), and two sympatric genetically distinct taxa (FbLB1, FbLB2), potentially belonging to a *Funisciurus bayonii* species complex. We then mapped their distribution, showing that the Congo and Kasaï Rivers represent significant biogeographical barriers limiting the distribution of *Funisciurus* species. In addition to providing a baseline for research on the impact of climate change and habitat degradation on the abundance and distribution of rope squirrels, our findings support research on the role of rope squirrel species in the transmission of pathogens of public health importance.

**Teaser text:** We studied the diversity and distribution of rope squirrels, genus *Funisciurus* Trouessart 1880, in the Congo Basin. By gathering the most comprehensive genetic and morphometric data to date on a significant part of the geographical and taxonomic range of these squirrels (specimens from seven provinces of the Democratic Republic of Congo), we identified six main taxa: one previously undescribed species, three allopatric taxa, and two genetically distinct sympatric taxa. We then mapped their distribution, showing that the Congo and Kasai rivers represent significant biogeographical barriers limiting the distribution of *Funisciurus* species. These results support research into the role of rope squirrel species in the transmission of pathogens of public health importance.

**Image for the online table of contents:** African rope squirrel

**Photo credit:** Bushnell photo trap set by Pascal Baelo

## INTRODUCTION

The Sciuridae family, which includes approximately 50 genera and 302 species (**Zachos et al., 2021, Steiner & Huettmann, 2023**), is distributed globally, with the exception of Antarctica. While some squirrel species are valued as a food source in certain regions (**Fa et al., 2009; Baelo et al., 2018**), they are increasingly recognized as important reservoirs and incidental hosts for zoonotic pathogens (**Han et al., 2015; e.g., Sklenovska et al., 2018; Schulze et al., 2020**). However, the phylogenetic, ecological, and life history characteristics of many squirrels, particularly those in sub-Saharan Africa, remain poorly understood. In Central Africa, the Sciuridae family is represented by eight genera, including the African pygmy squirrel (*Myosciurus pumilio*) and several species within the rope (*Funisciurus*), sun (*Heliosciurus*), ground (*Xerus*), bush (*Paraxerus*), palm (*Epixerus*), and giant (*Protoxerus*) genera (**Kingdon & Hoffmann, 2013; Thorington et al., 2012**). Four of these genera—*Funisciurus*, *Paraxerus*, *Heliosciurus*, and *Protoxerus*—are common across diverse habitats in the Democratic Republic of Congo (DRC), spanning dense rainforest to savannah ecosystems.

Unlike many large mammals that are highly susceptible to habitat disturbances, a few Sciuridae species have shown a remarkable capacity to adapt to human-altered environments. For instance, *Funisciurus anerythrus* has been observed at high population densities in palm oil plantations, reaching up to 500 individuals per square kilometer—far exceeding densities in primary lowland rainforests (**Tiee et al., 2018**). This adaptability heightens the frequency of human-squirrel interactions and raises concerns about potential zoonotic spillover risks, particularly in agricultural and peri-domestic settings (**Rimoin et al., 2017; Kumakamba et al., 2021**).

Recent studies underscore the role of squirrels in zoonotic disease transmission, for example bornavirus (**Hoffmann et al., 2015; Schulze et al., 2020**). Sub-Saharan squirrels, particularly those within the *Funisciurus* and *Heliosciurus* genera, have also been associated with the mpox virus (MPXV), of which a variant led to a global outbreak infecting over 80,000 people in 2022, emphasizing the importance of understanding the status of these squirrels as zoonotic reservoirs (**Chen et al., 2012; Curaudeau et al., 2023; Kinganda-Lusamaki et al., 2025**).

Studying the distribution and phylogenetic relationships of *Funisciurus* in sub-Saharan Africa could illuminate patterns of pathogen-host co-evolution within this genus. MPXV, for instance, comprises two main clades: the more virulent clade I and the less virulent clade II (**Nakazawa et al., 2015; Kinganda-Lusamaki et al., 2025**). Clade I MPXV occurs in the Congo Basin, and clade II in West Africa - though both clades have been exported via human infections to other regions in Africa and the rest of the world (**Otieno et al., 2025**). The distinct clades found to be infecting humans could be a reflection of genetically and geographically distinct natural reservoir(s). These geographically distinct clades may correlate with specific squirrel populations, suggesting possible co-evolutionary and biogeographical relationships affecting disease transmission dynamics.

Geographical barriers, particularly rivers, can influence the distribution of rodents and the spread of pathogens, limiting the ranges of arboreal mammals in West and Central Africa. While natural barriers such as the Volta, Sanaga and Congo rivers are known to limit the ranges of several small mammal species (**Kennis et al., 2011**), the influence of the Congo River on the distribution and evolutionary history of Sciuromorph rodents remains underexplored, leaving significant gaps in our understanding of their evolutionary history across the Congo Basin.

This study addresses these gaps by clarifying the diversity of *Funisciurus* taxa within the Congo Basin and assessing whether the Congo River serves as a significant geographic barrier affecting their distribution. Along with advancing biodiversity, evolutionary and biogeographic insights, our research provides critical data on the distribution of *Funisciurus* taxa across the Congo Basin, to serve as a baseline for comparison with the phylogeographies of zoonotic pathogens relevant for public health.

## 2. MATERIALS AND METHODS

### 2.1. SAMPLE COLLECTION

#### 2.1.1. SQUIRREL CAPTURE AND SAMPLING

we trapped and sampled squirrels from 2014 to 2022 in 17 localities across four provinces (Tshopo, Bas-Uele, Ituri and Tshuapa) in the Democratic Republic of Congo (DRC). Further, we used specimens from three other localities in three provinces (Kwilu, Kinshasa and Kongo Central), where squirrels were sampled in 1995 and preserved at the Royal Museum for Central Africa-Tervuren (Belgium) (**Supplementary Figure S1**).

We captured squirrels using traditional traps (**Supplementary Figure S2**), baited with a ripe palm nut and placed in different habitats (primary forests, secondary forests and fallow land), at heights ranging from 0.00 m (on the ground) to 7.37 m (tree branches) (average height =1.58m±1.29m). The various standard traps and techniques used to capture rodents (pitfall traps, Havahart traps, flap traps, Sherman traps) proved unsuitable or ineffective when applied to squirrels. Consequently, traditional traps were used, given their effectiveness in capturing tree squirrels.

The traditional traps were constructed from stems, flexible vines and rattan, which were wrapped around a branch to facilitate the animals’ movement towards the bait (**Supplementary Figure S2**). A wooden wedge was then attached to the wire to act as a trigger for the bait. The trap was baited to attract the animal and catch it by the neck. Traditional traps had one or two sliding knots, allowing squirrels to be caught in both directions of approach. Other traps were constructed from flexible rods and wires with a radius of 0.5 mm and rattan. These traps had a single noose and functioned in the same way as traps with two nooses.

We euthanized live captured squirrels with an overdose of isoflurane, adhering to the guidelines of the American Society of Mammalogists for euthanasia (**Sikes et al., 2016**), before collecting blood samples on Whatman filter paper and collecting tissue samples (tongue, kidney, liver, lung, spleen, and colon) and feces. These tissues were preserved either in 70-94% ethanol or in a DNA/RNA preservation solution (e.g. RNAlater or DNA/RNA Shield, Zymo Research). The preserved samples were sent to the Royal Belgian Institute of Natural Sciences (RBINS) and the University of Antwerp (Belgium) for genetic analysis. The carcasses were preserved in a 10% formalin solution and stored partly at the Centre de Surveillance de la Biodiversité in Kisangani (DRC) and partly at the University of Antwerp (Belgium). A list of all specimens is provided in Supplementary Table S1.

Field research permits were granted by the Centre de Surveillance de la Biodiversité in Kisangani of University of Kisangani at the University of Kisangani, with provincial and local authorities approving all mission orders. Samples exported to the Royal Belgian Institute of Natural Sciences and the University of Antwerp (Belgium) prior to the implementation of the Nagoya Protocol were covered by Centre de Surveillance de la Biodiversité material transfer agreements, while those exported after 2018 were authorized by a Nagoya export permit issued by the Congolese Ministry of the Environment and Sustainable Development (MEDD) (Permit numbers 003/ANCCB-RDC/SG-EDD/BTB/08/2019, 018/ANCCB-RDC/SG-EDD/BTB/07/2022, 021/SG-EDD/BTB/ANCCB-RDC/08/2023, 011/ANCCB-RDC/SG-EDD/BTB/2023). No CITES permits were required. Permission to import the samples into Belgium was granted by the Federal Agency for the Safety of the Food Chain (FASFC), permit numbers 2022DBP504, 2023DBP370, 2023DBP371, 2024DBP424.

#### 2.1.2. EXTERNAL MORPHOMETRY AND CRANIOMETRY

Squirrels were first identified *in situ* to genus or species level using **Kingdon & Hoffmann (2013**). We examined captured squirrels on the basis of external morphological criteria, *i.e*. coat, body length, tail length, hind foot length, and ear length. Sex and reproductive stages were determined by external observation, including the presence or absence of nipples (small or swollen) and the appearance of the vagina (closed or open) in females or the presence of testes (abdominal or scrotal) in males. Length was measured to the nearest millimeter on fresh specimens using a Mitutoyo caliper to an accuracy of 0.05 mm. Weight was measured using a spring balance (Pesola) and expressed in grams.

Only the skulls of adult squirrels were extracted and measured (**Supplementary Table S1**). A total of 23 measurements were taken on each skull (**Verheyen & Bracke, 1966; Van Der Straeten & Dieterlen, 1987; Verheyen et al., 1996**). The used measurements are listed with its corresponding number, abbreviation, and morphological character (**Supplementary Table S2).**

We also measured 29 skulls from the collections of the Royal Museum for Central Africa-Tervuren to link a craniometrically defined Operational Taxonomic Unit (OTU) to cytochrome b sequences previously published in GenBank (**Falendysz at al., 2017**).

#### 2.1.3. ANALYSIS OF MORPHOMETRIC AND CRANIOMETRIC DATA

The approach consisted of several steps. First, OTUs were designated based on preliminary Molecular Taxonomic Units (MOTUs) identified using cytochrome b data. Next, we tested the hypothesis that variation in craniometrics and external morphological data aligns with the MOTUs through multivariate analysis using R version 4.2.3 and Statistica version 12 (StatSoft Inc).

To evaluate sexual dimorphism within OTUs, all measurements were analyzed. However, as no significant sexual dimorphism was detected (**Supplementary Tables S3 and S4)**, subsequent analyses were conducted using specimens of both sexes. Basic statistics, including the number of observations, mean, minimum, maximum, standard deviation, and coefficient of variation, were calculated for all OTUs (**Supplementary Tables S5 and S6**)

A forward discriminant analysis was then performed using all 23 variables from the complete craniometric dataset, clearly highlighting the sequenced specimens. A second discriminant analysis focused on the OTUs *Funisciurus anerythrus* Left Bank (FaLB), *Funisciurus anerythrus* Right Bank (FaRB), and *Funisciurus cf congicus* (Fc) near Kikwit, using a partial craniometric dataset. For the Kikwit specimens, where some measurements were unavailable due to skull damage, a mean substitution method was applied to estimate missing values (Statistica) (**Supplementary Figure S4).** Finally, a third discriminant analysis incorporated all available external measurements for all OTUs, again clearly highlighting the sequenced specimens.

Not all specimens were intact, the tail and/or ear having been cut off and the skull bones fractured, the skull not extracted or not recovered, which explains the variations in the number of individuals (N) used in each type of analysis. The subset of tissues bearing a DNA barcode was selected at random on the basis of the geographical origin of the capture and the availability of morphometric measurements. We also measured 41 squirrel skulls collected in Kikwit, Kinshasa and Kongo Central (DRC) and stored at the Royal Museum for Central Africa, and analyzed their tissue samples stored at the Royal Belgian Institute of Natural Sciences.

### 2.2. MOLECULAR DATA ACQUISITION

We extracted DNA of squirrel tissue samples (tongue or liver or kidney or lung) using the Nucleospin Tissue mini kit according to the manufacturer’s instructions (Machery-Nagel, Düren, Germany), with an overnight pre-lysis incubation instead of 1-3h, and a one-minute elution at 70°C instead of RT. We then amplified a mitochondrial gene fragment using universal cytochrome b primers L14723 and H15915 (**Montgelard et al., 2002**) with Promega MasterMix PCR following the manufacturer’s instructions (Promega, Madison, USA). The initial denaturation was at 94°C for 3 min, followed by 35 cycles at 94°C for 1 min, 52°C for 1 min and 72°C for 1 min. Final extension was carried out at 72°C for 7 minutes.

We then purified the amplified products using ThermoFisher Exo-SAP IT PCR product clean-up reagent (ThermoFisher Scientific, Santa Clara, USA) and sent them to Macrogen (Amsterdam, The Netherlands) for sequencing. Finally, we manually cleaned the sequences based on the chromatograms and aligned them using the MAFFT algorithm within Geneious 2019.2.3. This alignment included in-house generated *Funisciurus* sequences, as well as two *Paraxerus* sequences serving as an outgroup. This alignment also included *Funisciurus sp* sequences downloaded from GenBank, on the condition that they did not contain stop-codons and clustered with our *Funisciurus* sequences rather than with the outgroup (accession numbers: MF694315, MF694313, MF694317, MF694320, MN597594 and MN597668), yielding a total of 143 sequences.

### 2.3. SPECIES DELIMITATION USING GENETIC DATA

We first inferred an approximately-maximum-likelihood phylogenetic tree using MAFFT-aligned mitochondrial sequences and the FastTree plugin in Geneious 2019.2.3, using default settings and rooting on the *Paraxerus* outgroup (**Price et al., 2009**). Briefly, the FastTree plugin worked as follows: a rough tree topology was first obtained through a neighbour-joining approach; tree length was then reduced using nearest-neighbour interchanges and subtree-prune-regraft-moves. Tree topology and branch length were improved using maximum-likelihood rearrangements, where each site was assigned to one of 20 geometrically spaced categories (0.05-20) using a Bayesian approach (gamma prior), representing each site’s most likely evolutionary category. Finally, to estimate the reliability of each split in the tree, the Shimodaira-Hasegawa test was used on three alternate topologies (NNIs) around the focal split. The resampling size was 1,000 (**Price et al., 2009**).

To further explore *Funisciurus* taxon delineation within our dataset, we used species delimitation algorithms that generate species partitions based either on distance (ASAP, Assemble Species by Autonomous Partitioning) or phylograms (bPTP, Bayesian Poisson Tree Processes), to offer complementary perspectives (**DeSalle, 2024**). First, we employed the ASAP method on our MAFFT alignment, with the following adjustments to meet ASAP requirements: five GenBank sequences (MF694315, MF694313, MF694317, MF694318, and MF694320) were removed to ensure overlap across sequences. ASAP is a frugal, computationally efficient method, which is not based on the reconstruction of phylogenies (**Puillandre et al., 2021**). Rather, this distance-based approach identifies species partitions by detecting barcode gaps—the difference between small intraspecific and large interspecific sequence distances (**Puillandre et al., 2021**).

ASAP uses an ascending hierarchical clustering algorithm, which sequentially merges sequences, based on pairwise distances, until all sequences are grouped within the same partition (**Additional file 1**). For each partition, a *p*-value (likelihood of panmixia, *i.e*., a low *p*-value signifies that merging the proposed subsets would lead to intra-subset genetic diversity that is higher than expected within a single panmictic species), and W-score (barcode gap width metric, calculated using the threshold distance, *i.e*., the midpoint between the distance that triggered the merging of subsets and the distance before merging) are computed (**Additional file 1**). A larger W-score reflects a larger barcoding gap, *i.e*., species are well separated. After clustering, the *p*-value and W-score of each partition are sorted by ascending and descending order, respectively, to compute the ASAP-score (averaged ranks of *p*-value and W-score for each partition). Consequently, smaller ASAP-scores reflect better partitions. Its main advantage over other automated barcode gap discovery approaches is that the maximal intraspecific genetic distance does not need to be specified by the user *a priori*, which makes it suitable for under-researched taxa such as African rope squirrels. We used the most recent ASAP tool, (**Puillandre et al., 2021**), and compared simple p-distances, Jukes-Cantor (JC69) and Kimura 2-parameter (K80) substitution models to investigate whether substitution models impacted the proposed species delimitation. Only results from JC69 are presented here.

For additional robustness, we applied the bPTP (**Zhang et al., 2013**) method. This method uses phylogenetic tree branching patterns to propose species partitions (*i.e*., evaluating whether the number of substitutions follow a Poisson distribution within *versus* between species) and provides Bayesian support values for these partitions.To meet bPTP requirements (**Zhang et al., 2013**), we ran an IQ-TREE analysis (using default settings from the IQ-TREE tool, (**Nguyen et al., 2015; Hoang, et al., 2017; Kalyaanamoorthy et al., 2017; Minh et al., 2020**) on the MAFFT alignment used in the ASAP method, with the following adjustments: only *Funisciurus* sequences, non-redundant haplotypes, and fully overlapping sequences were included, for a total of 47 sequences. The resulting phylogenetic tree was used as an input file for the bPTP analysis (**Additional file 2**).

Finally, we compared the species partitions proposed by ASAP and bPTP with the topology of the FastTree and the results of discriminant analyses based on morphometric data.

## 3. RESULTS

### 3.1. TRAPPING EFFORT

During a total of 99,035 trapping nights, the trapping effort resulted in the capture of 429 squirrels belonging to the genus *Funisciurus*, (success rate of 0.053%). Of these squirrels, 301 were caught on the left bank of the Congo River and 128 on the right bank. No reliable classification could be established based on external morphometric characteristics alone.

### 3.2. SPECIES PARTITIONING

Overall, approaches based on mitochondrial sequences yielded between four and six MOTUs, when excluding singletons and outgroups. The following MOTUs could be identified based on ASAP and bPTP results: FaLB, FaRB, Fc, FbLB and Fx. Based on ASAP results only, FbLB was further split into FbLB1 and FbLB2. The general species partitioning obtained using mitochondrial data was supported by craniometric data, with the exception of Fx, for which an insufficient number of cranial measures could be taken.

#### 3.2.1. SPECIES PARTITIONING USING ASAP

Using the best available species partition, with the lowest ASAP score (ASAP score = 1.50, **Table 1**, hereafter ASAP1), seven MOTUs of *Funisciurus* were distinguished, in addition to two outgroup MOTUs. However, one MOTU is represented by a single sequence (YK156), giving a total of six well-supported MOTUs consisting of two or more specimens each (**Figure 1, Additional file 1**). Using the second-best species partition, with the second-lowest ASAP score, (ASAP score = 3, **Table 1**), eight MOTUs of *Funisciurus* were distinguished, in addition to two outgroup MOTUs. However, two MOTUs are represented by a single sequence each (YK156 and NDU803), resulting in the same six well-supported MOTUs identified in the first partition (**Additional file 1**). Using the smallest *p-*value to determine the best available species partition (ASAP score = 5.50, p = 0.00022, **Table 1**, hereafter ASAP2), five *Funisciurus* MOTUs can be distinguished, in addition to two outgroup taxa. These include, again, a MOTU represented by a single sequence (YK156), yielding a total of four well supported MOTUs consisting of two or more specimens each. In contrast to previous partitions, FaLB, FaRB and Fc are merged within a single MOTU (**Figure 1, Additional file 1**).

**Figure 1.**
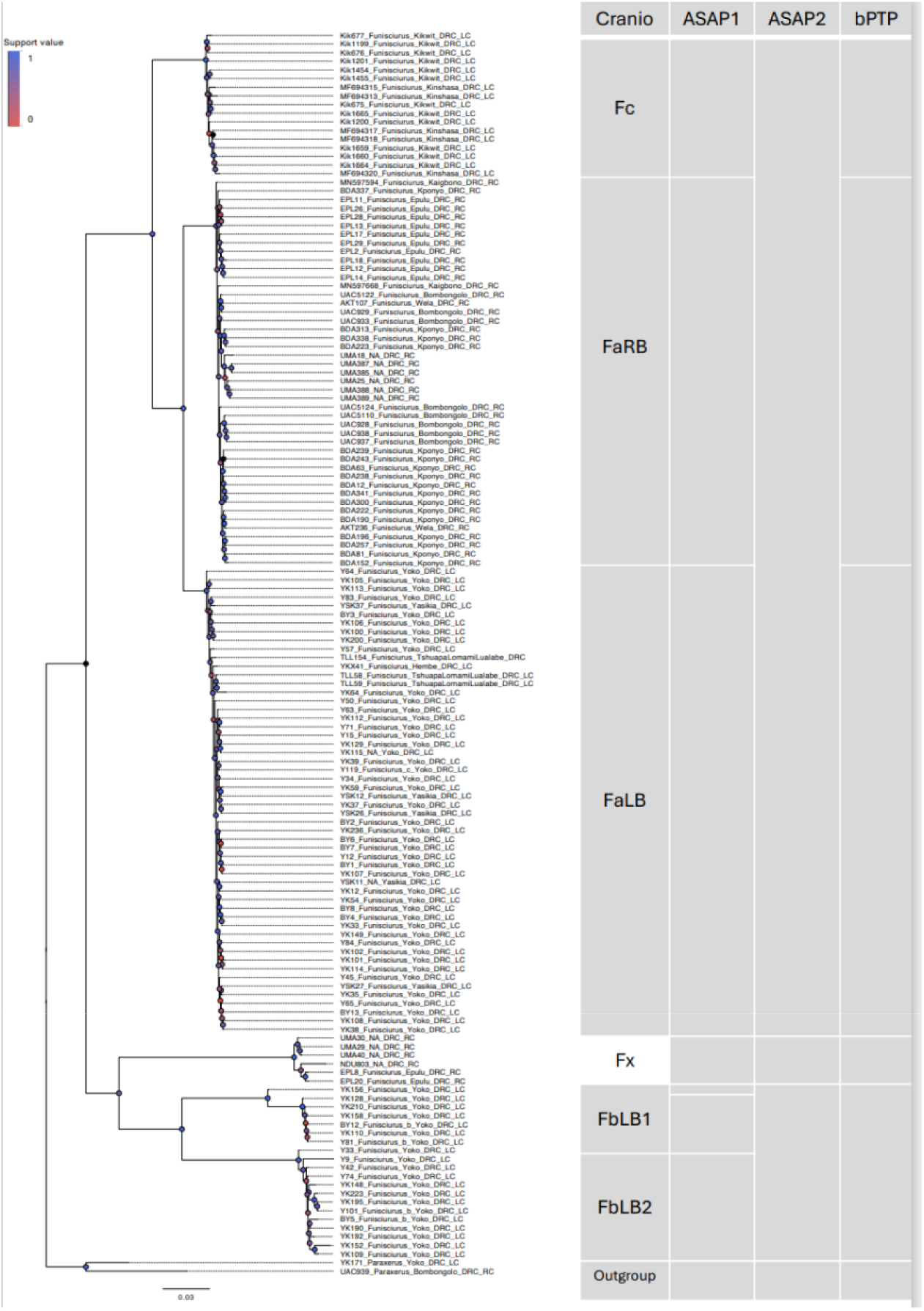
Proposed rope squirrel Operational Taxonomic Units using a combination of 725 methods. Above the branches, methods used to propose species partitions. Cranio: Craniometry; ASAP1: Assemble Species by Autonomous Partitioning (ASAP), lowest ASAP score; ASAP2: *idem*, lowest p-value; bPTP: bayesian Poisson Tree Processes; RB: right bank of the Congo River; LB: left bank of the Congo River. Fx: previously undescribed *Funisciurus* species; Fb: *Funisciurus cf bayonii*; Fc: *Funisciurus cf congicus*; Fa: *Funisciurus anerythrus*. The tree was made using FastTree on MAFFT-assembled cytochrome b sequences.

**Table 1.** Species partitions ranked in ascending order of ASAP scores. The *p*-value reflects the likelihood that Partition is different from Partitionn-1 and is ranked in ascending order. The threshold distance is used to compute the W-score and represents the mid-point between the distance that triggered the merging of subsets into Partition and the previous distance in the list. The W-score is the relative barcode gap width associated with Partitionn-1 and is ranked in descending order. The ASAP-score is the average of the ranks of the *p*-value and of the W-score. A smaller ASAP-score thus signifies a better partition (**Puillandre et al., 2021**). Read: Considering an ASAP-score of 2.00 and a p-value of 0.0177, 9 subsets are identifiable in the dataset, including two outgroups.

| Number of subsets | ASAP-score | P-val (rank) | W (rank) | Threshold dist. |
| --- | --- | --- | --- | --- |
| *9 | 2.00 | 1.77e-02 (3) | 7.83e-04 (1) | 0.020430 |
| 5 | 3.50 | 1.96e-01 | 2.03e-04 (3) | 0.118381 |
| *8 | 4.50 | 3.46e-03 (02) | 4.68e-05 (7) | 0.029208 |
| 6 | 5.00 | 3.80e-04 (1) | 2.56e-05 (9) | 0.075144 |
| *12 | 7.00 | 5.45e-01 (8) | 4.72e-05 (6) | 0.0008821 |
| 8 | 11.50 | 6.87e-01 (15) | 3.11e-05 (8) | 0.052757 |
| *10 | 12.00 | 8.40e-01 (22) | 5.20e-04 (2) | 0.015671 |
| 59 | 14.00 | 5.73e-01 (10) | 6.22e-06 (18) | 0.000899 |
| 60 | 15.50 | 6.79e-01 (14) | 6.22e-06 (17) | 0.000450 |
| 2 | 15.50 | 8.14e-01 (20) | 1.15e-05 (11) | 0.180112 |

#### 3.2.2. SPECIES PARTITIONING USING bPTP

Using the bPTP method applied to our IQ-TREE, six MOTUs of *Funisciurus* were identified with support values ranging from 0.697-0.984. However, one of them consists of a single sequence (YK110, support value: 1.000). Thus, bPTP identifies five well supported MOTUs represented by two or more specimens: Fx (0.984), Fc (0.900), FaLB (0.820), FaRB (0.817), and FbLB (0.697) (Figure 1, Additional file 2).

#### 3.2.3. SPECIES PARTITIONING BASED ON EXTERNAL MORPHOMETRIC CHARACTERISTICS AND SEXUAL DIMORPHISM

Comparing Student’s t-tests with the craniometric (**Supplementary Table S3**) and external (**Supplementary Table S4**) measurements of males and females of each species did not reveal any sexual dimorphism. In all subsequent analyses, sexes were therefore lumped.

The discriminant analysis carried out on the external measurements (Weight, Length of hind foot, Length of ear, Length of tail, Length of body) revealed low discriminating power (Wilk’s Lambda = 0.229) (**Supplementary Figure S3**). There is no distinction between *F. anerythrus* from the right bank and from the left bank. *F.cf bayonii* (Fb) is clearly separated from *F. anerythrus* along root 1. No external measurements are available for *F.cf congicus*.

#### 3.2.4. SPECIES PARTITIONING BASED ON CRANIAL MORPHOMETRIC CHARACTERISTICS

Unlike external measurements, craniometric measurements of adult rodents offer greater stability and are among the most effective methods for identifying rodent species. For this reason, 220 adult squirrel skulls were measured and 23 measurements were taken on each skull (**Supplementary Table S2**).

A forward discriminant analysis, (**Supplementary Figure S5**) omitting measures M8 (width of the interorbital narrowing), M9 (width of the zygomatic arch) and M19 (maximum width of the cranial cavity) to maximize the number of observations for *F. cf congicus* (Fc), measures (M1: maximum length of the skull; M2: condilo-basal length; M3: henselion-basion length; M4: henselion-palation length; M5: length of the cleft palate; M6: length of the diastema; M7: distance between the anterior edge of the alveolus of M1 and the cutting edge of the upper incisor; M10: minimum width of the palate; M11: length of the upper molar row; M12 : external width of upper molar row at M1; M13: maximum molar width; M14: minimum width of zygomatic plate; M15: maximum width of nasals; M16: maximum length of nasals; M17: length of lower molar row; M18: length of the tympanic bulla; M20: depth of the upper incisors; M21: height of the rostrum at the anterior edge of the alveolus of M1; M22: maximum width of the rostrum; M23: distance between the extreme points of the coronoid and angular processes) reveals a good discrimination. *F. anerythrus* (FaRB and FaLB) and *F cf bayonii* (FbLb) are clearly separated along root 1.

*F.cf congicus* (only six sequenced specimens with mesured skulls, see **figure 1**) is clearly different from both other species. *F. anerythrus* Right Bank (FaRB) and Left Bank (FaLB) are well differentiated along root 2. The number of sequenced specimens of the two genotypes of *F.cf bayonii* (FbLB1 and FbLB2) are too small to separate them in this analysis (**Supplementary Figure S5**).

A discriminant analysis, performed on the OTUs of *F. anerythrus* from the right bank and left bank, as well as FbLB1 and FbLB2 using a craniometric dataset limited to the skulls of specimens with matching genetic sequences, while omitting the often missing M6, M7 and M20 measurements (**Supplementary Figure S3**), reveals subtle differences between *F. cf bayonii* sp1 (FbLB1) and *F. cf bayonii* sp2 (FbLB2). However, the limited availability of specimens with both sequence data and craniometric measurements has hampered the differentiation of these potentially distinct OTUs. The same figure also reveals clear differences between *F. anerythrus* from the two banks of the Congo River (FaRB and FaLB) and no difference between *F. anerythrus* from the right bank (FaRB and Fx in **Supplementary Figure S6**).

#### 3.2.5 OVERALL SPECIES PARTITIONING

Our most conservative results, found using a barcoding gap analysis with partition scoring where we retained the partition with the lowest *p*-value (ASAP2), suggest the presence of four MOTUs consisting of at least two specimens each. Here, *F. anerythrus* and *F. cf congicus* form one MOTU, whereas *F. cf bayonii*, and a previously undescribed *Funisciurus* species (here, Fx), each form another MOTU.

Our second most conservative result, using a tree-based algorithm inferring species delimitation based on the distribution of substitutions between and within species (bPTP), suggests the presence of five MOTUs. These are *F. cf bayonii* (FbLB), the undescribed *Funisciurus* species (Fx), *F. anerythrus* individuals found on the left bank of the Congo River (FaLB), *F. anerythrus* individuals found on the right bank of the Congo River (FaRB), and *F. congicus* (Fc).

The discriminant analysis conducted on craniometric data suggests the presence of five OTUs, however differently distributed: *F.cf bayonii* individuals form two weakly supported OTUs (FbLB1 and FbLB2), while Fc, FaLB and FaRB each form a well-supported OTU. While Fx does not appear to be different from FaRB, the limited number of measurable skulls for Fx prevents any conclusions about craniometric differences between Fx and FaRB.

Last, our least conservative results, based on a barcoding gap analysis with partition scoring where we retained the partition with the lowest ASAP score (ASAP1), suggest the presence of six well-supported MOTUs of *Funisciurus* represented by at least two individuals each. These MOTUs are Fc, FaLB, FaRB, Fx, FbLB1, and FbLB2, and are in line with the FastTree topology (**Figure 1**).

### 3.3. DISTRIBUTION AND HYDROLOGICAL BARRIERS

The six main well-supported taxa occupy distribution areas separated by hydrological barriers (**Figure 2**). The previously undescribed *Funisciurus* species (Fx) originates exclusively from the right bank of the Congo River (Figure 2a) and the *F. cf bayonii* species (FbLB1 and FbLB2) (Figure 2a) are sympatric and distributed exclusively on the left bank of the Congo River. In addition, FaRB is distributed exclusively on the right bank of the Congo River, and FaLB is distributed exclusively on the left bank of the Congo River. *F. cf congicus* (Fc) is distributed exclusively on the left bank of the Kasaï River (Figure 2b).

**Figure 2.**
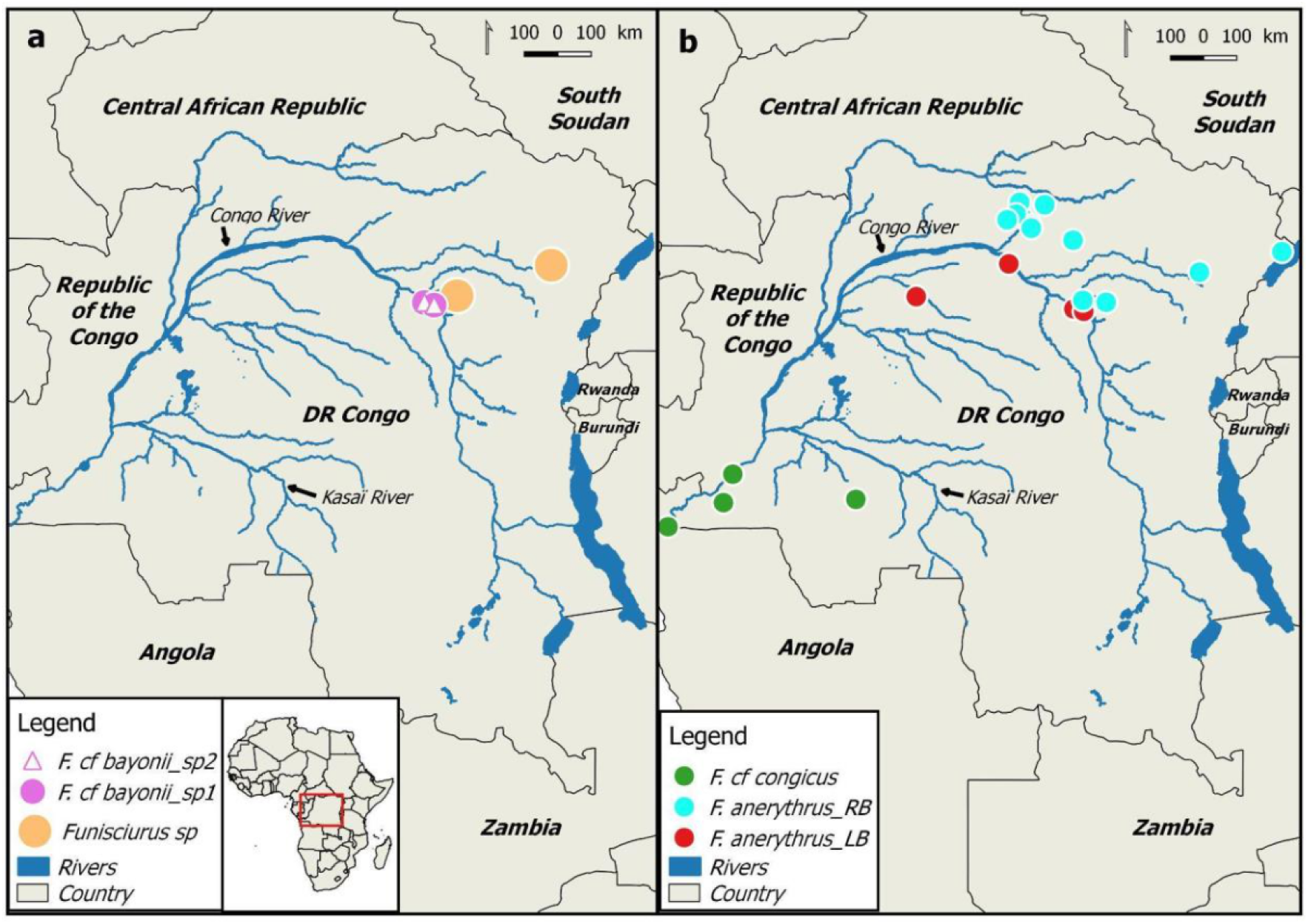
Maps showing the distribution of the species and subspecies identified in relation to the Congo River and its tributaries, (**a**) *Funisciurus sp* from the right bank of the Congo River and the sympatric *F. cf bayonii sp1* and *F. cf bayonii sp2* on the left bank of the Congo River; (**b**) *F. anerythrus_RB* on the right bank, *F. anerythrus_LB* on the left bank of the Congo River, and *F. cf congicus* on the left bank of the Kasaï River.

## 4. DISCUSSION

This study had two main objectives: first, to ascertain the number of *Funisciurus* OTUs present in the Congo Basin; and second, to investigate the role of the Congo River as a geographical barrier between *Funisciurus* taxa. For this purpose, we integrated morphometric, molecular and zoogeographic data from the most extensive *Funisciurus* species dataset to date. We first identified a discrepancy between the previously described species of *Funisciurus* in our study area, and the individuals present in our dataset (**Table 2**, **Figure 1**). We then found that both the Congo River and the Kasaï River delineate the distribution of *Funisciurus* OTUs (**Figure 2**). This highlights the importance of hydrological barriers in the speciation processes of small arboreal mammals in the Congo Basin and calls for an update in the taxonomy and IUCN distribution maps of rope squirrels.

Previously, five rope squirrel species had been described in our study area: *F. anerythrus* (Thomas, 1890), *F. bayonii* (Bocage, 1890), *F. congicus* (Kuhl, 1820), *F. pyrropus* (F. Cuvier, 1833), and the mountain-dwelling *F. carruthersi* Thomas, 1906 (**Kingdon & Hoffmann, 2013; Thorington et al., 2012; Gerrie et al., 2016**).

To minimise the risk of OTU over-segregation or fusion in our analysis (**Sloan et al., 2017; Nesi et al., 2011**), our final OTU partitioning is based on both craniometric and molecular OTU delineation, as well as a zoogeographic analysis of their distribution. To do so, we have integrated multiple OTU delineation methods (**Demos et al., 2013**), *i.e.,* a discriminant analysis of craniometric data, two barcoding gap analyses with partition scoring (ASAP1 and ASAP2), and a tree-based algorithm inferring species delimitation based on the distribution of substitutions between and within species (bPTP), with OTU distribution maps, where well-supported OTUs distributed in the same hydrological basin indicate the presence of non-interbreeding, sympatric species; while allopatric OTUs could theoretically interbreed, but are prevented to do so by hydrological barriers. Thus, using our most conservative results (ASAP2) and our updated OTU distribution, we suggest that there exist four *Funisciurus* species in the Congo Basin, namely *F. anerythrus* (composed of three allopatric subspecies: Fc, FaLB, and FaRB); two sympatric, but genetically and craniometrically different, thus non-interbreeding *F. cf bayonii* species (FbLB1, FbLB2); and the previously undescribed *Funisciurus* species (Fx).

Several key points regarding the taxonomy of *F. anerythrus*, *F. bayonii*, and an undescribed *Funisciurus* species warrant further discussion. According to previous studies, *F. anerythrus*—originally described from Buguera, Uganda—has a distribution spanning from Uganda to Angola and Togo (**Thorington & Hoffmann, 2005**), covering all central Congolese forest blocks (**Colin, 1991**). **Kingdon & Hoffmann (2013)** identified four subspecies based on geographical and morphological variations. The southernmost, *F. a. anerythrus*, occurs from western Uganda to southern DRC, reaching Mount Kabobo and northern Angola. It has a reddish-brown dorsal coat and a yellow to red ventral coat. The northernmost, *F. a. bandarum*, ranges from the Central African Republic to southern Chad, featuring a buff-tinged dorsal coat and a pale grey to orange ventral side. Either of these may correspond to our OTU FaRB, but limited skull and DNA data prevent verification. *F. a. mystax*, found in southern Cameroon and Gabon, has a reddish-brown dorsal and orange ventral coat, while *F. a. raptorum*, endemic to Benin and Nigeria, displays a grey dorsal and whitish ventral surface. Due to morphological similarities and limited overlap in our dataset, these findings reinforce the need for a thorough taxonomic reassessment of the *F. anerythrus* complex.

Conversely, previous studies indicate that *F. bayonii* is found mostly in western DRC, with no described subspecies (**Thorington et al., 2012; Kingdon & Hoffmann, 2013; Gerrie et al., 2016**). This species has only been documented outside the Central African forest block and was previously considered a species of wooded savannah mosaics (**Thorington et al. 2012**). However, our results show that two species morphologically similar to *F. bayonii* are present in Yoko and Yasikia, located near the left bank of the Congo River, in evergreen tropical forests. This result corroborates the need for further taxonomic revision to clarify the status of these species, including their ecological niches.

Last, we report the existence of a species that does not cluster with either *F. anerythrus* nor *F.* cf *bayonii*, and which is distributed north of the Congo River (e.g., UMA30). Its range does not overlap with the known range of the mountain-restricted *F. carruthersi* Thomas, 1906, and it does not display the typical red-leggedness of *F. pyrropus* (F. Cuvier, 1833). Although its appearance resembles that of individuals of *F. anerythrus* found on the right bank of the Congo River (FaRB), all OTU delineation techniques based on genetic data suggest that it is a yet undescribed species of rope squirrel.

Beyond taxonomic considerations, we highlight that the distribution ranges of these squirrels are different, and much smaller than previously thought, rendering their populations more vulnerable to environmental change and habitat degradation (**Steiner M. & Huettmann F., 2023**). Current IUCN distribution maps indicate that *F. congicus*’ northeastern expansion is limited by the Congo River, while *F. bayonii* is confined southwest of the Kasaï River. (data retrieved from IUCN Red List (https://www.iucnredlist.org)**; Gerrie et al., 2016**). However, our findings contradict this. We observed that *F. congicus* (Fc) is restricted southwest of the Kasaï River, with no specimens found between the Kasaï and Congo Rivers. Additionally, unlike the IUCN map for *F. bayonii*, we found that the sympatric FbLB1 and FbLB2 OTUs are exclusively limited to this interfluvial region, where the Kasaï River restricts southwestern expansion, and the Congo River blocks northeastern expansion. Similarly, contrary to IUCN data for *F. anerythrus*, the Congo River separates FaLB (southwest) and FaRB (northeast). The undescribed Fx is found only northeast of the Congo River (**Figure 2**). These findings highlight the role of hydrological barriers in driving allopatric speciation in arboreal Sciuridae, consistent with research on primates (**Anthony et al., 2007; Harcourt & Wood, 2012; Fonteyn et al., 2023**) and small mammals rodentia (**Katuala et al., 2008; Kennis et al., 2011**). They further highlight that the distribution range of each *Funisciurus* OTU is smaller than previously thought (**Supplementary Figure S7)**. Consequently, habitat degradation or environmental change may have unexpectedly severe impacts on *Funisciuru*s populations (**Supplementary Figure S8)**, of which regular monitoring is necessary. We therefore advocate for updated IUCN distribution maps and regular monitoring of the *Funisciurus* OTUs identified above, to ensure that conservation measures can be taken where appropriate.Regular monitoring will also support research on the potential epidemiological importance of *Funisciurus* species.

## CONCLUSION

Our morphometric and molecular analyses enabled us to identify six main rope squirrel OTUs in the Congo Basin, namely *F. cf congicus*, *F. anerythrus*_LB, *F. anerythrus*_RB, *Funisciurus sp*, *F. cf bayonii*_LB1, and *F. cf bayonii*_LB2. Further integration of these approaches with zoogeographic analyses suggests that Fc (*F. cf congicus* (Kuhl, 1820)), *F. anerythrus*_LB and *F. anerythrus*_RB are subspecies of *F. anerythrus* (Thomas, 1890), while the two sympatric and non-interbreeding *F. cf bayonii*_LB1, and *F. cf bayonii*_LB2 each are a separate species, possibly forming a complex with *F. bayonii* (Bocage, 1890). Last, we report on a previously undescribed species of *Funisciurus* (Fx). We further highlight the role of the Congo and Kasaï as hydrological barriers, delineating the distribution of our OTUs, with *F. cf congicus* found exclusively southwest of the Kasaï, *F. anerythrus*_LB, *F. cf bayonii_*LB1 and *F. cf bayonii*_LB2 found between the Kasaï and Congo, and *F. anerythrus*_RB and Fx found northeast of the Congo River. These results call for an update of IUCN maps, and for regular monitoring of *Funisciurus* populations, which may be more vulnerable to environmental change and habitat degradation than previously thought. Last, we highlight that further taxonomic and ecological studies are needed to describe each OTU and ascertain its ecological niche, which will support research on zoonotic pathogen spillover risks associated with each OTU. In this perspective, we also recommend a complete review of the taxonomic status of other tree squirrels, in particular the genera *Paraxerus, Heliosciurus* and *Protoxerus*.

## ACKNOWLEDGEMENTS

This research was funded by the Flemish Interuniversity Council South-Initiative (VLIR-SI) (AL, EV), the Flemish Interuniversity Council (VLIR-CUI) (GG, EV), the Research Foundation—Flanders (G051322N), and a FED-tWIN scholarship funded by the Belgian federal government (Prf-2019-prf004_OMEga) (SG). The CEBioS program (Royal Belgian Institute of Natural Sciences) provided financial support for the CSB team’s fieldwork (PB, CM, GG) and research stays in Belgium (PB).

We thank the Faculty of Science and the Centre de Surveillance de la Biodiversité of University of Kisangani for their supervision. We also thank Emmanuel Gilissen and Marthys Rotonda for allowing us access to Squirrel specimens of the collection of MRAC-Tervuren.

## Full e-mail addresses of authors

**Claude Mande,**

**Anne Laudisoit,** **ou**

**Nicaise Amundala**

**Nicolas Laurent**

**Ganiyat Temidayo Saliu,**

**Joachim Mariën**

**Sophie Gryseels**

**Guy-crispin Gembu**

**Jan Hulselmans**

**Erik Verheyen**

## Contributions of the authors

PB, EV and JH conceptualized the study and designed the methodology. PB collected the data and identified the species in the field. LF, PB, GTS and NL generated molecular data. PB, JH, and EV analyzed the morphometric data. PB produced the map. LF, SG and EV analysed the molecular data. PB, LF, JH and EV wrote the original version. PB, LF, CM, AL, NA, NL, GTS, JN, JM, SG, GG, JH and EV reviewed and edited the document. All authors approved the final manuscript.

## Data availability

The generated and/or analyzed datasets can be made available for research purposes upon request to the corresponding authors. The sequences from this study will be submitted to GenBank upon acceptance.

