## Supplemental data for "Reassessing the diversity and distribution of African rope squirrels (*Funisciurus* Trouessart, 1880)"

**Supplementary tables**

**Table S1.** List of specimens examined morphologically and/or genetically (1 = Specimen (skull/tissue) examined/analysed; 0 = Specimen (skull/tissue) not examined/analysed)

| **Province** | **Locality** | **Geographic coordinates** | **Label** | **Morphometric** | | **Molecular** | **Observation** |
|---|---|---|---|---|---|---|---|
| | | | | **Coat** | **Skull** | **Cyt b** | |
| Bas Uele | Aboso | 2.89655°N; 23.11306°E | LIK759 | 1 | 0 | 0 | |
| Bas Uele | Aboso | 2.89655°N; 23.11306°E | LIK782 | 1 | 0 | 0 | |
| Bas Uele | Ahupa | 2.65748°N; 23.79874°E | UAC902 | 1 | 0 | 0 | |
| Bas Uele | Bogala | 2.95623°N; 23.53877°E | UAC918 | 1 | 0 | 0 | |
| Bas Uele | Bogala | 2.95623°N; 23.53877°E | UAC923 | 1 | 0 | 0 | |
| Bas Uele | Bombongolo | 3.05659°N; 23.36565°E | UAC925 | 1 | 0 | 0 | |
| Bas Uele | Bombongolo | 3.05659°N; 23.36565°E | UAC926 | 1 | 0 | 0 | |
| Bas Uele | Bombongolo | 3.05659°N; 23.36565°E | UAC929 | 1 | 0 | 1 | |
| Bas Uele | Bombongolo | 3.05659°N; 23.36565°E | UAC936 | 1 | 0 | 0 | |
| Bas Uele | Bombongolo | 3.05659°N; 23.36565°E | UAC 5110 | 1 | 0 | 1 | |
| Bas Uele | Bombongolo | 3.05659°N; 23.36565°E | UAC 5122 | 1 | 0 | 1 | |
| Bas Uele | Bombongolo | 3.05659°N; 23.36565°E | UAC 5124 | 1 | 0 | 1 | |
| Tshopo | Mbiye | 0.45013889°N; 25.29930556°E | MBIYE 523 | 1 | 0 | 0 | |
| Tshopo | Mbiye | 0.45013889°N; 25.29930556°E | MBIYE 524 | 1 | 0 | 0 | |
| Tshopo | Mbiye | 0.45013889°N; 25.29930556°E | MBIYE 527 | 1 | 0 | 0 | |
| Tshuapa | Inkanamongo | 0.71375°N; 20.530778°E | EBO62 | 1 | 0 | 0 | |
| Tshuapa | Inkanamongo | 0.71375°N; 20.530778°E | EBO98 | 1 | 0 | 0 | |
| Tshuapa | Inkanamongo | 0.71375°N; 20.530778°E | EBO314 | 1 | 0 | 0 | |
| Tshuapa | Inkanamongo | 0.71375°N; 20.530778°E | EBO315 | 1 | 0 | 0 | |
| Bas Uele | Kaigbono | 3.39021°N; 23.46676°E | LIK 194 | 1 | 0 | 0 | |
| Bas Uele | Kaigbono | 3.39021°N; 23.46676°E | LIK 280 | 1 | 0 | 0 | |
| Bas Uele | Kaigbono | 3.39021°N; 23.46676°E | LIK 926 | 0 | 0 | 0 | |

| | | | | | | | |
|---|---|---|---|---|---|---|---|
| Bas Uele | Kaigbono | 3.39021°N; 23.46676°E | LIK 930 | 0 | 0 | 0 | |
| Bas Uele | Kaigbono | 3.39021°N; 23.46676°E | LIK 931 | 0 | 0 | 0 | |
| Kwilu | Kikwit | -5.038308°N; 18.817844°E | KIK 1199 | 0 | 0 | 1 | MRAC |
| Kwilu | Kikwit | -5.038308°N; 18.817844°E | KIK 1200 | 0 | 0 | 1 | MRAC |
| Kwilu | Kikwit | -5.038308°N; 18.817844°E | KIK 1201 | 0 | 0 | 1 | MRAC |
| Kwilu | Kikwit | -5.038308°N; 18.817844°E | KIK 1454 | 0 | 0 | 1 | MRAC |
| Kwilu | Kikwit | -5.038308°N; 18.817844°E | KIK 1455 | 0 | 0 | 1 | MRAC |
| Kwilu | Kikwit | -5.038308°N; 18.817844°E | KIK 1659 | 0 | 0 | 1 | MRAC |
| Kwilu | Kikwit | -5.038308°N; 18.817844°E | KIK 1660 | 0 | 0 | 1 | MRAC |
| Kwilu | Kikwit | -5.038308°N; 18.817844°E | KIK 1664 | 0 | 0 | 1 | MRAC |
| Kwilu | Kikwit | -5.038308°N; 18.817844°E | KIK 1665 | 0 | 0 | 1 | MRAC |
| Kwilu | Kikwit | -5.038308°N; 18.817844°E | KIK 1669 | 0 | 0 | 0 | MRAC |
| Kwilu | Kikwit | -5.038308°N; 18.817844°E | KIK 1671 | 0 | 0 | 0 | MRAC |
| Kwilu | Kikwit | -5.038308°N; 18.817844°E | KIK 676 | 0 | 0 | 1 | MRAC |
| Kwilu | Kikwit | -5.038308°N; 18.817844°E | KIK 677 | 0 | 0 | 1 | MRAC |
| Bas Uele | Kponyo | 3.3243°N; 24.17641°E | BLI02 | 1 | 0 | 0 | |
| Bas Uele | Kponyo | 3.3243°N; 24.17641°E | BDA112 | 1 | 0 | 0 | |
| Bas Uele | Kponyo | 3.3243°N; 24.17641°E | BDA152 | 1 | 0 | 1 | |
| Bas Uele | Kponyo | 3.3243°N; 24.17641°E | BDA196 | 1 | 0 | 1 | |
| Bas Uele | Kponyo | 3.3243°N; 24.17641°E | BDA238 | 1 | 0 | 1 | |
| Bas Uele | Kponyo | 3.3243°N; 24.17641°E | BDA257 | 1 | 0 | 1 | |
| Bas Uele | Kponyo | 3.3243°N; 24.17641°E | BDA313 | 1 | 0 | 1 | |
| Bas Uele | Kponyo | 3.3243°N; 24.17641°E | BDA338 | 1 | 0 | 1 | |
| Bas Uele | Kponyo | 3.3243°N; 24.17641°E | BDA341 | 1 | 0 | 1 | |
| Tshopo | Masako | 0.60841667°N; 25.2609167°E | M011 | 1 | 0 | 0 | |
| Tshopo | Masako | 0.60841667°N; 25.2609167°E | M012 | 1 | 0 | 0 | |
| Tshopo | Masako | 0.60841667°N; 25.2609167°E | M018 | 1 | 0 | 0 | |
| Tshopo | Masako | 0.60841667°N; 25.2609167°E | M020 | 1 | 0 | 0 | |
| Tshopo | Masako | 0.60841667°N; 25.2609167°E | M029 | 1 | 0 | 0 | |

submitted (Sypplementary data)

| | | | | | | |
|---|---|---|---|---|---|---|
| Tshopo | Masako | 0.60841667°N; 25.2609167°E | M034 | 1 | 0 | 0 |
| Tshopo | Masako | 0.60841667°N; 25.2609167°E | M005 | 1 | 0 | 0 |
| Tshopo | Masako | 0.60841667°N; 25.2609167°E | M027 | 1 | 0 | 0 |
| Tshopo | Masako | 0.60841667°N; 25.2609167°E | M028 | 1 | 0 | 0 |
| Ituri | Ndeke 3 | 1.98676°N; 30.91548°E | NDU 803 | 0 | 0 | 1 |
| Tshopo | Obenge | 1.383222°N; 25.038694°E | TLL 154 | 0 | 0 | 1 |
| Tshopo | Obenge | 1.383222°N; 25.038694°E | TLL 155 | 0 | 0 | 0 |
| Tshopo | Obenge | 1.383222°N; 25.038694°E | TLL 58 | 0 | 0 | 1 |
| Tshopo | Obenge | 1.383222°N; 25.038694°E | TLL 59 | 0 | 0 | 1 |
| Tshopo | Yoko | 0.29638°N ; 25.28722°E | BY001 | 1 | 0 | 0 |
| Tshopo | Yoko | 0.29638°N ; 25.28722°E | BY002 | 1 | 0 | 1 |
| Tshopo | Yoko | 0.29638°N ; 25.28722°E | BY003 | 1 | 0 | 1 |
| Tshopo | Yoko | 0.29638°N ; 25.28722°E | BY004 | 1 | 0 | 1 |
| Tshopo | Yoko | 0.29638°N ; 25.28722°E | BY006 | 1 | 0 | 1 |
| Tshopo | Yoko | 0.29638°N ; 25.28722°E | BY007 | 1 | 0 | 1 |
| Tshopo | Yoko | 0.29638°N ; 25.28722°E | BY008 | 1 | 0 | 1 |
| Tshopo | Yoko | 0.29638°N ; 25.28722°E | BY009 | 1 | 0 | 0 |
| Tshopo | Yoko | 0.29638°N ; 25.28722°E | BY011 | 1 | 0 | 0 |
| Tshopo | Yoko | 0.29638°N ; 25.28722°E | BY013 | 1 | 0 | 1 |
| Tshopo | Yoko | 0.29638°N ; 25.28722°E | BY005 | 1 | 0 | 1 |
| Tshopo | Yoko | 0.29638°N ; 25.28722°E | BY012 | 1 | 0 | 1 |
| Tshopo | Yoko | 0.29638°N ; 25.28722°E | BY158 | 0 | 0 | 1 |
| Tshopo | Yoko | 0.29638°N ; 25.28722°E | Y007 | 1 | 0 | 0 |
| Tshopo | Yoko | 0.29638°N ; 25.28722°E | Y008 | 1 | 0 | 0 |
| Tshopo | Yoko | 0.29638°N ; 25.28722°E | Y009 | 1 | 0 | 0 |
| Tshopo | Yoko | 0.29638°N ; 25.28722°E | Y010 | 1 | 0 | 0 |
| Tshopo | Yoko | 0.29638°N ; 25.28722°E | Y012 | 1 | 1 | 1 |
| Tshopo | Yoko | 0.29638°N ; 25.28722°E | Y015 | 1 | 0 | 1 |

| | | | | | | |
|---|---|---|---|---|---|---|
| Tshopo | Yoko | 0.29638°N ; 25.28722°E | Y016 | 1 | 0 | 0 |
| Tshopo | Yoko | 0.29638°N ; 25.28722°E | Y017 | 1 | 0 | 0 |
| Tshopo | Yoko | 0.29638°N ; 25.28722°E | Y018 | 1 | 0 | 0 |
| Tshopo | Yoko | 0.29638°N ; 25.28722°E | Y023 | 1 | 0 | 0 |
| Tshopo | Yoko | 0.29638°N ; 25.28722°E | Y031 | 1 | 0 | 0 |
| Tshopo | Yoko | 0.29638°N ; 25.28722°E | Y034 | 1 | 0 | 1 |
| Tshopo | Yoko | 0.29638°N ; 25.28722°E | Y043 | 1 | 0 | 0 |
| Tshopo | Yoko | 0.29638°N ; 25.28722°E | Y044 | 1 | 0 | 0 |
| Tshopo | Yoko | 0.29638°N ; 25.28722°E | Y045 | 1 | 0 | 1 |
| Tshopo | Yoko | 0.29638°N ; 25.28722°E | Y046 | 1 | 0 | 0 |
| Tshopo | Yoko | 0.29638°N ; 25.28722°E | Y048 | 1 | 0 | 0 |
| Tshopo | Yoko | 0.29638°N ; 25.28722°E | Y050 | 1 | 0 | 1 |
| Tshopo | Yoko | 0.29638°N ; 25.28722°E | Y057 | 1 | 1 | 1 |
| Tshopo | Yoko | 0.29638°N ; 25.28722°E | Y064 | 1 | 0 | 1 |
| Tshopo | Yoko | 0.29638°N ; 25.28722°E | Y071 | 1 | 0 | 1 |
| Tshopo | Yoko | 0.29638°N ; 25.28722°E | Y083 | 1 | 0 | 1 |
| Tshopo | Yoko | 0.29638°N ; 25.28722°E | Y084 | 1 | 0 | 1 |
| Tshopo | Yoko | 0.29638°N ; 25.28722°E | Y085 | 1 | 0 | 0 |
| Tshopo | Yoko | 0.29638°N ; 25.28722°E | Y097 | 1 | 0 | 0 |
| Tshopo | Yoko | 0.29638°N ; 25.28722°E | Y119 | 1 | 0 | 1 |
| Tshopo | Yoko | 0.29638°N ; 25.28722°E | Y120 | 1 | 0 | 0 |
| Tshopo | Yoko | 0.29638°N ; 25.28722°E | Y028 | 1 | 0 | 0 |
| Tshopo | Yoko | 0.29638°N ; 25.28722°E | Y074 | 1 | 1 | 1 |
| Tshopo | Yoko | 0.29638°N ; 25.28722°E | Y082 | 1 | 0 | 0 |
| Tshopo | Yoko | 0.29638°N ; 25.28722°E | Y100 | 1 | 0 | 0 |
| Tshopo | Yoko | 0.29638°N ; 25.28722°E | Y101 | 1 | 0 | 1 |
| Tshopo | Yoko | 0.29638°N ; 25.28722°E | YK-086 | 1 | 0 | 0 |
| Tshopo | Yoko | 0.29638°N ; 25.28722°E | YK-090 | 1 | 0 | 0 |

[illegible] et al., submitted (Sypplementary data)

| | | | | | | |
|---|---|---|---|---|---|---|
| Tshopo | Yoko | 0.29638°N ; 25.28722°E | YK-001 | 1 | 0 | 0 |
| Tshopo | Yoko | 0.29638°N ; 25.28722°E | YK-003 | 1 | 0 | 0 |
| Tshopo | Yoko | 0.29638°N ; 25.28722°E | YK-004 | 1 | 0 | 0 |
| Tshopo | Yoko | 0.29638°N ; 25.28722°E | YK-005 | 1 | 0 | 0 |
| Tshopo | Yoko | 0.29638°N ; 25.28722°E | Yk-006 | 1 | 0 | 0 |
| Tshopo | Yoko | 0.29638°N ; 25.28722°E | Yk-007 | 1 | 0 | 0 |
| Tshopo | Yoko | 0.29638°N ; 25.28722°E | Yk-009 | 1 | 0 | 0 |
| Tshopo | Yoko | 0.29638°N ; 25.28722°E | YK-015 | 1 | 0 | 0 |
| Tshopo | Yoko | 0.29638°N ; 25.28722°E | YK-018 | 1 | 0 | 0 |
| Tshopo | Yoko | 0.29638°N ; 25.28722°E | YK-020 | 1 | 0 | 0 |
| Tshopo | Yoko | 0.29638°N ; 25.28722°E | YK-024 | 0 | 0 | 0 |
| Tshopo | Yoko | 0.29638°N ; 25.28722°E | YK-026 | 1 | 0 | 0 |
| Tshopo | Yoko | 0.29638°N ; 25.28722°E | YK-029 | 1 | 0 | 0 |
| Tshopo | Yoko | 0.29638°N ; 25.28722°E | YK-033 | 1 | 0 | 1 |
| Tshopo | Yoko | 0.29638°N ; 25.28722°E | YK-034 | 1 | 0 | 0 |
| Tshopo | Yoko | 0.29638°N ; 25.28722°E | YK-035 | 1 | 0 | 1 |
| Tshopo | Yoko | 0.29638°N ; 25.28722°E | YK-042 | 1 | 0 | 0 |
| Tshopo | Yoko | 0.29638°N ; 25.28722°E | YK-043 | 1 | 0 | 0 |
| Tshopo | Yoko | 0.29638°N ; 25.28722°E | YK-045 | 1 | 0 | 0 |
| Tshopo | Yoko | 0.29638°N ; 25.28722°E | YK-058 | 1 | 0 | 0 |
| Tshopo | Yoko | 0.29638°N ; 25.28722°E | YK-061 | 1 | 0 | 0 |
| Tshopo | Yoko | 0.29638°N ; 25.28722°E | YK-066 | 1 | 0 | 0 |
| Tshopo | Yoko | 0.29638°N ; 25.28722°E | YK-073 | 1 | 0 | 0 |
| Tshopo | Yoko | 0.29638°N ; 25.28722°E | YK-075 | 1 | 0 | 0 |
| Tshopo | Yoko | 0.29638°N ; 25.28722°E | YK-077 | 1 | 0 | 0 |
| Tshopo | Yoko | 0.29638°N ; 25.28722°E | YK-078 | 1 | 0 | 0 |
| Tshopo | Yoko | 0.29638°N ; 25.28722°E | YK-079 | 0 | 0 | 0 |
| Tshopo | Yoko | 0.29638°N ; 25.28722°E | YK-082 | 1 | 0 | 0 |

Bange et al., submitted (Sypplementary data)

| | | | | | | |
|---|---|---|---|---|---|---|
| Tshopo | Yoko | 0.29638°N ; 25.28722°E | YK-084 | 1 | 0 | 0 |
| Tshopo | Yoko | 0.29638°N ; 25.28722°E | YK-085 | 1 | 0 | 0 |
| Tshopo | Yoko | 0.29638°N ; 25.28722°E | YK 093 | 1 | 0 | 0 |
| Tshopo | Yoko | 0.29638°N ; 25.28722°E | YK 094 | 1 | 0 | 0 |
| Tshopo | Yoko | 0.29638°N ; 25.28722°E | YK 096 | 0 | 0 | 0 |
| Tshopo | Yoko | 0.29638°N ; 25.28722°E | YK 097 | 1 | 0 | 0 |
| Tshopo | Yoko | 0.29638°N ; 25.28722°E | YK 098 | 1 | 0 | 0 |
| Tshopo | Yoko | 0.29638°N ; 25.28722°E | YK 100 | 1 | 0 | 1 |
| Tshopo | Yoko | 0.29638°N ; 25.28722°E | YK 101 | 1 | 0 | 1 |
| Tshopo | Yoko | 0.29638°N ; 25.28722°E | YK 102 | 1 | 0 | 1 |
| Tshopo | Yoko | 0.29638°N ; 25.28722°E | YK 104 | 1 | 0 | 0 |
| Tshopo | Yoko | 0.29638°N ; 25.28722°E | YK 105 | 1 | 0 | 1 |
| Tshopo | Yoko | 0.29638°N ; 25.28722°E | YK 106 | 1 | 0 | 1 |
| Tshopo | Yoko | 0.29638°N ; 25.28722°E | YK 108 | 1 | 0 | 1 |
| Tshopo | Yoko | 0.29638°N ; 25.28722°E | YK 111 | 1 | 0 | 0 |
| Tshopo | Yoko | 0.29638°N ; 25.28722°E | YK 114 | 1 | 0 | 1 |
| Tshopo | Yoko | 0.29638°N ; 25.28722°E | YK 115 | 1 | 0 | 1 |
| Tshopo | Yoko | 0.29638°N ; 25.28722°E | YK 118 | 1 | 0 | 0 |
| Tshopo | Yoko | 0.29638°N ; 25.28722°E | YK 122 | 1 | 0 | 0 |
| Tshopo | Yoko | 0.29638°N ; 25.28722°E | YK 123 | 1 | 0 | 0 |
| Tshopo | Yoko | 0.29638°N ; 25.28722°E | YK 130 | 1 | 0 | 0 |
| Tshopo | Yoko | 0.29638°N ; 25.28722°E | YK 147 | 1 | 0 | 0 |
| Tshopo | Yoko | 0.29638°N ; 25.28722°E | YK 154 | 1 | 0 | 0 |
| Tshopo | Yoko | 0.29638°N ; 25.28722°E | YK 155 | 1 | 0 | 0 |
| Tshopo | Yoko | 0.29638°N ; 25.28722°E | YK 157 | 1 | 0 | 0 |
| Tshopo | Yoko | 0.29638°N ; 25.28722°E | YK 160 | 1 | 0 | 0 |
| Tshopo | Yoko | 0.29638°N ; 25.28722°E | YK 165 | 1 | 0 | 0 |
| Tshopo | Yoko | 0.29638°N ; 25.28722°E | YK 179 | 1 | 0 | 0 |

et al., submitted (Sypplementary data)

| | | | | | | |
|---|---|---|---|---|---|---|
| Tshopo | Yoko | 0.29638°N ; 25.28722°E | YK 180 | 1 | 0 | 0 |
| Tshopo | Yoko | 0.29638°N ; 25.28722°E | YK 181 | 1 | 0 | 0 |
| Tshopo | Yoko | 0.29638°N ; 25.28722°E | YK 182 | 1 | 0 | 0 |
| Tshopo | Yoko | 0.29638°N ; 25.28722°E | YK 187 | 1 | 0 | 0 |
| Tshopo | Yoko | 0.29638°N ; 25.28722°E | YK 194 | 1 | 0 | 0 |
| Tshopo | Yoko | 0.29638°N ; 25.28722°E | YK 197 | 1 | 0 | 0 |
| Tshopo | Yoko | 0.29638°N ; 25.28722°E | YK 198 | 1 | 0 | 0 |
| Tshopo | Yoko | 0.29638°N ; 25.28722°E | YK 199 | 1 | 0 | 0 |
| Tshopo | Yoko | 0.29638°N ; 25.28722°E | YK 201 | 1 | 0 | 0 |
| Tshopo | Yoko | 0.29638°N ; 25.28722°E | YK 202 | 1 | 0 | 0 |
| Tshopo | Yoko | 0.29638°N ; 25.28722°E | YK 213 | 1 | 0 | 0 |
| Tshopo | Yoko | 0.29638°N ; 25.28722°E | YK 224 | 1 | 0 | 0 |
| Tshopo | Yoko | 0.29638°N ; 25.28722°E | YK 230 | 1 | 0 | 0 |
| Tshopo | Yoko | 0.29638°N ; 25.28722°E | YK 233 | 1 | 0 | 0 |
| Tshopo | Yoko | 0.29638°N ; 25.28722°E | YK 234 | 1 | 0 | 0 |
| Tshopo | Yoko | 0.29638°N ; 25.28722°E | YK 236 | 1 | 0 | 1 |
| Tshopo | Yoko | 0.29638°N ; 25.28722°E | YK 237 | 1 | 0 | 0 |
| Tshopo | Yoko | 0.29638°N ; 25.28722°E | YK 238 | 1 | 0 | 0 |
| Tshopo | Yoko | 0.29638°N ; 25.28722°E | YK 239 | 1 | 0 | 0 |
| Tshopo | Yoko | 0.29638°N ; 25.28722°E | YK 241 | 1 | 0 | 0 |
| Tshopo | Yoko | 0.29638°N ; 25.28722°E | YK 242 | 1 | 0 | 0 |
| Tshopo | Yoko | 0.29638°N ; 25.28722°E | YK 245 | 1 | 0 | 0 |
| Tshopo | Yoko | 0.29638°N ; 25.28722°E | YK 251 | 1 | 0 | 0 |
| Tshopo | Yoko | 0.29638°N ; 25.28722°E | YK-027 | 1 | 0 | 0 |
| Tshopo | Yoko | 0.29638°N ; 25.28722°E | YK-041 | 1 | 0 | 0 |
| Tshopo | Yoko | 0.29638°N ; 25.28722°E | YK-080 | 1 | 0 | 0 |
| Tshopo | Yoko | 0.29638°N ; 25.28722°E | YK 103 | 1 | 0 | 0 |
| Tshopo | Yoko | 0.29638°N ; 25.28722°E | YK 109 | 1 | 0 | 1 |

D[illegible] et al., submitted (Sypplementary data)

| | | | | | | |
|---|---|---|---|---|---|---|
| Tshopo | Yoko | 0.29638°N ; 25.28722°E | YK 116 | 1 | 0 | 0 |
| Tshopo | Yoko | 0.29638°N ; 25.28722°E | YK 119 | 1 | 0 | 0 |
| Tshopo | Yoko | 0.29638°N ; 25.28722°E | YK 148 | 1 | 0 | 1 |
| Tshopo | Yoko | 0.29638°N ; 25.28722°E | YK 163 | 1 | 0 | 0 |
| Tshopo | Yoko | 0.29638°N ; 25.28722°E | YK 167 | 1 | 0 | 0 |
| Tshopo | Yoko | 0.29638°N ; 25.28722°E | YK 172 | 1 | 0 | 0 |
| Tshopo | Yoko | 0.29638°N ; 25.28722°E | YK 175 | 1 | 0 | 0 |
| Tshopo | Yoko | 0.29638°N ; 25.28722°E | YK 177 | 1 | 0 | 0 |
| Tshopo | Yoko | 0.29638°N ; 25.28722°E | YK 178 | 1 | 0 | 0 |
| Tshopo | Yoko | 0.29638°N ; 25.28722°E | YK 183 | 1 | 0 | 0 |
| Tshopo | Yoko | 0.29638°N ; 25.28722°E | YK 184 | 1 | 0 | 0 |
| Tshopo | Yoko | 0.29638°N ; 25.28722°E | YK 185 | 1 | 0 | 0 |
| Tshopo | Yoko | 0.29638°N ; 25.28722°E | YK 186 | 1 | 0 | 0 |
| Tshopo | Yoko | 0.29638°N ; 25.28722°E | YK 188 | 1 | 0 | 0 |
| Tshopo | Yoko | 0.29638°N ; 25.28722°E | YK 189 | 1 | 0 | 0 |
| Tshopo | Yoko | 0.29638°N ; 25.28722°E | YK 191 | 1 | 0 | 0 |
| Tshopo | Yoko | 0.29638°N ; 25.28722°E | YK 203 | 1 | 0 | 0 |
| Tshopo | Yoko | 0.29638°N ; 25.28722°E | YK 204 | 1 | 0 | 0 |
| Tshopo | Yoko | 0.29638°N ; 25.28722°E | YK 205 | 1 | 0 | 0 |
| Tshopo | Yoko | 0.29638°N ; 25.28722°E | YK 206 | 1 | 0 | 0 |
| Tshopo | Yoko | 0.29638°N ; 25.28722°E | YK 212 | 1 | 0 | 0 |
| Tshopo | Yoko | 0.29638°N ; 25.28722°E | YK 216 | 1 | 0 | 0 |
| Tshopo | Yoko | 0.29638°N ; 25.28722°E | YK 223 | 1 | 0 | 1 |
| Tshopo | Yoko | 0.29638°N ; 25.28722°E | YK 225 | 1 | 0 | 0 |
| Tshopo | Yoko | 0.29638°N ; 25.28722°E | YK 228 | 1 | 0 | 0 |
| Tshopo | Yoko | 0.29638°N ; 25.28722°E | YK 229 | 1 | 0 | 0 |
| Tshopo | Yoko | 0.29638°N ; 25.28722°E | YK 231 | 1 | 0 | 0 |
| Tshopo | Yoko | 0.29638°N ; 25.28722°E | YK 232 | 1 | 0 | 0 |

[illegible] et al., submitted (Sypplementary data)

| | | | | | | |
|---|---|---|---|---|---|---|
| Tshopo | Yoko | 0.29638°N ; 25.28722°E | YK 235 | 1 | 0 | 0 |
| Tshopo | Yoko | 0.29638°N ; 25.28722°E | YK 243 | 1 | 0 | 0 |
| Tshopo | Yoko | 0.29638°N ; 25.28722°E | YK 244 | 1 | 0 | 0 |
| Tshopo | Yoko | 0.29638°N ; 25.28722°E | YK 246 | 1 | 0 | 0 |
| Tshopo | Yoko | 0.29638°N ; 25.28722°E | YK 247 | 1 | 0 | 0 |
| Tshopo | Yoko | 0.29638°N ; 25.28722°E | YK 248 | 1 | 0 | 0 |
| Tshopo | Yoko | 0.29638°N ; 25.28722°E | YK 249 | 1 | 0 | 0 |
| Ituri | Epulu | 1.40948°N; 28.5716°E | EPL014 | 1 | 0 | 1 |
| Ituri | Epulu | 1.40948°N; 28.5716°E | EPL026 | 1 | 0 | 1 |
| Ituri | Epulu | 1.40948°N; 28.5716°E | EPL 035 | 1 | 0 | 0 |
| Ituri | Epulu | 1.40948°N; 28.5716°E | EPL 036 | 1 | 0 | 0 |
| Ituri | Epulu | 1.40948°N; 28.5716°E | EPL 040 | 1 | 0 | 0 |
| Ituri | Epulu | 1.40948°N; 28.5716°E | EPL 050 | 1 | 0 | 0 |
| Ituri | Epulu | 1.40948°N; 28.5716°E | EPL 056 | 1 | 0 | 0 |
| Bas Uele | Sukisa | 2.31579°N; 24.98309°E | RBTL 347 | 1 | 0 | 0 |
| Bas Uele | Sukisa | 2.31579°N; 24.98309°E | RBTL 349 | 1 | 0 | 0 |
| Bas Uele | Sukisa | 2.31579°N; 24.98309°E | RBTL 444 | 1 | 0 | 0 |
| Bas Uele | Sukisa | 2.31579°N; 24.98309°E | RBTL 445 | 1 | 0 | 0 |
| Bas Uele | Sukisa | 2.31579°N; 24.98309°E | RBTL 446 | 1 | 0 | 0 |
| Tshopo | UMA | 0.55252778°N; 25.9243611°E | UM 018 | 1 | 0 | 0 |
| Tshopo | UMA | 0.55252778°N; 25.9243611°E | UM 029 | 1 | 0 | 1 |
| Tshopo | UMA | 0.55252778°N; 25.9243611°E | UM 030 | 1 | 0 | 0 |
| Tshopo | UMA | 0.55252778°N; 25.9243611°E | UMA385 | 0 | 0 | 1 |
| Tshopo | UMA | 0.55252778°N; 25.9243611°E | UM 387 | 1 | 0 | 1 |
| Tshopo | UMA | 0.55252778°N; 25.9243611°E | UM 389 | 1 | 1 | 1 |
| Tshopo | UMA | 0.55252778°N; 25.9243611°E | UM 416 | 1 | 0 | 0 |
| Tshopo | UMA | 0.55252778°N; 25.9243611°E | UM 439 | 1 | 0 | 0 |
| Tshopo | UMA | 0.55252778°N; 25.9243611°E | UM 469 | 1 | 0 | 0 |

| | | | | | | | |
|---|---|---|---|---|---|---|---|
| Bas Uele | Wela | 2.7380556°N; 23.7827777°E | AKT-107 | 1 | 0 | 1 | |
| Bas Uele | Wela | 2.7380556°N; 23.7827777°E | AKT-236 | 1 | 0 | 1 | |
| Tshopo | Yasikia | 0.3645°N; 25.0064°E | YSK011 | 1 | 0 | 1 | |
| Tshopo | Yasikia | 0.3645°N; 25.0064°E | YSK012 | 1 | 0 | 1 | |
| Tshopo | Yasikia | 0.3645°N; 25.0064°E | YSK026 | 1 | 0 | 1 | |
| Tshopo | Yasikia | 0.3645°N; 25.0064°E | YSK027 | 1 | 0 | 1 | |
| Tshopo | Yasikia | 0.3645°N; 25.0064°E | YSK037 | 1 | 0 | 1 | |
| Tshopo | Yasikia | 0.3645°N; 25.0064°E | YSK102 | 1 | 0 | 0 | |
| Tshopo | Yasikia | 0.3645°N; 25.0064°E | YSK133 | 1 | 0 | 0 | |
| Tshopo | Yasikia | 0.3645°N; 25.0064°E | YSK062 | 1 | 0 | 0 | |
| Tshopo | Yasikia | 0.3645°N; 25.0064°E | YSK079 | 1 | 0 | 0 | |
| Tshopo | Yasikia | 0.3645°N; 25.0064°E | YSK100 | 1 | 0 | 0 | |
| Tshopo | Yasikia | 0.3645°N; 25.0064°E | YSK101 | 1 | 0 | 0 | |
| Tshopo | Yasikia | 0.3645°N; 25.0064°E | YSK131 | 1 | 0 | 0 | |
| Tshopo | Yasikia | 0.3645°N; 25.0064°E | YSK132 | 1 | 0 | 0 | |
| | | | MF694313 | 0 | 0 | 1 | GenBank |
| | | | MF694315 | 0 | 0 | 1 | GenBank |
| | | | MF694317 | 0 | 0 | 1 | GenBank |
| | | | MF694318 | 0 | 0 | 1 | GenBank |
| | | | MN597594 | 0 | 0 | 1 | GenBank |
| | | | MN597668 | 0 | 0 | 1 | GenBank |
| Bas Uele | Bombongolo | 3.05659°N; 23.36565°E | UAC 928 | 1 | 1 | 1 | |
| Bas Uele | Bombongolo | 3.05659°N; 23.36565°E | UAC 933 | 1 | 1 | 1 | |
| Bas Uele | Bombongolo | 3.05659°N; 23.36565°E | UAC 937 | 1 | 1 | 1 | |
| Bas Uele | Bombongolo | 3.05659°N; 23.36565°E | UAC 938 | 1 | 1 | 1 | |
| Bas Uele | Kponyo | 3.3243°N; 24.17641°E | BDA 12 | 1 | 1 | 1 | |
| Bas Uele | Kponyo | 3.3243°N; 24.17641°E | BDA 63 | 1 | 1 | 1 | |
| Bas Uele | Kponyo | 3.3243°N; 24.17641°E | BDA 81 | 1 | 1 | 1 | |

| | | | | | | |
|---|---|---|---|---|---|---|
| Bas Uele | Kponyo | 3.3243°N; 24.17641°E | BDA 222 | 1 | 1 | 1 |
| Bas Uele | Kponyo | 3.3243°N; 24.17641°E | BDA 223 | 1 | 1 | 1 |
| Bas Uele | Kponyo | 3.3243°N; 24.17641°E | BDA 239 | 1 | 1 | 1 |
| Bas Uele | Kponyo | 3.3243°N; 24.17641°E | BDA 243 | 1 | 1 | 1 |
| Bas Uele | Kponyo | 3.3243°N; 24.17641°E | BDA 300 | 1 | 1 | 1 |
| Bas Uele | Kponyo | 3.3243°N; 24.17641°E | BDA 337 | 1 | 1 | 1 |
| Ituri | Epulu | 1.40948°N; 28.5716°E | EPL 002 | 1 | 1 | 1 |
| Ituri | Epulu | 1.40948°N; 28.5716°E | EPL 011 | 1 | 1 | 1 |
| Ituri | Epulu | 1.40948°N; 28.5716°E | EPL 013 | 1 | 1 | 1 |
| Ituri | Epulu | 1.40948°N; 28.5716°E | EPL 017 | 1 | 1 | 1 |
| Ituri | Epulu | 1.40948°N; 28.5716°E | EPL 018 | 1 | 1 | 1 |
| Ituri | Epulu | 1.40948°N; 28.5716°E | EPL 029 | 1 | 1 | 1 |
| Ituri | Epulu | 1.40948°N; 28.5716°E | EPL 012 | 1 | 1 | 1 |
| Ituri | Epulu | 1.40948°N; 28.5716°E | EPL 028 | 1 | 1 | 1 |
| Tshopo | Uma | 0.55252778°N; 25.9243611°E | UM 025 | 1 | 1 | 1 |
| Tshopo | Uma | 0.55252778°N; 25.9243611°E | UM 388 | 1 | 1 | 1 |
| Tshuapa | Inkanamongo | 0.71375°N; 20.530778°E | EBO 287 | 1 | 1 | 0 |
| Tshopo | Yoko | 0.29638°N ; 25.28722°E | Y001 | 1 | 1 | 0 |
| Tshopo | Yoko | 0.29638°N ; 25.28722°E | Y006 | 1 | 1 | 0 |
| Tshopo | Yoko | 0.29638°N ; 25.28722°E | Y020 | 1 | 1 | 0 |
| Tshopo | Yoko | 0.29638°N ; 25.28722°E | Y027 | 1 | 1 | 0 |
| Tshopo | Yoko | 0.29638°N ; 25.28722°E | Y029 | 1 | 1 | 0 |
| Tshopo | Yoko | 0.29638°N ; 25.28722°E | Y030 | 1 | 1 | 0 |
| Tshopo | Yoko | 0.29638°N ; 25.28722°E | Y035 | 1 | 1 | 0 |
| Tshopo | Yoko | 0.29638°N ; 25.28722°E | Y052 | 1 | 1 | 0 |
| Tshopo | Yoko | 0.29638°N ; 25.28722°E | Y056 | 1 | 1 | 0 |
| Tshopo | Yoko | 0.29638°N ; 25.28722°E | Y072 | 1 | 1 | 0 |
| Tshopo | Yoko | 0.29638°N ; 25.28722°E | Y075 | 1 | 1 | 0 |

| | | | | | | |
|---|---|---|---|---|---|---|
| Tshopo | Yoko | 0.29638°N ; 25.28722°E | Y077 | 1 | 1 | 0 |
| Tshopo | Yoko | 0.29638°N ; 25.28722°E | Y096 | 1 | 1 | 0 |
| Tshopo | Yoko | 0.29638°N ; 25.28722°E | Y098 | 1 | 1 | 0 |
| Tshopo | Yoko | 0.29638°N ; 25.28722°E | Y113 | 1 | 1 | 0 |
| Tshopo | Yoko | 0.29638°N ; 25.28722°E | Y114 | 1 | 1 | 0 |
| Tshopo | Yoko | 0.29638°N ; 25.28722°E | Y115 | 1 | 1 | 0 |
| Tshopo | Yoko | 0.29638°N ; 25.28722°E | Y121 | 1 | 1 | 0 |
| Tshopo | Yoko | 0.29638°N ; 25.28722°E | Y122 | 1 | 1 | 0 |
| Tshopo | Yoko | 0.29638°N ; 25.28722°E | YK 002 | 1 | 1 | 0 |
| Tshopo | Yoko | 0.29638°N ; 25.28722°E | YK 008 | 1 | 1 | 0 |
| Tshopo | Yoko | 0.29638°N ; 25.28722°E | YK 011 | 1 | 1 | 0 |
| Tshopo | Yoko | 0.29638°N ; 25.28722°E | YK 013 | 1 | 1 | 0 |
| Tshopo | Yoko | 0.29638°N ; 25.28722°E | YK 014 | 1 | 1 | 0 |
| Tshopo | Yoko | 0.29638°N ; 25.28722°E | YK 017 | 1 | 1 | 0 |
| Tshopo | Yoko | 0.29638°N ; 25.28722°E | YK 021 | 1 | 1 | 0 |
| Tshopo | Yoko | 0.29638°N ; 25.28722°E | YK 022 | 1 | 1 | 0 |
| Tshopo | Yoko | 0.29638°N ; 25.28722°E | YK 025 | 1 | 1 | 0 |
| Tshopo | Yoko | 0.29638°N ; 25.28722°E | YK 028 | 1 | 1 | 0 |
| Tshopo | Yoko | 0.29638°N ; 25.28722°E | YK 030 | 1 | 1 | 0 |
| Tshopo | Yoko | 0.29638°N ; 25.28722°E | YK 040 | 1 | 1 | 0 |
| Tshopo | Yoko | 0.29638°N ; 25.28722°E | YK 046 | 1 | 1 | 0 |
| Tshopo | Yoko | 0.29638°N ; 25.28722°E | YK 047 | 1 | 1 | 0 |
| Tshopo | Yoko | 0.29638°N ; 25.28722°E | YK 049 | 1 | 1 | 0 |
| Tshopo | Yoko | 0.29638°N ; 25.28722°E | YK 050 | 1 | 1 | 0 |
| Tshopo | Yoko | 0.29638°N ; 25.28722°E | YK 051 | 1 | 1 | 0 |
| Tshopo | Yoko | 0.29638°N ; 25.28722°E | YK 053 | 1 | 1 | 0 |
| Tshopo | Yoko | 0.29638°N ; 25.28722°E | YK 055 | 1 | 1 | 0 |
| Tshopo | Yoko | 0.29638°N ; 25.28722°E | YK 057 | 1 | 1 | 0 |

submitted (Sypplementary data)

| | | | | | | |
|---|---|---|---|---|---|---|
| Tshopo | Yoko | 0.29638°N ; 25.28722°E | YK 060 | 1 | 1 | 0 |
| Tshopo | Yoko | 0.29638°N ; 25.28722°E | YK 065 | 1 | 1 | 0 |
| Tshopo | Yoko | 0.29638°N ; 25.28722°E | YK 067 | 1 | 1 | 0 |
| Tshopo | Yoko | 0.29638°N ; 25.28722°E | YK 068 | 1 | 1 | 0 |
| Tshopo | Yoko | 0.29638°N ; 25.28722°E | YK 069 | 1 | 1 | 0 |
| Tshopo | Yoko | 0.29638°N ; 25.28722°E | YK 072 | 1 | 1 | 0 |
| Tshopo | Yoko | 0.29638°N ; 25.28722°E | YK 076 | 1 | 1 | 0 |
| Tshopo | Yoko | 0.29638°N ; 25.28722°E | YK 087 | 1 | 1 | 0 |
| Tshopo | Yoko | 0.29638°N ; 25.28722°E | YK 088 | 1 | 1 | 0 |
| Tshopo | Yoko | 0.29638°N ; 25.28722°E | YK 091 | 1 | 1 | 0 |
| Tshopo | Yoko | 0.29638°N ; 25.28722°E | YK 095 | 1 | 1 | 0 |
| Tshopo | Yoko | 0.29638°N ; 25.28722°E | YK 117 | 1 | 1 | 0 |
| Tshopo | Yoko | 0.29638°N ; 25.28722°E | YK 146 | 1 | 1 | 0 |
| Tshopo | Yoko | 0.29638°N ; 25.28722°E | YK 159 | 1 | 1 | 0 |
| Tshopo | Yoko | 0.29638°N ; 25.28722°E | YK 166 | 1 | 1 | 0 |
| Tshopo | Yoko | 0.29638°N ; 25.28722°E | YK 214 | 1 | 1 | 1 |
| Tshopo | Yoko | 0.29638°N ; 25.28722°E | YK 219 | 1 | 1 | 0 |
| Tshopo | Yoko | 0.29638°N ; 25.28722°E | YK 220 | 1 | 1 | 0 |
| Tshopo | Yoko | 0.29638°N ; 25.28722°E | YK 221 | 1 | 1 | 0 |
| Tshopo | Yoko | 0.29638°N ; 25.28722°E | YK 222 | 1 | 1 | 0 |
| Tshopo | Yoko | 0.29638°N ; 25.28722°E | Y011 | 1 | 1 | 0 |
| Tshopo | Yoko | 0.29638°N ; 25.28722°E | Y062 | 1 | 1 | 0 |
| Tshopo | Yoko | 0.29638°N ; 25.28722°E | Y086 | 1 | 1 | 0 |
| Tshopo | Yoko | 0.29638°N ; 25.28722°E | Y110 | 1 | 1 | 0 |
| Tshopo | Yoko | 0.29638°N ; 25.28722°E | YK 023 | 1 | 0 | 0 |
| Tshopo | Yoko | 0.29638°N ; 25.28722°E | YK 063 | 1 | 0 | 0 |
| Tshopo | Yoko | 0.29638°N ; 25.28722°E | YK 099 | 1 | 0 | 0 |
| Tshopo | Yoko | 0.29638°N ; 25.28722°E | YK 120 | 1 | 0 | 0 |

et al., submitted (Sypplementary data)

| | | | | | | |
|---|---|---|---|---|---|---|
| Tshopo | Yoko | 0.29638°N ; 25.28722°E | YK 121 | 1 | 0 | 0 |
| Tshopo | Yoko | 0.29638°N ; 25.28722°E | YK 124 | 1 | 0 | 0 |
| Tshopo | Yoko | 0.29638°N ; 25.28722°E | YK 126 | 1 | 0 | 0 |
| Tshopo | Yoko | 0.29638°N ; 25.28722°E | YK 127 | 1 | 0 | 0 |
| Tshopo | Yoko | 0.29638°N ; 25.28722°E | YK 131 | 1 | 0 | 0 |
| Tshopo | Yoko | 0.29638°N ; 25.28722°E | YK 150 | 1 | 0 | 0 |
| Tshopo | Yoko | 0.29638°N ; 25.28722°E | YK 153 | 1 | 0 | 0 |
| Tshopo | Yoko | 0.29638°N ; 25.28722°E | YK 156 | 1 | 1 | 1 |
| Tshopo | Yoko | 0.29638°N ; 25.28722°E | YK 161 | 1 | 1 | 0 |
| Tshopo | Yoko | 0.29638°N ; 25.28722°E | YK 196 | 1 | 0 | 1 |
| Tshopo | Yoko | 0.29638°N ; 25.28722°E | YK 207 | 1 | 1 | 0 |
| Tshopo | Yoko | 0.29638°N ; 25.28722°E | YK 215 | 1 | 0 | 0 |
| Tshopo | Yoko | 0.29638°N ; 25.28722°E | YK 226 | 1 | 0 | 0 |
| Tshopo | Yoko | 0.29638°N ; 25.28722°E | Y057 | 1 | 1 | 1 |
| Tshopo | Yoko | 0.29638°N ; 25.28722°E | Y063 | 1 | 1 | 1 |
| Tshopo | Yoko | 0.29638°N ; 25.28722°E | Y065 | 1 | 1 | 1 |
| Tshopo | Yoko | 0.29638°N ; 25.28722°E | YK 012 | 1 | 1 | 1 |
| Tshopo | Yoko | 0.29638°N ; 25.28722°E | YK 037 | 1 | 1 | 1 |
| Tshopo | Yoko | 0.29638°N ; 25.28722°E | YK 038 | 1 | 1 | 1 |
| Tshopo | Yoko | 0.29638°N ; 25.28722°E | YK 054 | 1 | 1 | 1 |
| Tshopo | Yoko | 0.29638°N ; 25.28722°E | YK 059 | 1 | 1 | 1 |
| Tshopo | Yoko | 0.29638°N ; 25.28722°E | YK 064 | 1 | 1 | 1 |
| Tshopo | Yoko | 0.29638°N ; 25.28722°E | YK 107 | 1 | 1 | 1 |
| Tshopo | Yoko | 0.29638°N ; 25.28722°E | YK 112 | 1 | 1 | 1 |
| Tshopo | Yoko | 0.29638°N ; 25.28722°E | YK 113 | 1 | 1 | 1 |
| Tshopo | Yoko | 0.29638°N ; 25.28722°E | YK 129 | 1 | 1 | 1 |
| Tshopo | Yoko | 0.29638°N ; 25.28722°E | YK 149 | 1 | 1 | 1 |
| Tshopo | Yoko | 0.29638°N ; 25.28722°E | YK 200 | 1 | 1 | 1 |

submitted (Sypplementary data)

| | | | | | | |
|---|---|---|---|---|---|---|
| Tshopo | Yoko | 0.29638°N ; 25.28722°E | YK 039 | 1 | 1 | 1 |
| Ituri | Epulu | 1.40948°N; 28.5716°E | EPL 008 | 1 | 1 | 1 |
| Ituri | Epulu | 1.40948°N; 28.5716°E | EPL 020 | 1 | 1 | 1 |
| Tshopo | Uma | 0.55252778°N; 25.9243611°E | UM 040 | 1 | 1 | 0 |
| Tshopo | Yoko | 0.29638°N ; 25.28722°E | Y081 | 1 | 1 | 1 |
| Tshopo | Yoko | 0.29638°N ; 25.28722°E | YK 110 | 1 | 1 | 1 |
| Tshopo | Yoko | 0.29638°N ; 25.28722°E | YK 128 | 1 | 1 | 1 |
| Tshopo | Yoko | 0.29638°N ; 25.28722°E | YK 158 | 1 | 0 | 0 |
| Tshopo | Yoko | 0.29638°N ; 25.28722°E | YK 210 | 1 | 1 | 1 |
| Tshopo | Yoko | 0.29638°N ; 25.28722°E | Y033 | 1 | 1 | 1 |
| Tshopo | Yoko | 0.29638°N ; 25.28722°E | Y042 | 1 | 1 | 1 |
| Tshopo | Yoko | 0.29638°N ; 25.28722°E | YK 152 | 1 | 1 | 1 |
| Tshopo | Yoko | 0.29638°N ; 25.28722°E | YK 190 | 1 | 1 | 1 |
| Tshopo | Yoko | 0.29638°N ; 25.28722°E | YK 192 | 1 | 1 | 1 |
| Tshopo | Yoko | 0.29638°N ; 25.28722°E | YK 195 | 1 | 1 | 1 |
| Bas Uele | Ahupa | 2.65748°N; 23.79874°E | UAC 901 | 1 | 1 | 0 |
| Bas Uele | Ahupa | 2.65748°N; 23.79874°E | UAC 913 | 1 | 1 | 0 |
| Tshuapa | Inkanamongo | 0.71375°N; 20.530778°E | EBO 50 | 1 | 1 | 0 |
| Tshuapa | Inkanamongo | 0.71375°N; 20.530778°E | EBO 230 | 1 | 1 | 0 |
| Ituri | Epulu | 1.40948°N; 28.5716°E | EPL 025 | 1 | 1 | 0 |
| Ituri | Epulu | 1.40948°N; 28.5716°E | EPL 031 | 1 | 1 | 0 |
| Ituri | Epulu | 1.40948°N; 28.5716°E | EPL 041 | 1 | 1 | 0 |
| Ituri | Epulu | 1.40948°N; 28.5716°E | EPL 045 | 1 | 1 | 0 |
| Ituri | Epulu | 1.40948°N; 28.5716°E | EPL 046 | 1 | 1 | 0 |
| Ituri | Epulu | 1.40948°N; 28.5716°E | EPL 047 | 1 | 1 | 0 |
| Ituri | Epulu | 1.40948°N; 28.5716°E | EPL 048 | 1 | 1 | 0 |
| Ituri | Epulu | 1.40948°N; 28.5716°E | EPL 049 | 1 | 1 | 0 |
| Ituri | Epulu | 1.40948°N; 28.5716°E | EPL 052 | 1 | 1 | 0 |

et al., submitted (Sypplementary data)

| | | | | | | |
|---|---|---|---|---|---|---|
| Ituri | Epulu | 1.40948°N; 28.5716°E | EPL 053 | 1 | 1 | 0 |
| Tshopo | Uma | 0.55252778°N; 25.9243611°E | UM 041 | 1 | 1 | 1 |
| Tshopo | Uma | 0.55252778°N; 25.9243611°E | UM 383 | 1 | 1 | 0 |
| Bas Uele | Wela | 2.7380556°N; 23.7827777°E | AKT-106 | 1 | 1 | 0 |
| Bas Uele | Wela | 2.7380556°N; 23.7827777°E | AKT-257 | 1 | 1 | 0 |
| Bas Uele | Ahupa | 2.65748°N; 23.79874°E | UAC 917 | 1 | 1 | 0 |
| Tshuapa | Inkanamongo | 0.71375°N; 20.530778°E | EBO 245 | 1 | 1 | 0 |
| Tshuapa | Inkanamongo | 0.71375°N; 20.530778°E | EBO 472 | 1 | 1 | 0 |
| Tshopo | Mombongo | 1.645278°N; 23.159722°E | YHM337 | 1 | 1 | 0 |
| Tshopo | Mombongo | 1.645278°N; 23.159722°E | YHM338 | 1 | 1 | 0 |
| Tshopo | Yoko | 0.29638°N ; 25.28722°E | YK 036 | 1 | 1 | 1 |
| Tshopo | Yoko | 0.29638°N ; 25.28722°E | YK 074 | 1 | 0 | 0 |
| Tshopo | Yoko | 0.29638°N ; 25.28722°E | YK 151 | 1 | 1 | 0 |
| Tshopo | Masako | 0.60841667°N; 25.2609167°E | M006 | 1 | 1 | 0 |
| Tshopo | Masako | 0.60841667°N; 25.2609167°E | M008 | 1 | 1 | 0 |
| Tshopo | Masako | 0.60841667°N; 25.2609167°E | M010 | 1 | 1 | 0 |
| Tshopo | Masako | 0.60841667°N; 25.2609167°E | M032 | 1 | 1 | 0 |
| Tshopo | Masako | 0.60841667°N; 25.2609167°E | M033 | 1 | 1 | 0 |
| Ituri | Epulu | 1.40948°N; 28.5716°E | EPL 032 | 1 | 1 | 0 |
| Ituri | Epulu | 1.40948°N; 28.5716°E | EPL 034 | 1 | 1 | 0 |
| Ituri | Epulu | 1.40948°N; 28.5716°E | EPL 042 | 1 | 1 | 0 |
| Ituri | Epulu | 1.40948°N; 28.5716°E | EPL 043 | 1 | 1 | 0 |
| Ituri | Epulu | 1.40948°N; 28.5716°E | EPL 051 | 1 | 1 | 0 |
| Tshopo | Uma | 0.55252778°N; 25.9243611°E | UM 019 | 1 | 1 | 1 |
| Tshopo | Uma | 0.55252778°N; 25.9243611°E | UM 384 | 1 | 1 | 0 |
| Tshopo | Uma | 0.55252778°N; 25.9243611°E | UM 440 | 1 | 1 | 0 |
| Tshopo | Uma | 0.55252778°N; 25.9243611°E | UM 510 | 1 | 1 | 0 |
| Bas Uele | Bogala | 2.95623°N; 23.53877°E | UAC 919 | 1 | 1 | 0 |

et al., submitted (Sypplementary data)

| | | | | | | | |
|---|---|---|---|---|---|---|---|
| Bas Uele | Bogala | 2.95623°N; 23.53877°E | UAC 922 | 1 | 1 | 0 | |
| Bas Uele | Bombongolo | 3.05659°N; 23.36565°E | UAC 934 | 1 | 1 | 0 | |
| Tshopo | Mbiye | 0.45013889°N; 25.29930556°E | MBIYE 525 | 1 | 1 | 0 | |
| Tshopo | Mbiye | 0.45013889°N; 25.29930556°E | MBIYE 526 | 1 | 1 | 0 | |
| Tshopo | Mbiye | 0.45013889°N; 25.29930556°E | MBIYE 528 | 1 | 1 | 0 | |
| Bas Uele | Kponyo | 3.3243°N; 24.17641°E | BDA 80 | 1 | 1 | 0 | |
| Bas Uele | Kponyo | 3.3243°N; 24.17641°E | BLI01 | 1 | 1 | 0 | |
| Tshopo | Yoko | 0.29638°N ; 25.28722°E | Y021 | 1 | 1 | 0 | |
| Tshopo | Yoko | 0.29638°N ; 25.28722°E | YK 070 | 1 | 0 | 0 | |
| Tshopo | Yoko | 0.29638°N ; 25.28722°E | YK 083 | 1 | 1 | 0 | |
| Tshopo | Yoko | 0.29638°N ; 25.28722°E | YK 193 | 1 | 1 | 0 | |
| Tshopo | Yoko | 0.29638°N ; 25.28722°E | YK 211 | 1 | 1 | 0 | |
| Ituri | Epulu | 1.40948°N; 28.5716°E | EPL 023 | 1 | 1 | 0 | |
| Ituri | Epulu | 1.40948°N; 28.5716°E | EPL 044 | 1 | 1 | 0 | |
| Ituri | Epulu | 1.40948°N; 28.5716°E | EPL 054 | 1 | 1 | 0 | |
| Bas Uele | Sukisa | 2.31579°N; 24.98309°E | RBTL 348 | 1 | 1 | 0 | |
| Bas Uele | Sukisa | 2.31579°N; 24.98309°E | RBTL 476 | 1 | 1 | 0 | |
| Tshopo | Uma | 0.55252778°N; 25.9243611°E | UM 386 | 1 | 1 | 0 | |
| Kwilu | Kikwit | -5.038308°N; 18.817844°E | KIK 675 | 0 | 1 | 1 | MRAC |
| Kwilu | Kikwit | -5.038308°N; 18.817844°E | KIK 883 | 0 | 1 | 0 | MRAC |
| Kwilu | Kikwit | -5.038308°N; 18.817844°E | KIK 881 | 0 | 1 | 0 | MRAC |
| Kinshasa | Kinshasa | -4.325°S; 15322222°E | RMCA 17346-M- | 0 | 1 | 0 | MRAC |
| Kinshasa | Kinshasa | -4.325°S; 15322222°E | RMCA 14053-M- | 0 | 1 | 0 | MRAC |
| Kinshasa | Kinshasa | -4.325°S; 15322222°E | RMCA 17354-M- | 0 | 1 | 0 | MRAC |
| Kinshasa | Kinshasa | -4.325°S; 15322222°E | RMCA 17349-M- | 0 | 1 | 0 | MRAC |
| Kinshasa | Kinshasa | -4.325°S; 15322222°E | RMCA 17800-M- | 0 | 1 | 0 | MRAC |

Baele et al., submitted (Sypplementary data)

| | | | | | | | |
|---|---|---|---|---|---|---|---|
| Kinshasa | Kinshasa | -4.325°S; 15322222°E | RMCA 17355-M- | 0 | 1 | 0 | MRAC |
| Kinshasa | Kinshasa | -4.325°S; 15322222°E | RMCA 17348-M- | 0 | 1 | 0 | MRAC |
| Kinshasa | Kinshasa | -4.325°S; 15322222°E | RMCA 17799-M- | 0 | 1 | 0 | MRAC |
| Kinshasa | Kinshasa | -4.325°S; 15322222°E | RMCA 17350-M- | 0 | 1 | 0 | MRAC |
| Kinshasa | Kinshasa | -4.325°S; 15322222°E | RMCA 14054-M- | 0 | 1 | 0 | MRAC |
| Kinshasa | Kinshasa | -4.325°S; 15322222°E | RMCA 17330-M- | 0 | 1 | 0 | MRAC |
| Kinshasa | Kinshasa | -4.325°S; 15322222°E | RMCA 17802-M- | 0 | 1 | 0 | MRAC |
| Kinshasa | Kinshasa | -4.325°S; 15322222°E | RMCA 17353-M- | 0 | 1 | 0 | MRAC |
| Kinshasa | Kinshasa | -4.325°S; 15322222°E | RMCA 17992-M- | 0 | 1 | 0 | MRAC |
| Kongo Central | Kisantu | -5.1338889°S; 15.058333°E | RMCA 720A-M- | 0 | 1 | 0 | MRAC |
| Kongo Central | Kisantu | -5.1338889°S; 15.058333°E | RMCA 720B-M- | 0 | 1 | 0 | MRAC |
| Kinshasa | Kinshasa | -4.325°S; 15322222°E | RMCA 17351-M- | 0 | 1 | 0 | MRAC |
| Kinshasa | Kinshasa | -4.325°S; 15322222°E | RMCA 31116-M- | 0 | 1 | 0 | MRAC |
| Kinshasa | Kinshasa | -4.325°S; 15322222°E | RMCA 17352-M- | 0 | 1 | 0 | MRAC |
| Kinshasa | Kinshasa | -4.325°S; 15322222°E | RMCA 9563-M- | 0 | 1 | 0 | MRAC |
| Kinshasa | Kinshasa | -4.325°S; 15322222°E | RMCA 17652-M- | 0 | 1 | 0 | MRAC |
| Kinshasa | Kinshasa | -4.325°S; 15322222°E | RMCA 17991-M- | 0 | 1 | 0 | MRAC |

Baelo et al., submitted (Sypplementary data)

| | | | | | | | |
|---|---|---|---|---|---|---|---|
| Kinshasa | Kinshasa | -4.325°S; 15322222°E | RMCA 31115-M- | 0 | 1 | 0 | MRAC |
| Kongo Central | Kongo Central | -5.816667°S; 13.483333°E | RMCA 123-M- | 0 | 1 | 0 | MRAC |
| Kinshasa | Kinshasa | -4.325°S; 15322222°E | RMCA 31117-M- | 0 | 1 | 0 | MRAC |

**Table S2.** Craniometric data (M1 to M23) of *Funisciurus* specimens examined (M1: maximum length of the skull; M2: condilo-basal length; M3: henselion-basion length; M4: henselion-palation length; M5: length of the cleft palate; M6: length of the diastema; M7: distance between the anterior edge of the alveolus of M1 and the cutting edge of the upper incisor; M8: width of the interorbital narrowing; M9: width of the zygomatic arch; M10: minimum width of the palate; M11: length of the upper molar row; M12 : external width of upper molar row at M1; M13: maximum molar width; M14: minimum width of zygomatic plate; M15: maximum width of nasals; M16: maximum length of nasals; M17: length of lower molar row; M18: length of the tympanic bulla; M19: maximum width of the cranial cavity; M20: depth of the upper incisors; M21: height of the rostrum at the anterior edge of the alveolus of M1; M22: maximum width of the rostrum; M23: distance between the extreme points of the coronoid and angular processes).

| **Label** | **M1** | **M2** | **M3** | **M4** | **M5** | **M6** | **M7** | **M8** | **M9** | **M10** | **M11** | **M12** | **M13** | **M14** | **M15** | **M16** | **M17** | **M18** | **M19** | **M20** | **M21** | **M22** | **M23** |
|---|---|---|---|---|---|---|---|---|---|---|---|---|---|---|---|---|---|---|---|---|---|---|---|
| **UAC 928** | 45.3 | 43.1 | 38.4 | 21.9 | 3.2 | 11.3 | 13.3 | 11.7 | 24.9 | 3.85 | 8 | 10 | 2.75 | 3 | 7.2 | 12.5 | 8 | 9.1 | 15.9 | 2.6 | 10 | 10 | 15.3 |
| **UAC 933** | 45.8 | 43.9 | 38.8 | 22.1 | 3 | 11.1 | 13.2 | 12.2 | 25.7 | 3.9 | 8 | 10.2 | 2.7 | 2.9 | 7 | 12 | 8.6 | 8.9 | 15.6 | 2.36 | 9.8 | 10.1 | 15.8 |
| **UAC 937** | 45.4 | 43.3 | 38.4 | 22.1 | 3.2 | 11.2 | 13.1 | 11 | 24.9 | 4.09 | 8.2 | 10 | 2.6 | 3.1 | 7.3 | 12.1 | 8 | 8.8 | 15.7 | 2.3 | 10 | 10.1 | 15.8 |
| **UAC 938** | 46.6 | 44.3 | 38.7 | 22.7 | 3.1 | 11.3 | 13.5 | 12.1 | 25.8 | 4.3 | 7.8 | 10.2 | 2.8 | 3 | 7.2 | 12.4 | 8.7 | 8.6 | 16 | 2.41 | 10.7 | 10.3 | 16.5 |
| **BDA 12** | 45.7 | 43.8 | 38.1 | 22.8 | 2.98 | 12.2 | 13.4 | 11.5 | | 4.75 | 7.89 | 9.99 | 2.39 | 3.52 | 7.62 | 12.5 | 9.26 | 8.12 | 16.1 | 2.43 | 10.6 | 10.7 | 16.5 |
| **BDA 63** | 45.3 | 43.7 | 37.7 | 21.5 | 2.9 | 12.1 | 12.8 | 11 | 25.3 | 3.8 | 8.2 | 10.2 | 2.4 | 2.9 | 6.9 | 12.4 | 8.1 | 8.8 | 16.2 | 2.4 | 10 | 10.1 | 15.2 |
| **BDA 81** | 45.9 | 43.9 | 38.4 | 21.9 | 3.75 | 11.2 | 12.6 | 11 | 25.9 | 4.99 | 8.38 | 10.2 | 2.57 | 3.67 | 7.25 | 12.8 | 8.19 | 8.75 | 16.6 | 2.5 | 10.9 | 10.4 | 15.8 |
| **BDA 222** | 44.7 | 42.1 | 37 | 21.5 | 3.22 | 11.4 | 13.1 | 10.4 | 25.2 | 4.45 | 7.9 | 10.2 | 2.64 | 3.33 | 7.16 | 12 | 9 | 9.09 | 16.2 | 2.21 | 10 | 10.7 | 15.1 |
| **BDA 223** | 45.9 | 43.5 | 38.3 | 21.7 | 2.81 | 12.2 | 13.2 | 12.1 | 25.7 | 4.32 | 8.23 | 10.2 | 2.58 | 2.85 | 7.6 | 12.6 | 8.45 | 8 | 16.6 | 2.33 | 10.2 | 10.9 | 16.1 |
| **BDA 239** | 47.1 | 45.2 | 39.2 | 23 | 3.3 | 12 | 14.4 | 12 | 26.1 | 4.33 | 8.42 | 10.8 | 2.75 | 3.13 | 6.98 | 12.7 | 8.18 | 9.04 | 16.7 | 2.48 | 10.4 | 10.6 | 15.7 |
| **BDA 243** | 47 | 45.7 | 39.7 | 22.9 | 2.86 | 12.7 | 14.4 | 11.7 | 26.6 | 3.88 | 8.44 | 10.8 | 2.61 | 3.02 | 7.51 | 12.6 | 8.37 | 9.79 | 16.4 | 2.4 | 11.3 | 10.5 | 16.1 |
| **BDA 300** | 44.7 | 42.7 | 37.1 | 21.3 | 3.09 | 11.2 | 12.6 | 10.6 | 24.9 | 4.28 | 8.26 | 10.1 | 2.76 | 2.86 | 7.5 | 11.5 | 8.57 | 9.06 | 15.7 | 2.75 | 10 | 10.1 | 15.3 |
| **BDA 337** | 46 | 43.9 | 37.6 | 21.6 | 3.12 | 11.9 | 13.2 | 11 | 25.1 | 4.55 | 8.13 | 10.1 | 2.88 | 3.02 | 7.41 | 12.3 | 8.66 | 8.32 | 16.2 | 2.6 | 10.5 | 10.2 | 15.6 |
| **EPL 002** | 46.4 | 43.7 | 38.1 | 22 | 3.44 | 11.1 | 12.5 | 11.1 | 25.8 | 4.76 | 8.51 | 10.3 | 2.69 | 3.88 | 6.5 | 12.7 | 8.1 | 9.18 | 16.3 | 2.3 | 10.6 | 10.2 | 14.8 |

| | | | | | | | | | | | | | | | | | | | | | | | |
|---|---|---|---|---|---|---|---|---|---|---|---|---|---|---|---|---|---|---|---|---|---|---|---|
| **EPL 011** | 44.9 | 41.7 | 37.5 | 20.9 | 2.4 | 11.4 | 12.5 | 11.4 | 24 | 4.4 | 7.42 | 10.4 | 2.4 | 3.76 | 6.03 | 11.6 | 8.1 | 8.21 | 16.2 | 2.25 | 10.4 | 10.2 | 14.6 |
| **EPL 013** | 45.8 | 43.9 | 38 | 22.2 | 3.04 | 11.4 | 12.6 | 10.6 | 24.9 | 3.94 | 8.5 | 10.5 | 2.66 | 4.01 | 6.62 | 11.6 | 8.3 | 9.38 | 16 | 2.52 | 10.7 | 9.81 | 15.3 |
| **EPL 017** | 45.6 | 42 | 36.9 | 21.4 | 3.04 | 11.5 | 12.7 | 11 | 25.1 | 3.76 | 7.9 | 9.99 | 2.21 | 4.04 | 5.94 | 12.3 | 8.17 | 9.02 | 15.9 | 2.44 | 10 | 10.1 | 14.3 |
| **EPL 018** | 44.8 | 42.9 | 36.9 | 21.4 | 3.16 | 10.6 | 12.2 | 11.9 | 25.5 | 3.67 | 8.06 | 10.5 | 2.58 | 3.68 | 6.84 | 11.5 | 8.1 | 9.3 | 15.7 | 2.4 | 10.1 | 10.3 | 15.5 |
| **EPL 029** | 46.8 | 44.7 | 39 | 22.6 | 3.63 | 11.9 | 13.3 | 12 | 26.2 | 3.95 | 8.78 | 10.6 | 2.63 | 3.94 | 6.61 | 11.9 | 8.48 | 9.37 | 16.2 | 2.67 | 10.9 | 10 | 15.6 |
| **EPL 012** | 45.5 | 44.1 | | 22.1 | 2.93 | 12.1 | 13.7 | 11.5 | 25.8 | 3.9 | 8.3 | 10.3 | 2.4 | | 7.3 | 12.2 | 7.8 | 9.09 | 16.2 | 2.6 | 11 | 10.5 | 15.2 |
| **EPL 028** | 43.8 | 42.1 | 37.1 | 21.3 | 2.8 | 11.2 | 13.1 | 11 | 24.2 | 3.91 | 8.2 | 10.1 | 2.4 | 3.2 | 7 | 12.1 | 7.81 | 8 | 15.5 | 2.6 | 10.8 | 10.4 | 15.6 |
| **UM 025** | 46.2 | 43.2 | 38.7 | 21.9 | 3.12 | 10.3 | 12.7 | 12 | 25.9 | 4.09 | 8.1 | 10.4 | 2.15 | 3.21 | 7.23 | 13.8 | 7.95 | 9.06 | 15.7 | 2.4 | 10.7 | 10.7 | 15 |
| **UM 388** | 43.9 | 41.2 | 36.9 | 21 | 3.37 | 10.4 | 12.6 | 12 | 24 | 4.13 | 8.43 | 10.5 | 2.45 | 3.25 | 6.83 | 12.7 | 8.07 | 9.31 | 16 | 2.7 | 11 | 9.84 | 15.3 |
| **EBO 287** | 46.9 | 44.7 | 39.3 | 22.9 | 3.35 | 12.9 | 13.9 | 11.8 | 26.8 | 4.55 | 7.98 | 11 | 2.66 | 3.15 | 7.52 | 13 | 8.36 | 9.42 | 16 | 2.21 | 10.8 | 10.9 | 16.2 |
| **Y001** | 47.1 | 44.7 | 39.6 | 22.7 | 3.58 | 11.9 | 14.1 | 11.9 | 26.2 | 4.28 | 8.22 | 10.3 | 2.55 | 3.14 | 6.68 | 12.9 | 8.08 | 10.3 | 17.8 | 2.43 | 11.6 | 11.8 | |
| **Y006** | 46.6 | 44.1 | 38.7 | 21.7 | 3.37 | 11.5 | 13.1 | 11.3 | 25.4 | 4.18 | 8.95 | 10.4 | 2.45 | 3.16 | 7.01 | 12.7 | 8.26 | 9.54 | 18.4 | 2.27 | 11.6 | 10.6 | 15 |
| **Y020** | 49.6 | 46.2 | 41.3 | 24.2 | 3.65 | 13.3 | 15.1 | 12.6 | 28.1 | 4.24 | 8.38 | 11.5 | 2.67 | 3.85 | 7.13 | 14.2 | 8.39 | 9.91 | 19 | 2.44 | 12.7 | 12.2 | 17.3 |
| **Y027** | 47.8 | 44.6 | 39.5 | 22.6 | 3.27 | 12.5 | 14 | 11.5 | 26.1 | 4.02 | 8.39 | 11.3 | 2.5 | 3.7 | 6.65 | 12.9 | 8.18 | 9.35 | 17.4 | 2.13 | 11.2 | 12 | 16.2 |
| **Y029** | 45.4 | 43.1 | 38.2 | 22 | 3.19 | 12.1 | 13.9 | 11.2 | 24.3 | 4.11 | 8 | 10.2 | 2.29 | 3.98 | 6.92 | 11.8 | 7.4 | 9 | 17.6 | 2.36 | 11.6 | 11.1 | 15.8 |
| **Y030** | 47.8 | 45.4 | 39.2 | 22.8 | 3.57 | 12.7 | 14 | 11.7 | 25.8 | 4.97 | 8.79 | 10.5 | 2.46 | 4.45 | 6.81 | 13.5 | 8.39 | 9.29 | 18.9 | 2.21 | 11.4 | 10.6 | 15.1 |
| **Y035** | 45.2 | 42.4 | 37.2 | 21.5 | 3.61 | 11.6 | 13.1 | 10.6 | | 3.72 | 7.89 | 10.4 | 2.35 | 3.2 | 6.72 | 12 | 7.72 | 10.2 | 17.7 | 2.06 | 11.3 | 10 | 15.1 |
| **Y052** | 45.7 | | | | | 11.4 | 13.1 | 11.2 | 24.9 | | | | | 3.37 | | | 8.06 | 9.53 | 17.7 | 2.43 | 11.3 | | 14.4 |
| **Y056** | 47.7 | 44.2 | 38.7 | 22.6 | 3.76 | 12.2 | 13.5 | 11.1 | 26.8 | 4.7 | 8.32 | 11 | 2.72 | 2.55 | 7.01 | 13.3 | 7.96 | 9.49 | 18.5 | | 11.9 | 11 | |
| **Y072** | 49 | 45.6 | 40.9 | 23.2 | 3 | 12.9 | 14.8 | 11.9 | 27.1 | 3.9 | 8.2 | 10.6 | 2.62 | 3.21 | 7.05 | 13.1 | 7.96 | 9.7 | 17.9 | 2.4 | 12 | 11.4 | 16.9 |
| **Y075** | 47.4 | 44.7 | 39.5 | 22.3 | 3.25 | 12.3 | 14.6 | 11.5 | 25.8 | 4.89 | 8.67 | 10.6 | 2.65 | 4.06 | 7.38 | 13 | 8.2 | 9.44 | 18.9 | 2.3 | 11.6 | 11.2 | 15.8 |
| **Y077** | 46.5 | 43 | 39.3 | 22 | 3.51 | 11.6 | 12.9 | 11.2 | 25.5 | 3.73 | 8.2 | 10.1 | 2.66 | 3.67 | 6.53 | 12.5 | 8.21 | 9.74 | 18.3 | 2.24 | 11.5 | 10.8 | 15.8 |
| **Y096** | 47.1 | 44.1 | 38.5 | 22.5 | 3.28 | 12.5 | 14.4 | 11.9 | 26.2 | 4.3 | 8.02 | 11.2 | 2.61 | 3.35 | 6.79 | 13.3 | 7.75 | 9.12 | 18 | 2.28 | 11.3 | 11.1 | 15.2 |
| **Y098** | 46.5 | 43.9 | 38.3 | 21.7 | 3.25 | 12.1 | 13.3 | 11 | 25.3 | 3.68 | 8.54 | 10.6 | 2.58 | 3.1 | 6.5 | 12.7 | 8.33 | 9.83 | 18.4 | 2.14 | 11.4 | 10.4 | 15.7 |
| **Y113** | 47.7 | 45.3 | 40.4 | 22.9 | 3.83 | 12.5 | 14.2 | 12 | 25.8 | 4.35 | 8.35 | 10.8 | 2.38 | 3.01 | 7.05 | 12.6 | 7.85 | 9.64 | 18.6 | 2.21 | 11.5 | 11.5 | 15.7 |
| **Y114** | 48.4 | 45.5 | 39.7 | 23.3 | 3.34 | 11.9 | 13.5 | 11.7 | 25.9 | 4.6 | 8.7 | 10.8 | 2.51 | 3.11 | 6.89 | 13.2 | 8.08 | 10.2 | 18.6 | 2.78 | 11.7 | 11.6 | 15.2 |
| **Y115** | 49.2 | 45.8 | 39.9 | 22.7 | 3.21 | 12.4 | 14 | 12.9 | 26.7 | 4.97 | 8.11 | 11.7 | 2.57 | 3.98 | 6.65 | 13.6 | 8.26 | 9.96 | 18.9 | 2.58 | 12.1 | 11.1 | 16 |
| **Y121** | 45.4 | 42.4 | 37.3 | 21 | 3.08 | 11.2 | 13 | 12.1 | 24.9 | 3.56 | 8.04 | 10.1 | 2.3 | 3.31 | 6.02 | 12.7 | 7.59 | 8.62 | 18.2 | 2.07 | 11.5 | 10.4 | 15.3 |
| **Y122** | 47.8 | 44.5 | 39.9 | 22.7 | 3.13 | 12.5 | 13.1 | 12.8 | 26.3 | 3.85 | 7.81 | 10.9 | 2.64 | 3.5 | 6.64 | 12.8 | 7.86 | 9.25 | 17.9 | 2.32 | 12.2 | 11.5 | |
| **YK 002** | 48.7 | 45.7 | 39.7 | 22 | 3.1 | 12.1 | 13.9 | 12.6 | | 3.95 | 9.2 | 11.8 | 2.85 | 3.45 | 6.9 | 13.7 | 8.1 | 9.1 | 18.1 | 2.3 | 11.3 | 11.6 | 16.8 |
| **YK 008** | 47 | 44.1 | 38.8 | 22 | 3.51 | 11.9 | 13.8 | 11.8 | 26.6 | 4.05 | 7.9 | 10.4 | 2.45 | 4.1 | 6.58 | 12.1 | 7.7 | 9.06 | 17.4 | 2.35 | 10.9 | 11 | |

Baele et al., submitted (Supplementary data)

| | | | | | | | | | | | | | | | | | | | | | | | |
|---|---|---|---|---|---|---|---|---|---|---|---|---|---|---|---|---|---|---|---|---|---|---|---|
| **YK 011** | 47.8 | 45 | 39.3 | 22.5 | 3.46 | 11.9 | 13.6 | 12.1 | | 4.03 | 8.63 | 10.3 | 2.81 | 4.01 | 6.66 | 13.2 | 8.47 | 9.4 | 18.4 | 2.26 | 11.8 | 11.3 | 15.5 |
| **YK 013** | 47.4 | 44.1 | 40 | 22.6 | 3.52 | 12.3 | 13.9 | 11.3 | 26.7 | 4.81 | 8.36 | 10.4 | 2.65 | 4.62 | 6.62 | 13.6 | 7.82 | 9.98 | 18.2 | 2.43 | 12.2 | 11.5 | 15.5 |
| **YK 014** | 47.8 | 44.6 | 39.8 | 22.8 | 3.42 | 12.1 | 13.9 | 12.1 | | 3.85 | 8.73 | 10.5 | 2.49 | 3.81 | 6.44 | 12.9 | 8.3 | 9.48 | 17.7 | 2.31 | 11.6 | 11.1 | 16.6 |
| **YK 017** | 44.1 | 41.7 | 36.1 | 21.4 | 3.16 | 11.4 | 12.8 | 9.99 | 24.7 | 3.62 | 8.29 | 10.1 | 2.47 | | 7 | 12.4 | 7.93 | 9.01 | 17 | 2.24 | 11.1 | 10.7 | 15.2 |
| **YK 021** | 48.9 | 46.7 | 40.5 | 23.3 | 3.3 | 12.3 | 14.2 | 12 | 25.9 | 4.6 | 8.3 | 10.6 | 2.35 | 3.75 | 7.5 | 13.5 | 8.1 | 10 | 18.5 | 2.5 | 11.2 | 11.5 | 15.2 |
| **YK 022** | 50.8 | 47.6 | 42 | 23.6 | 3.4 | 13.2 | 13.5 | 11.8 | | 5.1 | 10 | 12.5 | 2.75 | 3.4 | 7.85 | 13.3 | 9.3 | 10.9 | 18.7 | 2.9 | 12.2 | 12.3 | |
| **YK 025** | 48.1 | 45.3 | 39.8 | 23.4 | 3.5 | 11.8 | 14 | 11.9 | 26.4 | 3.87 | 8.21 | 10.8 | 2.31 | 2.9 | 6.87 | 13.2 | 7.25 | 9.6 | 18 | 2.19 | 11.3 | 10.9 | 15.2 |
| **YK 028** | 49.8 | 46.8 | 42.2 | 22.9 | 3.2 | 12.2 | 13.7 | 12.6 | 27.5 | 4.2 | 9.8 | 12.3 | 2.9 | 3.6 | 7.9 | 13.4 | 9.3 | 10.3 | 18 | 2.1 | 11.1 | 11.1 | 16.5 |
| **YK 030** | 44.1 | 42 | 36.9 | 21 | 3.28 | 11.5 | 12.5 | | | 4.1 | 8.6 | 10.2 | 2.34 | | 6.53 | 11.2 | 8.01 | 9.09 | 17.1 | 2.12 | | 11.4 | 14.6 |
| **YK 040** | 44.4 | 41.5 | 36.7 | 21 | 2.81 | 11.6 | 12.5 | 11 | 25.3 | 4.38 | 8.5 | 9.72 | 2.39 | 3.32 | 6.41 | 12.1 | 7.62 | 9.16 | 17.4 | 2.29 | 10.9 | 10.6 | 14.6 |
| **YK 046** | 45.2 | 43.1 | 37.1 | 21.1 | 3.57 | 12.1 | 13.7 | 11.5 | 25.3 | 4.12 | 8.19 | 10.4 | 2.07 | 3.81 | 6.94 | 11.3 | 7.74 | 9 | 17.5 | 2.55 | 11.6 | 10.9 | 15 |
| **YK 047** | 46.8 | 43.6 | 38 | 22.2 | | 12 | 13 | 11.7 | 25.4 | 4.44 | 8.31 | 10.5 | 2.17 | 3.29 | 7.33 | 12.6 | 8.27 | 9.43 | 17.8 | 2.32 | 12 | 11.1 | 15.9 |
| **YK 049** | 48.9 | 46.6 | 40.6 | 23.5 | 3.67 | 13 | 14.6 | 11.6 | 26.4 | 4.5 | 8.61 | 10.8 | 2.53 | 4.49 | 6.5 | 13.2 | 8.3 | 9.61 | 18.8 | 2.33 | 12.5 | 11.8 | 16.5 |
| **YK 050** | 47.5 | 44.4 | 39.2 | 22.7 | 3.6 | 13 | 14 | 12.4 | 26 | 3.49 | 7.84 | 10.7 | 2.35 | 3.33 | 6.32 | 13.1 | 7.57 | 9.46 | 17.6 | 2.33 | 11.8 | 10.9 | 15.4 |
| **YK 051** | 47.9 | 44.1 | 39.7 | 22.1 | 3.8 | 11.8 | 14.1 | 12.2 | 25.8 | 3.92 | 8.61 | 10.2 | 2.38 | 3.91 | 7 | 12.3 | 8.06 | 9.52 | 18.7 | 2.25 | 12 | 11.4 | 16 |
| **YK 053** | 45.6 | 42.5 | 38.2 | 21.5 | 3.28 | 11.4 | 12.9 | 11.9 | 24.7 | 4.61 | 8.5 | 10.3 | 2.3 | 3.99 | 6.91 | 12.3 | 8.26 | 9.4 | 17.4 | 2.21 | 11.1 | 11.4 | 15 |
| **YK 055** | 47.3 | 43.6 | 38.4 | 21.8 | 3.41 | 11.4 | 13.4 | 11.5 | 25.9 | 3.86 | 8.37 | 10.3 | 2.37 | 4.38 | 6.75 | 12.6 | 7.98 | 9.73 | 18.7 | 2.53 | 11.3 | 10.4 | 15.8 |
| **YK 057** | 47.4 | 43.9 | 39.9 | 21.9 | 3.86 | 11.5 | 13.5 | 11.4 | | 3.76 | 8.4 | 10.7 | 2.41 | 3.36 | 6.43 | 12.5 | 8.29 | 8.89 | 17.5 | 2.06 | 11.6 | 11.5 | 15.3 |
| **YK 060** | 46.5 | 43.3 | 38.9 | 22.4 | 3.6 | 12.5 | 14.2 | 11.4 | 25.8 | 3.64 | 8.19 | 9.93 | 2.34 | 3.44 | 6.54 | 13 | | 9.35 | 16.8 | 2.41 | 11.3 | 11.6 | |
| **YK 065** | 46.4 | 44.4 | 38.5 | 22.2 | 2.8 | 12.1 | 14.1 | 11 | | 4.61 | 7.83 | 10.3 | 2.13 | 4.46 | 6.28 | 12.4 | 7.97 | 9.46 | 17.7 | 2.45 | 11.1 | 10.8 | 14.5 |
| **YK 067** | 48.5 | 45.1 | 40.1 | 23 | 3.97 | 12.5 | 14.6 | 12.2 | 26.8 | 4.12 | 8.31 | 10.5 | 2.55 | 3.1 | 6.68 | 13.5 | 8.03 | 9.81 | 18.9 | 2.35 | 12 | 11.6 | 16.7 |
| **YK 068** | 48.5 | 44.8 | 40.4 | 22.2 | 3.3 | 12.1 | 13.5 | 11.5 | 27.5 | 3.63 | 8.52 | 10.4 | 2.35 | 3.58 | 6.39 | 13.4 | 8.15 | 9.23 | 17.9 | 2.22 | 12.3 | 11 | 15.5 |
| **YK 069** | 47.6 | 44.6 | 39.2 | 22.4 | 3.18 | 12.3 | 13.7 | 10.9 | | 3.85 | 8.85 | 10.3 | 2.35 | 4.9 | 6.5 | 12.6 | 8.33 | 9.66 | 18.1 | 2.39 | 11.4 | 10.7 | 14.9 |
| **YK 072** | 49 | 45.6 | 40.9 | 23.2 | 3 | 12.9 | 14.8 | 11.9 | 27.1 | 3.9 | 8.2 | 10.6 | 2.62 | 3.21 | 7.05 | 13.1 | 7.96 | 9.7 | 18.9 | 2.4 | 12 | 11.4 | 16.9 |
| **YK 076** | 46.9 | 44.6 | 40 | 22.4 | 3.5 | 12.3 | 14 | 11.3 | 25.6 | 3.37 | 8.44 | 10.4 | 2.31 | 3.78 | 6.52 | 12.6 | 8.39 | 9.56 | 18.4 | 2.2 | 11.6 | 10.4 | 15.2 |
| **YK 087** | 49.7 | 46.2 | 40.7 | 23.4 | 3.61 | 12.8 | 14.4 | 12.9 | | 3.97 | 8.66 | 11.3 | 2.53 | 3.47 | 6.75 | 13.9 | 8.14 | 9.26 | 18.3 | 2.31 | 11.9 | 12.3 | 16.5 |
| **YK 088** | 50.3 | 46.9 | 40.9 | 23.6 | 3.89 | 12.9 | 13.8 | 12.3 | | 4.83 | 8.79 | 11.4 | 2.4 | 4.2 | 7.78 | 13.8 | 8.05 | 9.68 | 19 | 2.43 | 11.5 | 11.4 | 16.8 |
| **YK 091** | 46.7 | 43.6 | 38.7 | 22 | 3.01 | 12.2 | 13.6 | 11.5 | | 4.07 | 8.01 | 10 | 2.66 | 3.53 | 7.33 | 12.6 | 7.95 | 9.67 | 17.9 | 2.4 | 11.6 | 11 | 14.6 |
| **YK 095** | 49.1 | 45.3 | 40.7 | 24 | 3.3 | 13.1 | 14.1 | 12 | 26.4 | 4.25 | 9.16 | 10.6 | 2.65 | 3.82 | 7.29 | 13.3 | 8.4 | 9.9 | 18 | 2.79 | 11.8 | 11.5 | 16 |
| **YK 117** | 46.4 | 44.4 | 39.1 | 22.3 | 2.71 | 11.4 | 13.2 | 11.3 | 25.4 | 4.11 | 8.27 | 10.2 | 2.72 | 3.62 | 7.04 | 12.3 | 8 | 9.66 | 17.3 | 2.58 | 10.3 | 11.1 | 15.2 |

Baele et al., submitted (Supplementary data)

| | | | | | | | | | | | | | | | | | | | | | | | |
|---|---|---|---|---|---|---|---|---|---|---|---|---|---|---|---|---|---|---|---|---|---|---|---|
| **YK 146** | 47.1 | 45.1 | 38.7 | 22.7 | 3.47 | 11.9 | 13.8 | 11.6 | 26.6 | 4.92 | 8.68 | 10.5 | 2.32 | 3.32 | 7.46 | 13.3 | 8.63 | 9.06 | 17.5 | 2.7 | 11.1 | 10.5 | 15.6 |
| **YK 159** | 47.6 | 44.7 | 40.3 | 22.3 | 3.03 | 12.5 | 13.4 | 12 | 26.2 | 3.7 | 8.6 | 10.4 | 2.39 | 3.15 | 7.43 | 13.3 | 8.48 | 9.5 | 17.2 | 2.53 | 10.6 | 11.6 | 16.1 |
| **YK 166** | 49.2 | 45.6 | 40.3 | 23.1 | 3.24 | 12.1 | 13.8 | 12.3 | 26.9 | 4.83 | 8.28 | 10.4 | 2.57 | 3.11 | 7.58 | 13.2 | 7.87 | 9.55 | 18.6 | 2.32 | 11.1 | 11.3 | 16.4 |
| **YK 214** | 44.8 | 42 | 37.8 | 21.8 | 2.9 | 11.6 | 13.3 | 11.2 | 24.1 | 3.59 | 8.75 | 10.1 | 2.4 | 3.26 | 6.75 | 11.8 | 7.58 | 9.17 | 17.2 | 2.51 | 10.2 | 11.2 | 14.9 |
| **YK 219** | 46.3 | 44.2 | 37.9 | 22.1 | 3.21 | 11.8 | 13.4 | 11.8 | 26 | 4.08 | 8.47 | 10.7 | 2.43 | 3.71 | 7.16 | 12.3 | 7.93 | 9.53 | 17.9 | 2.16 | 10.3 | 10.6 | 15.5 |
| **YK 220** | 46.9 | 44.3 | 39 | 22.2 | 2.8 | 12.4 | 13.6 | 12.1 | 25.8 | 4.09 | 8.49 | 10.4 | 2.4 | 2.94 | 7.16 | 12.5 | 7.92 | 9.63 | 17.6 | 2.4 | 10.3 | 10.6 | 16.3 |
| **YK 221** | 48.5 | 46.3 | 40 | 23 | 3.09 | 12.6 | 14.1 | 11.4 | 26.4 | 4.22 | 8.18 | 10.2 | 2.29 | 3.52 | 6.85 | 13.1 | 8.12 | 9.57 | 17.6 | 2.13 | 11.3 | 11.2 | 15.3 |
| **YK 222** | 48.8 | 46.4 | 41.1 | 23.2 | 2.8 | 12.7 | 14.3 | 12.4 | 27.4 | 3.62 | 8.87 | 10.8 | 2.25 | 3.38 | 8.06 | 13.2 | 8.56 | 9.27 | 17.7 | 2.13 | 10.9 | 11.7 | 15.7 |
| **Y011** | 43.6 | 40.1 | 35.4 | 19.5 | 2.29 | 10.1 | 11.5 | 10.7 | 23.8 | 3.12 | 7.23 | 9.9 | 2.44 | 4.21 | 5.9 | 11.8 | 7.33 | 9.25 | 16.8 | 2.52 | 10.4 | 10.3 | 14.6 |
| **Y062** | 43.1 | 40.2 | 35.8 | 20.8 | 2.25 | 9.96 | 11.4 | 10 | 23.4 | 3.57 | 7.71 | 9.87 | 2.79 | 2.34 | 6.2 | 10.9 | 7.74 | 8.99 | 15 | 2.11 | 11.3 | 10.6 | 15.7 |
| **Y086** | 42.7 | 39.8 | 34.6 | 19.2 | 2.61 | 9.74 | 11.6 | 9.99 | 22.9 | 4.01 | 7.37 | 9.62 | 2.34 | 2.05 | 6.06 | 10.2 | 7.61 | 8.84 | 15.3 | 2.51 | 11.2 | 10.7 | 16 |
| **Y110** | 42.1 | 39.9 | 34.8 | 19.5 | 2.61 | 9.75 | 11.4 | 10.2 | 24 | 3.7 | 7.2 | 9 | 2.67 | 2.5 | 6.08 | 10.2 | 7.44 | 9.08 | 15.3 | 2.28 | 10.9 | 10.6 | 15.2 |
| **YK 023** | 43.9 | 41.5 | 35.5 | 21 | 2.21 | 10.6 | 11 | 10.3 | 23.2 | 3.76 | 7.92 | 9.59 | 2.36 | 1.83 | 6.31 | 11.6 | 7.95 | 9.04 | 15.1 | 2.01 | 11.3 | 10.6 | |
| **YK 063** | 42.2 | 40.1 | 35.8 | 20.1 | 3.55 | 11 | 12.6 | 11 | 22.6 | 4.5 | 8.02 | 10.3 | 2.35 | 4.11 | 7.09 | 12.2 | 7.35 | 9.05 | 16.3 | 2.21 | 11.2 | 10.6 | |
| **YK 099** | 41.9 | 40 | 34.7 | 19.8 | 2.65 | 9.73 | 11.3 | 11 | 23.7 | 4.17 | 8.1 | 10.3 | 2.8 | 3.11 | 6.69 | 9.4 | 8.12 | 9.13 | 15 | 2.5 | 10.1 | 10.1 | 15.3 |
| **YK 120** | 43.7 | 41.4 | 36.4 | 20.7 | 2.86 | 10.1 | 12 | 11.5 | 24.8 | 4.42 | 8.33 | 10.7 | 2.53 | 2.53 | 7.5 | 9.94 | 8.34 | 9.7 | 16.8 | 2.3 | 10.1 | 9.88 | 15.8 |
| **YK 121** | 41.2 | 38.7 | 33.8 | 18.9 | 2.69 | 9.4 | 11 | 9.91 | 23 | 3.83 | 7.88 | 9.57 | 2.38 | 2.3 | 6.47 | 9.62 | 8.02 | 9.09 | 15.3 | 2.57 | 9.37 | 9.12 | 15 |
| **YK 124** | 42.5 | 40.4 | 34.5 | 19.3 | 2.69 | 9.88 | 11.2 | 10.2 | 23.6 | 4.79 | 8.06 | 9.37 | 2.29 | 2.91 | 6.85 | 10.6 | 7.85 | 9.01 | 15.6 | 2.22 | 9.85 | 9.2 | 15.4 |
| **YK 126** | 43.4 | 41.1 | 35.5 | 20 | 2.54 | 10.3 | 11.8 | 10.2 | 24 | 3.19 | 7.23 | 9.82 | 2.43 | 3.02 | 6.98 | 9.68 | 7.17 | 9.28 | 16.5 | 2.6 | 10.7 | 10.2 | 15 |
| **YK 127** | 41.7 | 39.9 | 34.2 | 19.8 | 2.4 | 9.47 | 11 | 10 | 22.1 | 3.09 | 7.42 | 8.86 | 2.43 | 2.8 | 7.29 | 10.2 | 7.6 | 8.94 | 15 | 2.4 | 9.69 | 9.6 | 15.2 |
| **YK 131** | 42.7 | 40.5 | 34.4 | 19.3 | 2.48 | 10.7 | 12.4 | 10.6 | 23.6 | 3.03 | 7.18 | 9.68 | 2.72 | 2.13 | 6.45 | 9.63 | 7.02 | 8.98 | 14.6 | 2.26 | 9.77 | 9.35 | 14.1 |
| **YK 150** | 43.2 | 40.6 | 34.9 | 20.1 | 2.84 | 10.3 | 11.3 | 11.2 | 23.1 | 3.98 | 7.86 | 9.8 | 2.38 | 2.16 | 7.17 | 10.6 | 8.2 | 9.03 | 15 | 2.72 | 10.1 | 9.48 | 14.7 |
| **YK 153** | 41.3 | 39.3 | 34.5 | 19.9 | 2.48 | 10.2 | 11.1 | 11.1 | 22.5 | 3.71 | 7.88 | 9.6 | 2.75 | 2.41 | 6.9 | 9.71 | 7.42 | 9.02 | 15.1 | 2.8 | 9.86 | 10.7 | 15.3 |
| **YK 156** | 41.7 | 39.7 | 34.5 | 19.9 | 2.59 | 10.6 | 11.7 | 10.6 | 22.6 | 3.48 | 7.56 | 9.56 | 2.35 | 2.67 | 6.54 | 10.6 | 7.75 | 9.13 | 15.1 | 2.59 | 10.1 | 10.3 | 15 |
| **YK 161** | 43.1 | 40.8 | 36.3 | 20.2 | 2.32 | 10.7 | 11.8 | 11.1 | 23.1 | 3.48 | 7.7 | 10.2 | 2.37 | 2.64 | 6.61 | 10.8 | 8.25 | 8.81 | 16.6 | 2.71 | 10.7 | 10.2 | 15 |
| **YK 196** | 40.9 | 39.3 | 34 | 19.4 | 2.21 | 9.58 | 11.1 | 10.7 | 22.4 | 3.7 | 7.3 | 9.45 | 2.3 | 2.75 | 6.3 | 10.4 | 7.32 | 9.06 | 14.2 | 2.14 | 9.71 | 9.45 | 15.4 |
| **YK 207** | 42 | 39.8 | 34.2 | 20.4 | 2.93 | 9.3 | 10.9 | 10.9 | 23 | 4.02 | 7.64 | 9.4 | 2.1 | 3.1 | 6.1 | 9.81 | 7.44 | 8.86 | 14.6 | 2.25 | 10.2 | 8.96 | 15.6 |
| **YK 215** | 42.1 | 40.5 | 34.1 | 19 | 2.3 | 9.66 | 11.6 | 10.5 | 23.1 | 3.2 | 7.91 | 9.85 | 2.44 | 2.99 | 6.02 | 10.5 | 7.51 | 9.2 | 15 | 2.07 | 9.48 | 9 | 15.4 |
| **YK 226** | 42 | 39.4 | 34.1 | 19.7 | 2.18 | 10 | 11.1 | 10.7 | 22.8 | 3.76 | 7.8 | 9.81 | 2.25 | 2.97 | 5.94 | 10.9 | 7.66 | 8.68 | 15.1 | 2.55 | 9.7 | 9.77 | |
| **Y057** | 48 | 45.2 | 40.2 | 23.4 | 3.17 | 12.4 | 14.4 | 12.3 | 27.1 | 3.85 | 8.3 | 10.7 | 2.52 | 3.2 | 6.51 | 13.3 | 7.89 | 9.7 | 18.8 | 2.27 | 12.2 | 11.1 | 15.8 |
| **Y063** | 49 | 46.1 | 40.2 | 24 | 3.11 | 12.8 | 15.4 | 13.1 | 26.7 | 4.38 | 8.18 | 11 | 2.57 | 4.09 | 7.85 | 13.6 | 7.8 | 9.69 | 19.3 | 2.45 | 12.2 | 11 | 15.8 |

Baek et al., submitted (Supplementary data)

| | | | | | | | | | | | | | | | | | | | | | | | |
|---|---|---|---|---|---|---|---|---|---|---|---|---|---|---|---|---|---|---|---|---|---|---|---|
| **Y065** | 47.5 | 43.7 | 39 | 22 | 3.21 | 11.8 | 13.6 | 11.7 | 25.5 | 4.31 | 8.23 | 10.4 | 2.63 | 3.95 | 6.48 | 12.8 | 7.81 | 9.65 | 18.7 | 2.1 | 11.8 | 11.4 | 14.7 |
| **YK 012** | 45.3 | 43.2 | 38.3 | 21.8 | 3.2 | 11.4 | 12.6 | 11.9 | 25.5 | 4.46 | 8.65 | 10.6 | 2.4 | 3.32 | 7.44 | 11.4 | 7.75 | 9.37 | 17.1 | 2.67 | 11.7 | 10.7 | 15 |
| **YK 037** | 45.3 | 42.4 | 38 | 21.5 | 2.96 | 11.4 | 12.8 | 10.7 | 23.9 | 3.83 | 7.84 | 10.2 | 2.33 | 2.39 | 6.45 | 11.1 | 7.7 | 9.6 | 16.8 | 2.15 | 11.1 | 11.2 | |
| **YK 038** | 45.4 | 42.8 | 37.6 | 21.4 | 2.92 | 12.5 | 12.7 | 11.7 | 24.6 | 3.86 | 8.06 | 9.82 | 2.4 | 3.58 | 6.69 | 11.3 | 8.46 | 9.56 | 17.9 | 2.22 | 11.2 | 11.4 | 15.2 |
| **YK 054** | 42.1 | 40.2 | 35.1 | 19.9 | 2.31 | 9.54 | 10.9 | 10.8 | 23.5 | 4.35 | 7.66 | 9.56 | 2.12 | 2.95 | 5.73 | 10.2 | 7.6 | 9.03 | 16.5 | 3.29 | 10.8 | 11.4 | 15.3 |
| **YK 059** | 47 | 45 | 38.9 | 22.8 | 3.62 | 13.1 | 13.6 | 11.4 | 25.5 | 4.09 | 8.21 | 10.6 | 2.55 | 3.5 | 5.75 | 13 | 7.97 | 9.64 | 17.2 | 2.3 | 11.2 | 11.2 | 15.6 |
| **YK 064** | 45.7 | 43 | 38.1 | 22.4 | 3.7 | 12.2 | 13.3 | 11.5 | 24.4 | 4.55 | 8.33 | 10.5 | 2.4 | 4.57 | 6.62 | 12.5 | 8.24 | 9.03 | 18.3 | 2.4 | 11.8 | 11.5 | |
| **YK 107** | 46.5 | 44.2 | 39.5 | 21.3 | 3.06 | 12.6 | 13.7 | 11.4 | 26.3 | 4.96 | 8.6 | 10.4 | 2.67 | 3.16 | 7.49 | 12.3 | 8.04 | 9.09 | 17.3 | 2.4 | 11.3 | 11.1 | 16 |
| **YK 112** | 49.2 | 46.1 | 40.3 | 23 | 2.75 | 12.4 | 14.2 | 12 | 27.4 | 4.51 | 8.89 | 11 | 2.74 | 3.26 | 7.43 | 13.4 | 8.23 | 9.85 | 18.6 | 2.32 | 11.2 | 11.3 | 16.6 |
| **YK 113** | 47.7 | 45.7 | 40.2 | 23 | 2.94 | 12.6 | 14.2 | 11.3 | 26 | 4.02 | 8.59 | 10.2 | 2.44 | 3.5 | 8.06 | 12.4 | 7.96 | 9.52 | 17.5 | 2.48 | 11.1 | 11 | 15.7 |
| **YK 129** | 47.4 | 44.9 | 39.7 | 23 | 2.88 | 11.9 | 13.6 | 11.6 | 27 | 3.62 | 8.64 | 10.1 | 2.55 | 3.2 | 7.5 | 12.5 | 8.12 | 9.4 | 17.9 | 2.4 | 10.6 | 11.1 | 15.3 |
| **YK 149** | 51.9 | 48.6 | 42.7 | 24.6 | 3.24 | 11.3 | 13.3 | 16.9 | 28.9 | 4.98 | 10.7 | 13.6 | 2.63 | 3.83 | 8.45 | 15.5 | 8.63 | 10.8 | 20.7 | 2.58 | 12.9 | 12.9 | 18 |
| **YK 200** | 47.1 | 45.2 | 38.9 | 22.4 | 3.32 | 12.2 | 13.6 | 11 | 25 | 3.75 | 7.7 | 10 | 2.46 | 2.98 | 7.15 | 12.9 | 7.97 | 9.86 | 18 | 2.59 | 11.2 | 11.7 | 15.2 |
| **YK 039** | 43.3 | 40.9 | 36.2 | 20.2 | 2.52 | 10.9 | 11.3 | 10.2 | 23.1 | 3.81 | 8.19 | 10.4 | 2.61 | 3.44 | 6.8 | 11 | 7.95 | 8.87 | 17.3 | 1.68 | 10.4 | 9.59 | |
| **EPL 008** | 46.3 | 43.6 | 37.2 | 22.1 | 3.42 | | | 11 | 24.6 | 4.74 | 7.94 | 10.2 | 2.7 | 3.33 | 5.83 | 12.8 | 8.18 | 9.45 | 16.1 | | 10.4 | 9.75 | 14 |
| **EPL 020** | 48.1 | 45.9 | 39.3 | 22.4 | 3.22 | 12.4 | 13.8 | 11.8 | 24.8 | 4.66 | 8.26 | 10.7 | 2.6 | 3.73 | 6.21 | 12.4 | 8.25 | 10.7 | 16.5 | 2.3 | 11.6 | 10.3 | 15.9 |
| **UM 040** | 48.6 | 45.6 | 39.5 | 22 | 3.67 | 11.7 | 13 | 12 | 26 | 4.6 | 8.52 | 10.2 | 2.38 | 3.2 | 6.87 | 12.8 | 8.3 | 10.2 | 16.3 | 2.06 | 12.1 | 11.2 | 15.5 |
| **Y081** | 43.6 | 40.7 | 35 | 20.6 | 3.22 | 10.9 | 11 | 10.1 | 22.9 | 4.08 | 7.19 | 9.98 | 2.71 | 2.01 | 6.44 | 10.8 | 7.13 | 9.19 | 15.7 | 2.21 | 11.2 | 10.4 | 14.8 |
| **YK 110** | 41.4 | 40.1 | 34 | 20.5 | 2.84 | 10.3 | 11.4 | 10 | 22.8 | 3.93 | 7.5 | 10.3 | 2.75 | 2.51 | 7.33 | 9.38 | 8.23 | 9.25 | 16.1 | 2.52 | 9.86 | 9.73 | 15.6 |
| **YK 128** | 41.4 | 38.7 | 34.1 | 18.8 | 2.6 | 9.86 | 11.3 | 10.3 | 22.6 | 3.29 | 7.71 | 9.73 | 2.4 | 2.21 | 6.93 | 10.7 | 7.86 | 8.55 | 15.3 | 2.2 | 9.7 | 9.01 | 15.3 |
| **YK 158** | 42.7 | 40.3 | 35.9 | 20.2 | 2.27 | 10.6 | 11.7 | 10.3 | 23 | 3.32 | 7.6 | 9.63 | 2.5 | 2.12 | 6.81 | 10.7 | 7.92 | 9.12 | 15 | 2.55 | 9.9 | 10.1 | 15.2 |
| **YK 210** | 41 | 39.1 | 33.7 | 18.7 | 2.39 | 9.31 | 10.3 | 10.1 | 22.6 | 3.38 | 7.22 | 9.35 | 2.5 | 2.72 | 5.7 | 10.5 | 7.3 | 8.59 | 14.7 | 2.33 | 9.21 | 9.41 | 14.6 |
| **Y033** | 41.3 | 39.7 | 34.6 | 19.4 | 2.84 | 10.1 | 11.5 | 10.2 | 23.5 | 3.65 | 7.14 | 9.81 | 2.56 | 2.34 | 6.22 | 10.4 | 7.08 | 8.94 | 15.3 | 2.37 | 10.3 | 10 | 14 |
| **Y042** | 41.4 | 39.4 | 34.8 | 19.1 | 2.83 | 10.2 | 12 | 10.3 | 22.8 | 3.65 | 7.2 | 9.04 | 2.49 | 2.02 | 7.12 | 11 | 7.35 | 9.47 | 15.1 | 2.04 | 11 | 10.5 | 14.5 |
| **YK 152** | 43.4 | 40.7 | 35.7 | 20.3 | 2.3 | 10.8 | 11.9 | 11 | 23.4 | 4.79 | 7.52 | 10.1 | 2.4 | 2.78 | 7.02 | 10.8 | 7.81 | 8.5 | 15.7 | 2.8 | 10.1 | 10.8 | 14.8 |
| **YK 190** | 41.1 | 39.3 | 33.7 | 18.8 | 2.66 | 9.86 | 11.5 | 10.4 | 21.5 | 3.02 | 6.75 | 8.83 | 2.29 | 2.2 | 6.1 | 10.7 | 6.98 | 8.7 | 14.5 | 2 | 9.45 | 9.4 | 15.2 |
| **YK 192** | 42.1 | 40.3 | 34.4 | 19.2 | 2.82 | 10.3 | 11.1 | 10.9 | 23 | 3.9 | 7.37 | 9.6 | 2.39 | 2.64 | 6 | 10 | 7.24 | 8.63 | 14.6 | 2.23 | 9.7 | 10.2 | 15.3 |
| **YK 195** | 42.6 | 40.5 | 34.8 | 19.4 | 2.75 | 9.66 | 10.9 | 10.2 | 23.2 | 3.88 | 7.37 | 9.38 | 2.12 | 2.85 | 6.54 | 10.9 | 7.7 | 9.74 | 14.8 | 2.3 | 10.3 | 9.8 | 15.6 |
| **UAC 901** | 45.6 | 43.1 | 36.8 | 21.2 | 3.24 | 11.3 | 13.9 | 11.7 | 24.7 | 4.28 | 8.18 | 10.3 | 2.66 | 3.55 | 6 | 12.4 | 8.19 | 9.23 | 16.4 | 2.7 | 9.52 | 9.93 | 14.7 |
| **UAC 913** | 44.6 | 43.1 | 36.2 | 21.3 | 2.8 | 11.9 | 13.5 | 11.5 | 24.8 | 4.15 | 7.95 | 10.6 | 2.45 | 3.64 | 6.43 | 11.8 | 8.5 | 9.07 | 15.9 | 2.43 | 10.2 | 10.4 | 14.5 |
| **EBO 50** | 48.1 | 45.9 | 39.8 | 23.1 | 3.2 | 12.8 | 13.5 | 12.1 | 27.2 | 4.26 | 8.78 | 10.8 | 2.88 | 3.4 | 7.5 | 12.9 | 8.2 | 9.55 | 15.6 | 2.1 | 11.2 | 10.6 | 17.1 |

Baek et al., submitted (Supplementary data)

| | | | | | | | | | | | | | | | | | | | | | | | |
|---|---|---|---|---|---|---|---|---|---|---|---|---|---|---|---|---|---|---|---|---|---|---|---|
| **EBO 230** | 46.5 | 44.4 | 39.1 | 22.6 | 3.08 | 12.9 | 14.2 | 12.2 | 26.3 | 3.6 | 8.4 | 10.6 | 2.86 | 3.08 | 7.33 | 12.5 | 7.89 | 9.59 | 15.9 | 2.22 | 11.5 | 11.1 | 16 |
| **EPL 025** | 46.7 | 44.6 | 38.9 | 22.7 | 2.82 | 11.5 | 13.2 | 11.3 | 26.4 | 3.72 | 8.67 | 10.7 | 2.65 | 3.7 | 6.7 | 12 | 8.75 | 9.19 | 16.2 | 2.56 | 10.4 | 10.4 | 15 |
| **EPL 031** | 45.8 | 43.2 | 36.9 | 21.7 | 2.94 | 11.1 | 12.3 | 12.1 | 24.7 | 3.62 | 8.3 | 10.2 | 2.59 | 3.75 | 7.1 | 12.4 | 8 | 9.14 | 15.7 | 2.98 | 10.1 | 9.62 | 14.8 |
| **EPL 041** | 46.7 | 44 | 38.5 | 22.1 | 3.46 | 10.9 | 12.5 | 12.1 | 26.2 | 3.33 | 8.46 | 10.5 | 2.76 | 3.82 | 6.31 | 12.6 | 8.35 | 9.15 | 15.7 | 2.26 | 11.1 | 10.3 | 16.1 |
| **EPL 045** | 46.1 | 43.5 | 37 | 21.2 | 3.38 | 11.8 | 13.1 | 11.5 | 24.9 | 3.97 | 7.77 | 10.1 | 2.44 | 3.8 | 6.36 | 11.8 | 7.75 | 9.19 | 16.2 | 2.12 | 10.4 | 10.2 | 14.7 |
| **EPL 046** | 46.8 | 44.3 | 38.5 | 22.1 | 3.59 | 11.6 | 13.7 | 12.1 | 25.8 | 4.42 | 8.4 | 11 | 2.7 | 4.3 | 7.22 | 13.4 | 8.54 | 8.77 | 16.4 | 2.22 | 11.1 | 10.9 | 16.1 |
| **EPL 047** | 44.9 | 41.4 | 36.4 | 20.8 | 2.94 | 11.1 | 12.7 | 10.9 | 24.6 | 3.52 | 7.3 | 10.1 | 2.26 | 3 | 6.27 | 11.5 | 8.25 | 9.5 | 16 | 2.35 | 10.2 | 9.89 | 14 |
| **EPL 048** | 45.1 | 42.6 | 38.5 | 21.7 | 3.4 | 10.7 | 12 | 11.9 | 24.8 | 3.98 | 8.92 | 10.4 | 2.8 | 3.78 | 6.62 | 12.2 | 8.5 | 9 | 15.7 | 2.29 | 10.5 | 10.1 | 15.3 |
| **EPL 049** | 44.7 | 42.3 | 37 | 20.8 | 3.33 | 11 | 12.5 | 11.8 | 25.1 | 4.35 | 8.35 | 10.3 | 2.41 | 3.67 | 6.72 | 12 | 7.62 | 10.1 | 15.9 | 2.07 | 11.2 | 10.1 | 14.7 |
| **EPL 052** | 45.9 | 43.8 | 37.7 | 21.8 | 3.53 | 11.6 | 13 | 12 | 26.8 | 4.11 | 8.43 | 10.6 | 2.57 | 3.15 | 6.4 | 13.1 | 8.14 | 8.7 | 16.7 | 2.55 | 10 | 11.1 | 15.2 |
| **EPL 053** | 46.7 | 44.4 | 38.8 | 22.5 | 3 | 11.9 | 12.9 | 11 | 26 | 4.29 | 9 | 10.5 | 2.51 | 3.42 | 6.78 | 12.5 | 8.7 | 9.4 | 16.1 | 2.7 | 10.7 | 10.9 | 15.7 |
| **UM 041** | 46.9 | 44.2 | 39.1 | 22.2 | 3.29 | 11.8 | 13.3 | 11.8 | 25.4 | 3.74 | 8.59 | 10.4 | 2.43 | 3.51 | 6.94 | 13.6 | 8.35 | 9.69 | 16.2 | 2.5 | 10.7 | 10.9 | 15.9 |
| **UM 383** | 46.8 | 43 | 39 | 21.8 | 3.1 | 11.3 | 12.5 | 11.8 | 24.9 | 3.46 | 8.9 | 10.4 | 2.43 | 3.17 | 6.7 | 13.1 | 8.91 | 9.02 | 15.8 | 2.14 | 11.1 | 10 | 15.7 |
| **AKT-106** | 45.1 | 43 | 37.2 | 21.4 | 3.01 | 11.2 | 12.2 | 12 | 25 | 3.88 | 8.4 | 10.1 | 2.66 | 4 | 6.16 | 12 | 7.98 | 9.22 | 15.5 | 2.41 | 9.8 | 10 | |
| **AKT-257** | 46.7 | 43.8 | 37.9 | 22.1 | 2.92 | 11.8 | 13.1 | 11.4 | 25.3 | 4.64 | 8.94 | 10.5 | 2.83 | 3.18 | 6.2 | 12.5 | 7.91 | 9.53 | 15.6 | 2.31 | 10.1 | 10.5 | 15.7 |
| **UAC 917** | 47.6 | 44.9 | 38.4 | 22.4 | 3.03 | 12.2 | 13.9 | 12.1 | 25.5 | 3.76 | 8.1 | 10.6 | 2.35 | 3.46 | 6.56 | 12.4 | 8.7 | 9.87 | 15.9 | 2.42 | 10.9 | 10.4 | 15.3 |
| **EBO 245** | 48.3 | 46 | 40.3 | 23.4 | 3.69 | 13.5 | 13.8 | 12.4 | 26.8 | 4.4 | 8.61 | 10.7 | 2.75 | 3.11 | 7.7 | 12.9 | 8.04 | 9.67 | 16 | 2.3 | 11.1 | 10.9 | 17.2 |
| **EBO 472** | 48 | 45.5 | 39.8 | 22.8 | 3.11 | 12.5 | 13.8 | 12.1 | 26.4 | 4.09 | 8.14 | 10.8 | 2.45 | 3.15 | 7.11 | 12.8 | 8.3 | 9.58 | 15.7 | 2.08 | 11.2 | 10.2 | 16.5 |
| **YHM337** | 45.2 | 42.9 | 37.1 | 21 | 3.03 | 11.7 | 12.3 | 12.3 | 23.2 | 3.62 | 8.07 | 9.86 | 2.44 | 2.79 | 6.49 | 12.1 | 8.6 | 9.59 | 14.9 | 2.42 | 10.8 | 9.68 | 15.3 |
| **YHM338** | 45.4 | 43.5 | 37.6 | 21.8 | 3.4 | 12.3 | 13.1 | 12.2 | 23.4 | 4.01 | 7.93 | 10.3 | 2.55 | 3.12 | 6.95 | 12.5 | 8.03 | 9.53 | 14.5 | 2.55 | 10.6 | 10.2 | 15.2 |
| **YK 036** | 45.8 | 42.9 | 38.6 | 21.9 | 3.2 | 11.9 | 13.7 | 11.5 | 25.8 | 4.59 | 8.38 | 10.4 | 2.44 | 3.1 | 6.01 | 12.7 | 8.25 | 9.7 | 16.7 | 2.11 | 10.5 | 10.1 | 14.9 |
| **YK 074** | 48.7 | 46.2 | 40.3 | 22.7 | 2.85 | 11.1 | 12.5 | 12.5 | 25.7 | 4.5 | 8 | 10.6 | 2.25 | 3 | 6.75 | 10.9 | 7.7 | 10.8 | 17.3 | 3.2 | 10.9 | 11.1 | 16.3 |
| **YK 151** | 50.4 | 46.8 | 42.3 | 23 | 3.4 | 12.1 | 13.7 | 13.1 | 27.8 | 4.86 | 8.85 | 10.6 | 2.56 | 3.16 | 7.85 | 14.5 | 8.79 | 9.48 | 18.8 | 2.42 | 11.5 | 11.4 | 16.9 |
| **M006** | 48.6 | 46 | 41.1 | 23.7 | 3.4 | 12.8 | 14.4 | 13 | 27.3 | 4.72 | 8.65 | 11.1 | 2.9 | 3.35 | 7.48 | 13.3 | 8.66 | 9.96 | 16.5 | 2.76 | 11.2 | 11.6 | 16.5 |
| **M008** | 48.3 | 45.4 | 39.2 | 22.7 | 4.1 | 11.9 | 13.4 | 13.2 | 25.6 | 4.75 | 8.55 | 10.8 | 2.67 | 3.7 | 7.1 | 12.6 | 8.26 | 9.94 | 17 | 2.49 | 11 | 10.3 | 15.4 |
| **M010** | 47.2 | 43.6 | 39.5 | 22.4 | 3.58 | 12.1 | 13.8 | 12.6 | 26.2 | 4.03 | 8.69 | 10.6 | 2.6 | 3.75 | 6.6 | 12.2 | 8.48 | 9.65 | 15.7 | 2.36 | 10.6 | 10.1 | 15.8 |
| **M032** | 45.9 | 43 | 37.8 | 21.9 | 3.33 | 11.6 | 12.4 | 11.7 | 25.3 | 3.94 | 7.54 | 9.62 | 2.43 | 3.63 | 6.25 | 12.7 | 7.65 | 9.29 | 16 | 2.55 | 10.4 | 10.1 | 14.5 |
| **M033** | 46.8 | 43.8 | 39.3 | 22 | 3.23 | 11.9 | 13.8 | 12.3 | 25.9 | 4.1 | 8.07 | 9.96 | 2.67 | 2.63 | 6.94 | 12.6 | 8.2 | 9.44 | 16.6 | 2.3 | 11 | 10.1 | 14.9 |
| **EPL 032** | 48.2 | 45.4 | 38.8 | 22.2 | 3.25 | 11.7 | 13.1 | 11.7 | 26.1 | 4.52 | 8.42 | 10.4 | 2.43 | 3.36 | 6.22 | 13.6 | 8.25 | 10.1 | 16.9 | 2.19 | 10.8 | 10.2 | 15.9 |
| **EPL 034** | 46.4 | 44.2 | 38.6 | 22.3 | 3.47 | 11.8 | 12.7 | 12.4 | 25.4 | 4.52 | 8.6 | 10.7 | 2.83 | 3.72 | 6.41 | 11.6 | 8.42 | 9.23 | 16.1 | 2.65 | 10.3 | 10.3 | 15.1 |
| **EPL 042** | 46.9 | 44.7 | 37.9 | 22.2 | 2.86 | 11.6 | 12.9 | 12.3 | 25.6 | 4.1 | 8 | 10.6 | 2.83 | 3.4 | 6.25 | 12.1 | 8.71 | 9.37 | 16.3 | 2.32 | 10.2 | 9.7 | 15.7 |

| | | | | | | | | | | | | | | | | | | | | | | | |
|---|---|---|---|---|---|---|---|---|---|---|---|---|---|---|---|---|---|---|---|---|---|---|---|
| **EPL 043** | 49.6 | 46.6 | 40.2 | 23.3 | 3.2 | 12.4 | 13.9 | 12.4 | 26.6 | 4.04 | 8.73 | 11.1 | 2.58 | 3.09 | 7.32 | 14.3 | 9.2 | 10.3 | 16.4 | 2.42 | 11.5 | 11.1 | 16.9 |
| **EPL 051** | 48.6 | 45.3 | 39.3 | 22.6 | 3.07 | 12.2 | 12.7 | 12.2 | 25.9 | 4.34 | 8.6 | 10.4 | 2.45 | 4 | 7.2 | 13 | 8.54 | 10.8 | 16.4 | 2.49 | 11.4 | 11.3 | 15.3 |
| **UM 019** | 49.5 | 46.9 | 40.6 | 23 | 3.65 | 12.7 | 14.2 | 11.9 | 27 | 4.3 | 8.2 | 10.4 | 2.5 | 3.56 | 7.01 | 14.5 | 8.01 | 10.4 | 17.2 | 2.45 | 12.8 | 11.2 | 16.1 |
| **UM 384** | 46.8 | 44.3 | 38.5 | 22.2 | 2.95 | 11.8 | 12.9 | 12.3 | 25 | 4.28 | 8.76 | 10.5 | 2.37 | 3.13 | 6.45 | 13.1 | 8.47 | 9.94 | 15.7 | 2.17 | 10.6 | 9.98 | 14.8 |
| **UM 440** | 46.5 | 44 | 37.9 | 22.1 | 3.56 | 11.2 | 13.9 | 11.9 | 25.1 | 4.6 | 8.09 | 10.1 | 2.43 | 3.81 | 7.4 | 13.4 | 8.02 | 10.1 | 16.8 | 2.88 | 11.9 | 10.1 | 14.8 |
| **UM 510** | 49.9 | 45.6 | 40.2 | 22.9 | 4 | 11.6 | 13 | 12.3 | 27.3 | 4.37 | 8.6 | 9.98 | 2.35 | 4.18 | 7.3 | 14.2 | 8.03 | 10.3 | 16.7 | 2.48 | 11.7 | 10.9 | 15.8 |
| **UAC 919** | 44.6 | 43 | 36.4 | 20.3 | 2.73 | 11.2 | 11.7 | 10.2 | 25.3 | 3.76 | 7.53 | 9.72 | 2.18 | 3.24 | 6.15 | 11.5 | 8.13 | 9.06 | 16.2 | 2.12 | 9.93 | 10 | 14.5 |
| **UAC 922** | 46 | 43.7 | 37.7 | 21.9 | 3.44 | 11.8 | 13.2 | 12.2 | 25.6 | 4.69 | 8.5 | 10.6 | 2.33 | 3.73 | 6.22 | 12.5 | 8.02 | 9.72 | 16.8 | 2.52 | 11.4 | 10.7 | 14.9 |
| **UAC 934** | 44.9 | 42.9 | 38.1 | 21.9 | 3.4 | 11.1 | 13.1 | 11 | 24.9 | 4.2 | 7.3 | 10 | 2.5 | 3 | 6.6 | 12.5 | 8.6 | 8.4 | 15.3 | 2.32 | 10 | 10 | 15 |
| **MBIYE 525** | 46.5 | 44.2 | 38.9 | 22.4 | 3.34 | 11.9 | 13.1 | 11.5 | 26.2 | 4.52 | 8.37 | 11.1 | 2.76 | 3.95 | 6.96 | 12.4 | 8.25 | 9.34 | 16.1 | 2.27 | 11.1 | 10.9 | 15.5 |
| **MBIYE 526** | 45.8 | 43.6 | 38.2 | 22.1 | 3.06 | 11.8 | 12.9 | 11.8 | 26.9 | 5 | 8.34 | 11.1 | 3 | 3.5 | 6.93 | 12.5 | 8.74 | 9.4 | 15.9 | 2.62 | 10.7 | 10.2 | 16 |
| **MBIYE 528** | 45.8 | 43 | 38.9 | 22.2 | 3.12 | 11.5 | 12.7 | 11.8 | 26.9 | 4.98 | 8.74 | 10.8 | 2.8 | 3.4 | 7 | 13.1 | 8.93 | 9.22 | 16 | 2.5 | 11.1 | 10.4 | 16 |
| **BDA 80** | 46.3 | 44.8 | 38.6 | 22.4 | 3.75 | 12.1 | 13 | 11.2 | 25.5 | 4.1 | 8.8 | 10.7 | 2.73 | 2.65 | 6.9 | 12.5 | 8.47 | 9.47 | 15.4 | 3 | 11.5 | 10.3 | 15.7 |
| **BLI01** | 46.6 | 44.2 | 38.2 | 22.4 | 3.25 | 11.7 | 13.1 | 12.5 | 26.3 | 3.9 | 8.21 | 11 | 2.85 | 3.66 | 6.93 | 12.2 | 8.4 | 9.12 | 16.2 | 2.47 | 11.3 | 10.8 | 17.1 |
| **Y021** | 41.3 | 38.6 | 34.2 | 20.2 | 2.65 | 9.91 | 10.8 | 10.3 | 23.4 | 4.25 | 7.7 | 8.82 | 2.63 | 1.92 | 6.65 | 10.5 | 7.58 | 9.06 | 16.4 | 2.12 | 10.2 | 10.8 | 15.4 |
| **YK 070** | 41.1 | 38.6 | 34.3 | 18.6 | 2.6 | 11.4 | 12.5 | 9.95 | | 3.61 | 7.35 | 9.06 | 2.22 | 3.83 | 6.21 | 10.5 | 7.43 | 9.07 | 16.1 | 2.36 | 10 | 9.05 | 14.1 |
| **YK 083** | 46.5 | 43.6 | 39.8 | 22.1 | 3.34 | 12.2 | 13.3 | 11.8 | 25.8 | 4.43 | 8.37 | 10.3 | 2.29 | 3.96 | 7.67 | 13.1 | 7.73 | 8.86 | 18 | 2.06 | 10.9 | 11.7 | 15.8 |
| **YK 193** | 47.8 | 46 | 39.4 | 22.4 | 3.19 | 12 | 13.7 | 11.2 | 25.2 | 3.79 | 8.54 | 10.6 | 2.81 | 2.44 | 7.18 | 11.5 | 7.93 | 9.1 | 15.1 | 2.43 | 10.8 | 10.4 | 15.7 |
| **YK 211** | 41.3 | 38.8 | 33 | 19.3 | 2.28 | 9.7 | 10.8 | 10.3 | 22.1 | 3 | 7.12 | 9.11 | 2.18 | 2.9 | 5.63 | 10.4 | 7.44 | 9.36 | 15 | 2.14 | 9.4 | 8.93 | 13.8 |
| **EPL 023** | 45.7 | 43.6 | 38 | 22.1 | 2.82 | 11.3 | 13.3 | 11.1 | 26 | 4 | 9.08 | 10.6 | 2.6 | 4.24 | 6.26 | 12.1 | 8.22 | 9.54 | 16.6 | 2.31 | 10.3 | 10.9 | 15.3 |
| **EPL 044** | 47 | 44.6 | 39.3 | 22 | 3.32 | 11.8 | 13.3 | 11.1 | 25.7 | 4.22 | 7.95 | 10.8 | 2.49 | 3.62 | 6.67 | 12 | 8.15 | 10.2 | 16 | 2.52 | 11.6 | 10.1 | 15.8 |
| **EPL 054** | 49.9 | 46 | 40.7 | 22.5 | 3.5 | 11.9 | 13.3 | 12.3 | 25.3 | 4.25 | 8.42 | 10.7 | 2.5 | 2.5 | 6.6 | 12.9 | 8.25 | 11 | 17 | 2.45 | 12.2 | 11.5 | |
| **RBTL 348** | 47 | 44.4 | 37.3 | 21.4 | 3.23 | 11.2 | 12.3 | 11 | 24.5 | 3.98 | 8.54 | 9.98 | 2.3 | 2.87 | 6.49 | 12 | 8.08 | 9.41 | 15.8 | 2.52 | 10.6 | 9.88 | 14.9 |
| **RBTL 476** | 46.4 | 44.2 | 38.1 | 22.1 | 3.22 | 11.9 | 13 | 11.2 | 25.5 | 4.54 | 8.14 | 10.6 | 2.49 | 3.62 | 6.65 | 11.9 | 8.03 | 9.13 | 14.9 | 2.63 | 10.7 | 10 | 14.9 |
| **UM 386** | 45.5 | 42.1 | 37.1 | 21.2 | 3.15 | 10.5 | 12.4 | 11.8 | 25.7 | 3.87 | 8.35 | 10.5 | 2.72 | 3.6 | 6.54 | 12.7 | 8.9 | 8.98 | 16.5 | 2.6 | 10.5 | 9.85 | 15.8 |
| **RMCA1** | 43.5 | 40 | 35.4 | 20.3 | 3.2 | 11.9 | 12.3 | 11.2 | 24 | 3.9 | 7.6 | 11.2 | 2.7 | 3.7 | 6.4 | 12.5 | 8.2 | 8.8 | 15.2 | 2.3 | 10.1 | 10.9 | 14.5 |
| **RMCA2** | 44 | 41 | 36.3 | 20.3 | 3.4 | 11.5 | 13.3 | | | 4.2 | 7.5 | 11 | 2.6 | 4.4 | 7 | 11.9 | 7.6 | 9.8 | | 2.2 | 11 | 9.9 | 15 |
| **RMCA3** | 43 | 40.6 | 35.6 | 20.4 | 3.2 | 11.4 | 13.2 | 10 | 12.8 | 4.3 | 7.4 | 10.8 | 2.4 | 4.3 | 6.5 | 11.3 | 7.5 | 9.5 | 15.5 | 2.2 | 11 | 10.1 | 14.8 |
| **RMCA 17346-M-** | 41.2 | 38.8 | 33.8 | 19.4 | 3.2 | 10.8 | 12.4 | 10.1 | | 4.1 | 7.2 | 10.3 | 2.5 | 4.1 | 6.5 | 11.2 | 7.5 | | | 2.1 | 10.3 | 9.5 | 14.4 |

Baelo et al., Submitted (Supplementary data)

| | | | | | | | | | | | | | | | | | | | | | | | |
|---|---|---|---|---|---|---|---|---|---|---|---|---|---|---|---|---|---|---|---|---|---|---|---|
| **RMCA 14053-M-** | 41.2 | 38.3 | 33.9 | 19.3 | 3.1 | 10.9 | 12.5 | 10.3 | 22.7 | 4 | 7.1 | 10.4 | 2.3 | 3.9 | 6.2 | 11.1 | 7.1 | 8.9 | 15 | 2.1 | 10.3 | 9.8 | 13.8 |
| **RMCA 17354-M-** | 40.5 | 37.9 | 33.2 | 19.2 | 3.1 | 10.7 | 12.3 | 10.1 | | 3.9 | 7.1 | 10.2 | 2.4 | 3.9 | 6.4 | 10.8 | | 8.7 | 14.4 | 2.1 | 10.2 | 10.1 | |
| **RMCA 17349-M-** | 40.5 | 38.1 | 33 | 19.1 | 3 | 10.6 | 12.2 | 10.2 | 22.4 | 3.9 | 7 | 10.2 | 2.4 | 4 | 6.4 | 10.9 | 7.5 | 8.8 | 14.4 | 2.2 | 10 | 10 | 13.9 |
| **RMCA 17800-M-** | 40.7 | | | | 3 | 10.7 | 12.2 | 10.2 | 22.1 | 3.9 | | 10.3 | 2.4 | 4 | 6.2 | 10.9 | 7.2 | | 14.8 | 2.2 | 10.2 | 9.2 | 13.9 |
| **RMCA 17355-M-** | 37.9 | 36.3 | 30.6 | 16.7 | 3.2 | 8.8 | 9.8 | 9.7 | | 3.8 | 7.4 | 8.8 | 2.6 | 3.2 | 6.2 | 9.3 | 7.3 | 9.2 | 14.5 | 2.1 | 9.4 | 9.2 | |
| **RMCA 17348-M-** | 37.3 | 36 | 30.3 | 16.8 | 2.9 | | | 9.5 | | 4.2 | 7.5 | 9 | 2.3 | 3.2 | 5.7 | 8.8 | 6.9 | | 14.7 | 2.1 | 8.9 | 8.8 | 13.8 |
| **RMCA 17799-M-** | | | | | 2.8 | 9.8 | 11.1 | | | 4.1 | | | 2.6 | 3.3 | 6 | 12.6 | 6.4 | | | 2.1 | 9.2 | 8.5 | 14.4 |
| **RMCA 17350-M-** | | 39.1 | 34.1 | 17.6 | 3.2 | 10.7 | 12.2 | 10.4 | 22.5 | 4.5 | 6.9 | 9.6 | 2.75 | 3.6 | 6.7 | | 7.2 | 8.9 | 13.9 | 2.4 | 10.5 | 9.3 | 13.9 |
| **RMCA 14054-M-** | | | | 18.1 | 2.8 | 8.9 | 10.2 | 10.6 | 22.7 | 4.2 | 7.4 | 9.6 | 2.6 | 3.6 | 6.2 | 8.9 | 7.3 | | 14.6 | 2.2 | 10.5 | 9.4 | 15 |
| **RMCA 17330-M-** | | | | | 2.5 | 9.5 | 10.2 | 10.4 | | 3.9 | 7.4 | 9.3 | 2.65 | 3.4 | 6 | 10.4 | 7.6 | | | 2.1 | 9.1 | 8.2 | 13.3 |
| **RMCA 17802-M-** | | | | | 2.5 | 8.3 | 10.5 | 9.5 | | 4 | | 8.5 | 2.2 | 4 | 5.8 | 9.6 | 6.8 | | | 2.1 | 9 | 8.5 | |
| **RMCA 17353-M-** | | | | 17.2 | 3.2 | 8.7 | 10 | 10.3 | 22.2 | 4.4 | 7.1 | 8.8 | 2.3 | 3.4 | 5.7 | | 8.2 | | | 2.1 | 10.2 | 9 | 14.6 |
| **RMCA 17992-M-** | | | | | | | | | 22.6 | | 7.9 | 8.4 | 2.3 | | | | 8 | | 15.5 | | | | |
| **RMCA 720A-M-** | | | | | 2.5 | | | | | | 7.3 | | 2.4 | | | | 6.8 | | | | | 8.5 | 13.4 |
| **RMCA 720B-M-** | | | | 17.1 | 2.8 | 9.8 | 11.2 | 10 | | 3.5 | 7.2 | 9 | 2.2 | 3.3 | 5.5 | | 7.3 | | 13.7 | 2.2 | 10.4 | 8 | 14.5 |
| **RMCA 17351-M-** | | | | | 2.8 | 9.7 | 10.2 | 9.6 | | 3.8 | 7.2 | 8.8 | 2.2 | | 5.6 | | 7.8 | | | 2.1 | 9.2 | 7.3 | 14.6 |
| **RMCA 31116-M-** | | | | | | | | | | | | | | | 5.9 | 10.1 | 7.6 | | | 2.3 | | 9.8 | 13.7 |

Baelo et al., submitted (Supplementary data)

| | | | | | | | | | | | | | | | | | | | | | | | |
|---|---|---|---|---|---|---|---|---|---|---|---|---|---|---|---|---|---|---|---|---|---|---|---|
| **RMCA 17352-M-** | | | | | 3.2 | 9.2 | 10.5 | 10.2 | | 4.4 | 7.3 | 9 | 2.7 | 3.1 | 5.6 | 9.3 | 7.2 | 8.1 | | 2.3 | 10 | 8.6 | 13.8 |
| **RMCA 9563-M-** | | | | | | | | | | | | | | | | | | | 2.2 | | | | |
| **RMCA 17652-M-** | 39.3 | 37.8 | | | 3.5 | | | | | | | | | | | 10 | 8.1 | | | 2.2 | | | 14.1 |
| **RMCA 17991-M-** | | | | 19.6 | 2.99 | 9.6 | 11 | 11.7 | 22.6 | 4.2 | 7.1 | 9.4 | 2.6 | 3.7 | 6.6 | 11.4 | 7.5 | 8.9 | 15.2 | 2.3 | 10.1 | 10.3 | 14.4 |
| **RMCA 31115-M-** | | | | | | | | | | | | | | | | | | | 2.2 | | | | |
| **RMCA 123-M-** | 39.3 | 37.8 | | | 3.5 | | | | | | | | | | | 10 | 8.1 | | | 2.2 | | | 14.1 |
| **RMCA 31117-M-** | | | | 19.6 | 2.99 | 9.6 | 11 | 11.7 | 22.6 | 4.2 | 7.1 | 9.4 | 2.6 | 3.7 | 6.6 | 11.4 | 7.5 | 8.9 | 15.2 | 2.3 | 10.1 | 10.3 | 14.4 |

Baelo et al., submitted (Sypplementary data)

**Table S3.** Sexual dimorphism based on cranial measurements of the species sampled from the left and right banks of the Congo River (ns= non-significant difference and * significant).

| Measure | N (M) | Mean (M) | SD (M) | N (F) | Mean (F) | SD (F) | t-value |
|---|---|---|---|---|---|---|---|
| | | | *F. anerythrus from the right bank* | | | | |
| M1 | 30 | 46.53 | 1.1583 | 30 | 46.51 | 1.5757 | 0.049 |
| M2 | 30 | 44.15 | 1.1119 | 30 | 43.77 | 1.3854 | 1.159 |
| M3 | 30 | 38.23 | 0.9861 | 30 | 38.3 | 1.266 | -0.251 |
| M4 | 30 | 22.05 | 0.5392 | 30 | 21.95 | 0.7579 | 0.601 |
| M5 | 30 | 3.17 | 0.2585 | 30 | 3.31 | 0.3586 | -1.66 |
| M6 | 30 | 11.74 | 0.5102 | 29 | 11.46 | 0.5571 | 1.97 |
| M7 | 30 | 13.11 | 0.6098 | 29 | 13.01 | 0.6287 | 0.599 |
| M8 | 30 | 11.68 | 0.5241 | 30 | 11.74 | 0.711 | -0.325 |
| M9 | 29 | 25.43 | 0.5875 | 30 | 25.63 | 0.8443 | -1.069 |
| M10 | 30 | 4.2 | 0.3169 | 30 | 4.18 | 0.4229 | 0.294 |
| M11 | 30 | 8.29 | 0.3091 | 30 | 8.37 | 0.4525 | -0.776 |
| M12 | 30 | 10.39 | 0.3089 | 30 | 10.44 | 0.323 | -0.584 |
| M13 | 30 | 2.55 | 0.1983 | 30 | 2.57 | 0.1635 | -0.412 |
| M14 | 30 | 3.45 | 0.3865 | 30 | 3.53 | 0.4239 | -0.773 |
| M15 | 30 | 6.81 | 0.4809 | 30 | 6.64 | 0.4386 | 1.422 |
| M16 | 30 | 12.49 | 0.6699 | 30 | 12.57 | 0.7272 | -0.48 |
| M17 | 30 | 8.28 | 0.3396 | 30 | 8.36 | 0.3381 | -0.926 |
| M18 | 30 | 9.46 | 0.6727 | 30 | 9.39 | 0.5598 | 0.394 |
| *M19* | *30* | *16.05* | *0.46* | *30* | *16.3* | *0.382* | *-2.314* |
| M20 | 30 | 2.48 | 0.2203 | 29 | 2.41 | 0.1911 | 1.332 |
| M21 | 30 | 10.81 | 0.7105 | 30 | 10.71 | 0.5791 | 0.596 |
| M22 | 30 | 10.35 | 0.4396 | 30 | 10.41 | 0.4954 | -0.458 |
| M23 | 29 | 15.41 | 0.6504 | 29 | 15.34 | 0.6761 | 0.402 |
| | | | *F. anerythrus from the left bank* | | | | |
| M1 | 48 | 47.25 | 1.495 | 35 | 47.463 | 1.4857 | 0.653 |
| M2 | 47 | 44.42 | 1.3655 | 35 | 44.734 | 1.3472 | 1.021 |
| M3 | 47 | 39.2 | 1.2955 | 35 | 39.482 | 1.2086 | 0.995 |
| M4 | 47 | 22.43 | 0.7251 | 35 | 22.514 | 0.7633 | 0.501 |
| M5 | 46 | 3.324 | 0.2643 | 35 | 3.2491 | 0.328 | -1.139 |
| M6 | 48 | 12.17 | 0.4978 | 35 | 12.265 | 0.5888 | 0.805 |
| M7 | 48 | 13.61 | 0.6041 | 35 | 13.795 | 0.6071 | 1.357 |
| M8 | 47 | 11.71 | 0.5985 | 35 | 11.864 | 0.5426 | 1.189 |
| M9 | 40 | 25.81 | 1.0496 | 31 | 26.156 | 0.8939 | 1.474 |
| M10 | 47 | 4.143 | 0.4285 | 35 | 4.2023 | 0.4124 | 0.628 |
| M11 | 47 | 8.431 | 0.4481 | 35 | 8.3989 | 0.3209 | -0.359 |
| M12 | 47 | 10.59 | 0.5386 | 35 | 10.546 | 0.4273 | -0.409 |
| M13 | 47 | 2.491 | 0.1516 | 35 | 2.492 | 0.2042 | 0.024 |
| M14 | 46 | 3.56 | 0.5423 | 35 | 3.448 | 0.3926 | -1.033 |
| M15 | 47 | 6.957 | 0.4463 | 35 | 6.9649 | 0.564 | 0.066 |
| M16 | 47 | 12.81 | 0.5872 | 35 | 12.757 | 0.7959 | -0.332 |
| M18 | 48 | 9.555 | 0.357 | 35 | 9.5251 | 0.4103 | -0.358 |

Baelo et al. submitted (Supplementary data)

| | | | | | | | |
|---|---|---|---|---|---|---|---|
| M19 | 48 | 17.91 | 0.9431 | 35 | 17.61 | 0.8825 | -1.463 |
| M20 | 47 | 2.343 | 0.187 | 35 | 2.3523 | 0.2238 | 0.205 |
| M21 | 47 | 11.41 | 0.5685 | 35 | 11.387 | 0.4545 | -0.168 |
| M22 | 47 | 11.07 | 0.5258 | 35 | 11.166 | 0.489 | 0.854 |
| M23 | 42 | 15.59 | 0.6288 | 33 | 15.841 | 0.7763 | 1.526 |
| | | | *F. cf bayonii* | | | | |
| M1 | 25 | 42.28 | 0.8982 | 11 | 42.172 | 0.919 | -0.32 |
| M2 | 25 | 39.9 | 0.7644 | 11 | 40.244 | 0.6696 | 1.305 |
| M3 | 25 | 34.66 | 0.7813 | 11 | 34.953 | 0.9001 | 0.975 |
| M4 | 25 | 19.67 | 0.659 | 11 | 19.93 | 0.4703 | 1.186 |
| M5 | 25 | 2.572 | 0.2678 | 11 | 2.6055 | 0.3769 | 0.304 |
| M6 | 25 | 10.09 | 0.4716 | 11 | 10.067 | 0.5309 | -0.126 |
| M7 | 25 | 11.33 | 0.4475 | 11 | 11.446 | 0.5117 | 0.693 |
| M8 | 25 | 10.43 | 0.3671 | 11 | 10.627 | 0.4798 | 1.329 |
| M9 | 25 | 23.04 | 0.5311 | 11 | 23.14 | 0.7857 | 0.437 |
| M10 | 25 | 3.71 | 0.4772 | 11 | 3.8245 | 0.5078 | 0.653 |
| *M11* | *25* | *7.48* | *0.32* | *11* | *7.776* | *0.376* | *2.426* |
| *M12* | *25* | *9.55* | *0.372* | *11* | *9.912* | *0.527* | *2.361* |
| M13 | 25 | 2.424 | 0.1814 | 11 | 2.5009 | 0.2117 | 1.108 |
| *M14* | *25* | *2.5* | *0.511* | *11* | *2.966* | *0.484* | *2.552* |
| M15 | 25 | 6.394 | 0.4322 | 11 | 6.7845 | 0.5613 | 2.28 |
| M16 | 25 | 10.58 | 0.512 | 11 | 10.239 | 0.8177 | -1.532 |
| M17 | 25 | 7.586 | 0.3567 | 11 | 7.6918 | 0.4014 | 0.79 |
| M18 | 25 | 8.986 | 0.2943 | 11 | 9.1391 | 0.2243 | 1.54 |
| *M19* | *25* | *15.3* | *0.606* | *11* | *15.8* | *0.985* | *2.045* |
| M20 | 25 | 2.344 | 0.2325 | 11 | 2.41 | 0.4191 | 0.612 |
| M21 | 25 | 10.2 | 0.6339 | 11 | 10.176 | 0.5466 | -0.12 |
| M22 | 25 | 9.939 | 0.6288 | 11 | 10.017 | 0.6628 | 0.338 |
| M23 | 23 | 14.99 | 0.5583 | 9 | 15.361 | 0.2276 | 1.935 |

**Table S4.** Sexual dimorphism based on externals measurements (LF, Length of hind foot; LE, Length of ear; LT, Length of tail; LB, Length of body; ns= non-significant difference and * significant).

| Measure | N (M) | Mean (M) | SD (M) | N (F) | Mean (F) | SD (F) | t-value | df | p | Sign. |
|---|---|---|---|---|---|---|---|---|---|---|
| *F. anerythrus from the right bank* | | | | | | | | | | |
| **Poids** | 50 | 182.29 | 21 | 52 | 185.78 | 28.64 | 0.3299 | 55.703 | 0.7427 | ns |
| **LP** | 51 | 39.95 | 6.2 | 53 | 40.27 | 4.79 | 1.1979. | 49.722 | 0.2366 | ns |
| **LO** | 50 | 16.81 | 2.35 | 48 | 16.1 | 2.01 | -0.47253 | 55.95 | 0.6384 | ns |
| **LQ** | 48 | 144.41 | 26.2 | 53 | 152.92 | 14.76 | 0.91583 | 57.367 | 0.3636 | ns |
| **LC** | 51 | 187.05 | 27.51 | 53 | 182.37 | 19.13 | -1.4207 | 56.876 | 0.1609 | ns |
| *F. anerythrus from the left bank* | | | | | | | | | | |
| **Poids** | 71 | 186.36 | 24.82 | 44 | 208.45 | 19.24 | 3.7623 | 53.642 | 0.0004 | * |
| **LP** | 71 | 39.39 | 4.74 | 44 | 40.3 | 4.28 | -0.97509 | 56.484 | 0.3337 | ns |
| **LO** | 66 | 17.1 | 1.34 | 41 | 17.15 | 1.38 | 0.0373 | 57.956 | 0.9703 | ns |
| **LQ** | 58 | 153.81 | 13.24 | 40 | 154.02 | 10.48 | 0.35311 | 55.02 | 0.7254 | ns |
| **LC** | 71 | 189.81 | 20.72 | 44 | 186.5 | 15.88 | -1.8154 | 57.945 | 0.0746 | ns |
| *F. cf bayonii* | | | | | | | | | | |
| **Poids** | 39 | 133.41 | 12.32 | 21 | 136.9 | 16.15 | 0.22136 | 38.194 | 0.826 | ns |
| **LP** | 39 | 37.93 | 2.69 | 21 | 37.32 | 2.78 | -0.85628 | 37.686 | 0.3973 | ns |
| **LO** | 37 | 15.12 | 2.12 | 21 | 15.21 | 1.42 | -0.40261 | 30.705 | 0.69 | ns |
| **LQ** | 39 | 167.33 | 17.09 | 20 | 167.5 | 19.43 | 1.3811 | 33.507 | 0.1764 | ns |
| **LC** | 39 | 173.2 | 22.38 | 20 | 171 | 18.15 | -0.35662 | 39.227 | 0.7233 | ns |

Submitted (Supplementary data)

**Table S5.** External morphometric characteristics of the species sampled from the left (LB) and right (RB) banks of the Congo River (weight in grams, the length in millimeters, N, number of individuals; SD, standard deviation and CV, coefficient of variation).

| Species | Parameter | Weight | Length of hind foot | Length of ear | Length of tail | Length of body |
|---|---|---|---|---|---|---|
| | N | 92 | 94 | 88 | 68 | 94 |
| | Minimum | 123.7 | 14.23 | 11.81 | 126 | 140 |
| | Maximum | 287 | 46.08 | 20.5 | 188 | 300 |
| *F. anerythrus*_RB | Mean | *185.12* | *40.16* | *16.35* | *160.97* | *186.26* |
| | Variance | 907.59 | 32.33 | 3.82 | 162.6 | 476.45 |
| | SD | 30.13 | 5.69 | 1.95 | 12.75 | 21.83 |
| | CV | 0.16 | 0.14 | 0.11 | 0.07 | 0.11 |
| | N | 82 | 83 | 78 | 82 | 82 |
| | Minimum | 110 | 26.95 | 13.1 | 122 | 154 |
| | Maximum | 284 | 46.33 | 20.09 | 188 | 208 |
| *F. anerythrus*_LB | Mean | *196.14* | *39.35* | *17.07* | *160.34* | *182.16* |
| | Variance | 745.99 | 23.24 | 1.68 | 89.93 | 116.53 |
| | SD | 27.31 | 4.82 | 1.3 | 9.48 | 10.79 |
| | CV | 0.13 | 0.12 | 0.07 | 0.05 | 0.05 |
| | N | 60 | 60 | 58 | 47 | 59 |
| | Minimum | 109 | 26.6 | 10.05 | 156 | 131 |
| | Maximum | 179 | 49.96 | 18.1 | 178 | 208 |
| *F. cf bayonii* | Mean | 134.63 | 37.73 | 15.15 | 166.74 | 177.36 |
| | Variance | 189.15 | 7.4 | 3.41 | 32.67 | 167.13 |
| | SD | 13.75 | 2.72 | 1.85 | 5.72 | 12.93 |
| | CV | 0.1 | 0.07 | 0.12 | 0.03 | 0.07 |

In general, no external biometric measurements of *F. anerythrus* were stable on either bank of the Congo River. However, for *F. anerythrus* on the left bank, weight and hind foot length showed moderate variability (0.1 ≤ CV < 0.2), and three other measures - ear length, hind foot length and body length - were relatively unstable (0.05 ≤ CV < 0.1). In contrast, for *F. anerythrus* on the right bank, only hind foot length showed moderate instability, while weight, ear length and body length showed considerable variability (0.1 ≤ CV < 0.2). The results of the descriptive statistics analysis for *F. anerythrus* specimens from both banks

Baelo et al., submitted (Supplementary data)

indicated a significant difference in weight ($t = 3.0852$, $df = 51.621$, p-value = $0.003265 < 0.05$), with individuals from the left bank having a slightly higher mean weight (196.14±27.31) compared to those from the right bank (185.12±30.131). However, the analysis did not reveal significant differences in other variables, including hind foot length.

**Table S6**. Cranial morphometric characteristics of the species sampled from the left and right banks of the Congo River (N, number of individuals)

| | N | Minimum | Maximum | Mean | Variance | Standard deviation | Coefficient of variation |
|---|---|---|---|---|---|---|---|
| *F. anerythrus from the right bank* | | | | | | | |
| M1 | 60 | 43.93 | 49.94 | 46.52 | 1.88 | 1.37 | 0.03 |
| M2 | 60 | 41.2 | 46.9 | 43.96 | 1.59 | 1.26 | 0.03 |
| M3 | 60 | 36.22 | 41.12 | 38.27 | 1.27 | 1.13 | 0.03 |
| M4 | 60 | 20.29 | 23.72 | 22 | 0.43 | 0.65 | 0.03 |
| M5 | 60 | 2.4 | 4.1 | 3.24 | 0.1 | 0.32 | 0.1 |
| M6 | 59 | 10.34 | 12.84 | 11.6 | 0.3 | 0.55 | 0.05 |
| M7 | 59 | 11.66 | 14.41 | 13.06 | 0.38 | 0.62 | 0.05 |
| M8 | 60 | 10.16 | 13.19 | 11.71 | 0.38 | 0.62 | 0.05 |
| M9 | 59 | 23.97 | 27.27 | 25.53 | 0.53 | 0.73 | 0.03 |
| M10 | 60 | 3.33 | 4.99 | 4.19 | 0.14 | 0.37 | 0.09 |
| M11 | 60 | 7.3 | 9.08 | 8.33 | 0.15 | 0.39 | 0.05 |
| M12 | 60 | 9.62 | 11.08 | 10.42 | 0.1 | 0.31 | 0.03 |
| M13 | 60 | 2.15 | 2.9 | 2.56 | 0.03 | 0.18 | 0.07 |
| M14 | 60 | 2.5 | 4.3 | 3.49 | 0.16 | 0.4 | 0.12 |
| M15 | 60 | 5.83 | 7.62 | 6.72 | 0.22 | 0.46 | 0.07 |
| M16 | 60 | 11.48 | 14.45 | 12.53 | 0.48 | 0.69 | 0.06 |
| M17 | 60 | 7.62 | 9.26 | 8.32 | 0.11 | 0.34 | 0.04 |
| M18 | 60 | 8 | 10.98 | 9.42 | 0.38 | 0.61 | 0.07 |
| M19 | 60 | 14.94 | 17.2 | 16.18 | 0.19 | 0.44 | 0.03 |
| M20 | 59 | 2.06 | 3 | 2.45 | 0.04 | 0.21 | 0.08 |
| M21 | 60 | 9.52 | 12.78 | 10.76 | 0.42 | 0.64 | 0.06 |
| M22 | 60 | 9.62 | 11.59 | 10.38 | 0.22 | 0.47 | 0.04 |
| M23 | 58 | 13.98 | 17.1 | 15.37 | 0.43 | 0.66 | 0.04 |
| *F. anerythrus from the left bank* | | | | | | | |
| M1 | 84 | 41.05 | 50.75 | 47.26 | 2.65 | 1.63 | 0.03 |
| M2 | 83 | 38.55 | 47.6 | 44.48 | 2.26 | 1.5 | 0.03 |
| M3 | 83 | 34.33 | 42.3 | 39.26 | 1.87 | 1.37 | 0.03 |
| M4 | 83 | 18.58 | 24.15 | 22.42 | 0.72 | 0.85 | 0.03 |
| M5 | 82 | 2.6 | 3.97 | 3.28 | 0.09 | 0.3 | 0.09 |
| M6 | 84 | 11.1 | 13.47 | 12.2 | 0.29 | 0.54 | 0.04 |
| M7 | 84 | 12.3 | 15.39 | 13.67 | 0.38 | 0.62 | 0.04 |
| M8 | 83 | 9.95 | 13.14 | 11.75 | 0.37 | 0.61 | 0.05 |
| M9 | 71 | 23.22 | 28.1 | 25.96 | 0.99 | 0.99 | 0.03 |
| M10 | 83 | 3.37 | 5.1 | 4.16 | 0.18 | 0.42 | 0.1 |
| M11 | 83 | 7.35 | 10 | 8.4 | 0.17 | 0.41 | 0.04 |
| M12 | 83 | 9.06 | 12.5 | 10.55 | 0.27 | 0.52 | 0.04 |
| M13 | 83 | 2.07 | 2.9 | 2.49 | 0.03 | 0.18 | 0.07 |
| M14 | 82 | 2.39 | 4.9 | 3.52 | 0.23 | 0.48 | 0.13 |

Baelo et al., submitted (Supplementary data)

| | | | | | | | |
|---|---|---|---|---|---|---|---|
| M15 | 83 | 5.75 | 8.06 | 6.95 | 0.25 | 0.5 | 0.07 |
| M16 | 83 | 10.5 | 14.5 | 12.76 | 0.52 | 0.72 | 0.05 |
| M17 | 83 | 7.25 | 9.3 | 8.08 | 0.12 | 0.35 | 0.04 |
| M18 | 84 | 8.62 | 10.85 | 9.54 | 0.14 | 0.38 | 0.03 |
| M19 | 84 | 14.53 | 19.26 | 17.76 | 0.88 | 0.94 | 0.05 |
| M20 | 83 | 2.06 | 3.2 | 2.35 | 0.04 | 0.2 | 0.08 |
| M21 | 83 | 10 | 12.68 | 11.38 | 0.29 | 0.54 | 0.04 |
| M22 | 83 | 9.05 | 12.3 | 11.09 | 0.31 | 0.55 | 0.05 |
| M23 | 76 | 14.11 | 17.32 | 15.68 | 0.52 | 0.72 | 0.04 |
| *F. cf bayonii* | | | | | | | |
| M1 | 36 | 40.91 | 43.87 | 42.24 | 0.8 | 0.89 | 0.02 |
| M2 | 36 | 38.6 | 41.47 | 40 | 0.56 | 0.75 | 0.02 |
| M3 | 36 | 33.04 | 36.4 | 34.75 | 0.67 | 0.82 | 0.02 |
| M4 | 36 | 18.65 | 21.01 | 19.75 | 0.38 | 0.61 | 0.03 |
| M5 | 36 | 2.18 | 3.55 | 2.58 | 0.09 | 0.3 | 0.12 |
| M6 | 36 | 9.3 | 11.04 | 10.08 | 0.23 | 0.48 | 0.05 |
| M7 | 36 | 10.3 | 12.56 | 11.37 | 0.22 | 0.46 | 0.04 |
| M8 | 36 | 9.91 | 11.46 | 10.49 | 0.17 | 0.41 | 0.04 |
| M9 | 36 | 21.5 | 24.8 | 23.07 | 0.37 | 0.61 | 0.03 |
| M10 | 36 | 3 | 4.79 | 3.74 | 0.23 | 0.48 | 0.13 |
| M11 | 36 | 6.75 | 8.33 | 7.57 | 0.13 | 0.36 | 0.05 |
| M12 | 36 | 8.82 | 10.7 | 9.66 | 0.2 | 0.45 | 0.05 |
| M13 | 36 | 2.1 | 2.8 | 2.45 | 0.04 | 0.19 | 0.08 |
| M14 | 36 | 1.83 | 4.21 | 2.64 | 0.29 | 0.54 | 0.2 |
| M15 | 36 | 5.63 | 7.5 | 6.51 | 0.25 | 0.5 | 0.08 |
| M16 | 36 | 9.38 | 12.2 | 10.48 | 0.4 | 0.63 | 0.06 |
| M17 | 36 | 6.98 | 8.34 | 7.62 | 0.14 | 0.37 | 0.05 |
| M18 | 36 | 8.5 | 9.74 | 9.03 | 0.08 | 0.28 | 0.03 |
| M19 | 36 | 14.17 | 17.26 | 15.42 | 0.59 | 0.77 | 0.05 |
| M20 | 36 | 1.68 | 3.29 | 2.36 | 0.09 | 0.3 | 0.13 |
| M21 | 36 | 9.21 | 11.25 | 10.19 | 0.36 | 0.6 | 0.06 |
| M22 | 36 | 8.93 | 11.35 | 9.96 | 0.4 | 0.63 | 0.06 |
| M23 | 32 | 13.78 | 15.96 | 15.09 | 0.26 | 0.51 | 0.03 |

Baele et al., submitted (Supplementary data)

### Supplementary figures

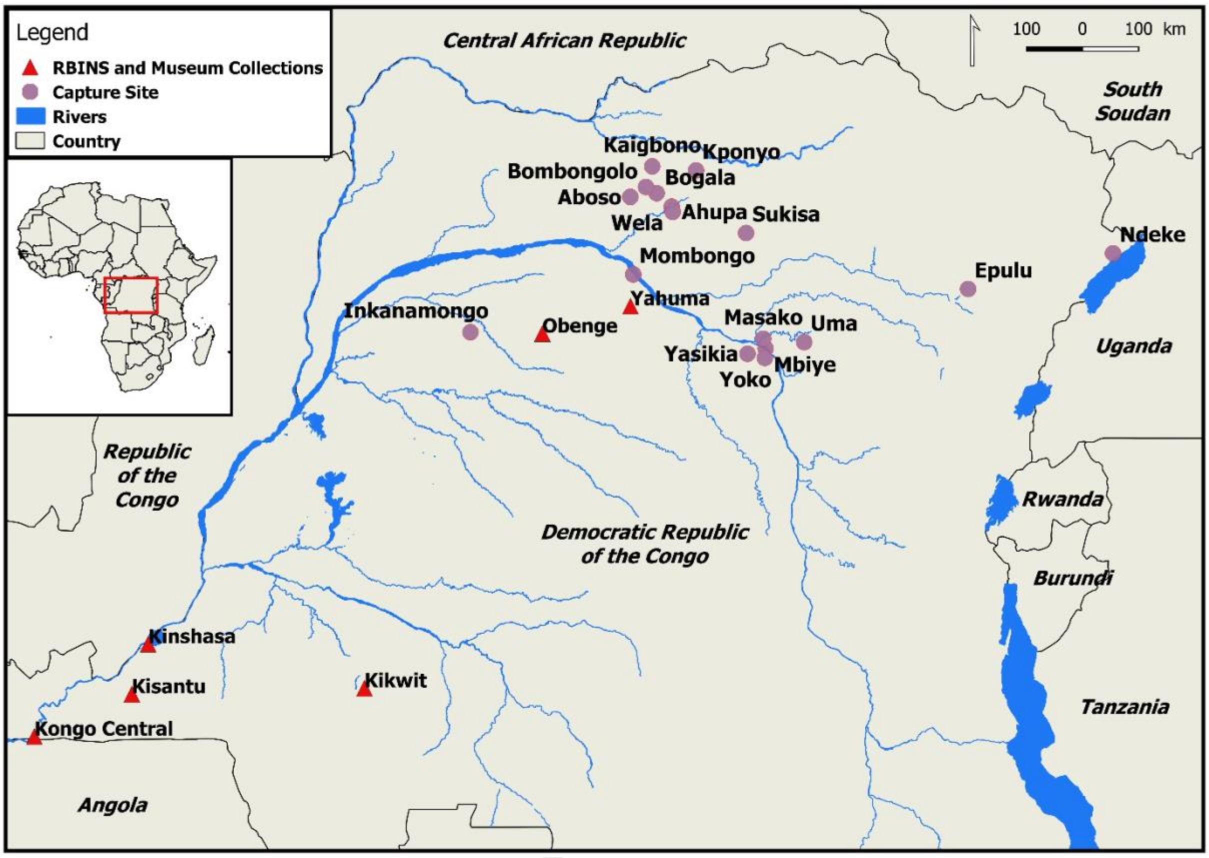


**Figure S1.** Geographical location of collection sites and localities from museum collection (specimens). GenBank sequences were obtained from individuals captured near Kinshasa and Kaigbono. RBINS and Museum Collections: specimens obtained from the Royal Belgian Institute for Natural Sciences and Royal Museum of Central Africa.

Baelo et al., submitted

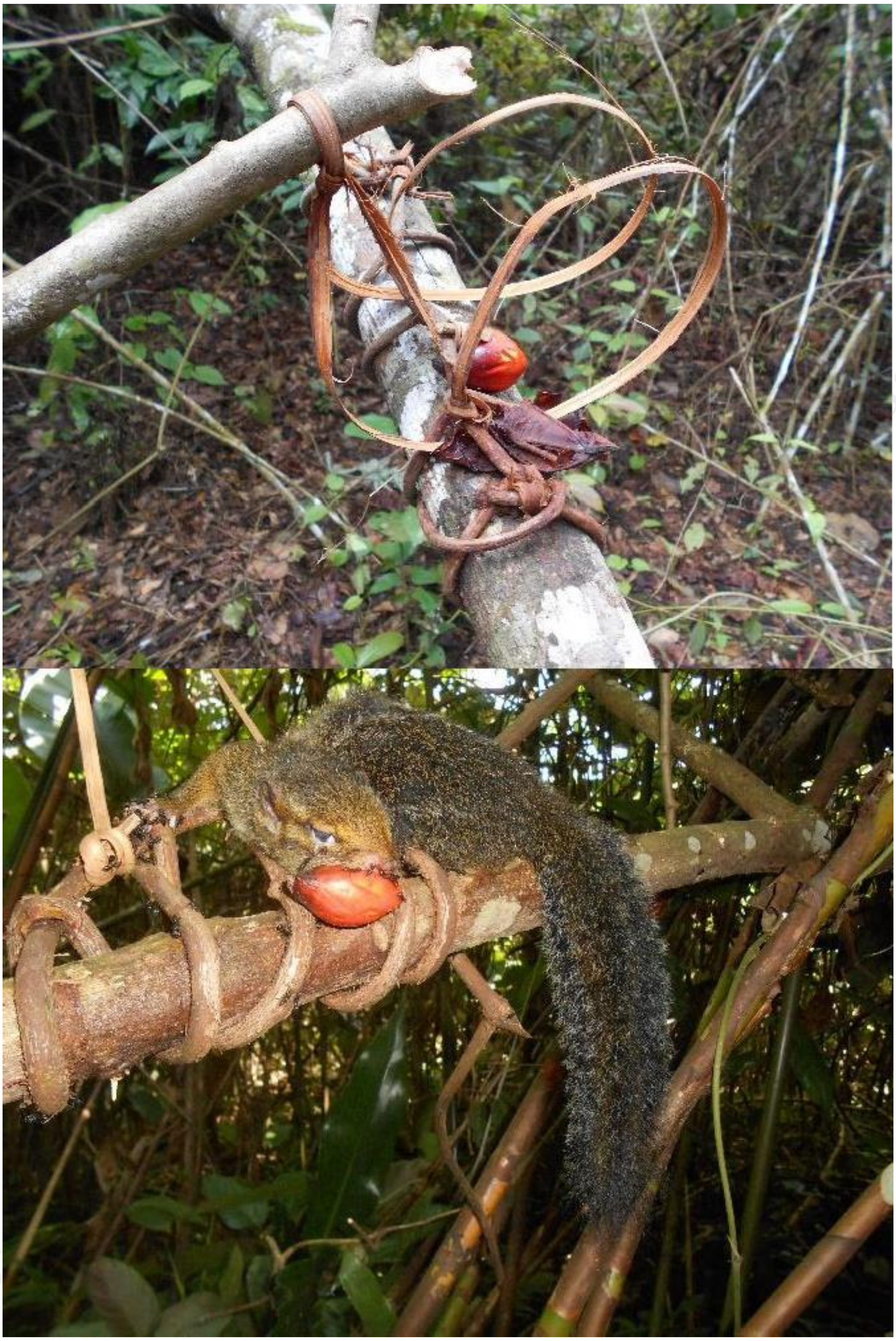

**Figure S2**. Traditional trap baited with palm nuts. On the right, a captured rope squirrel. (*Funisciurus sp*.).

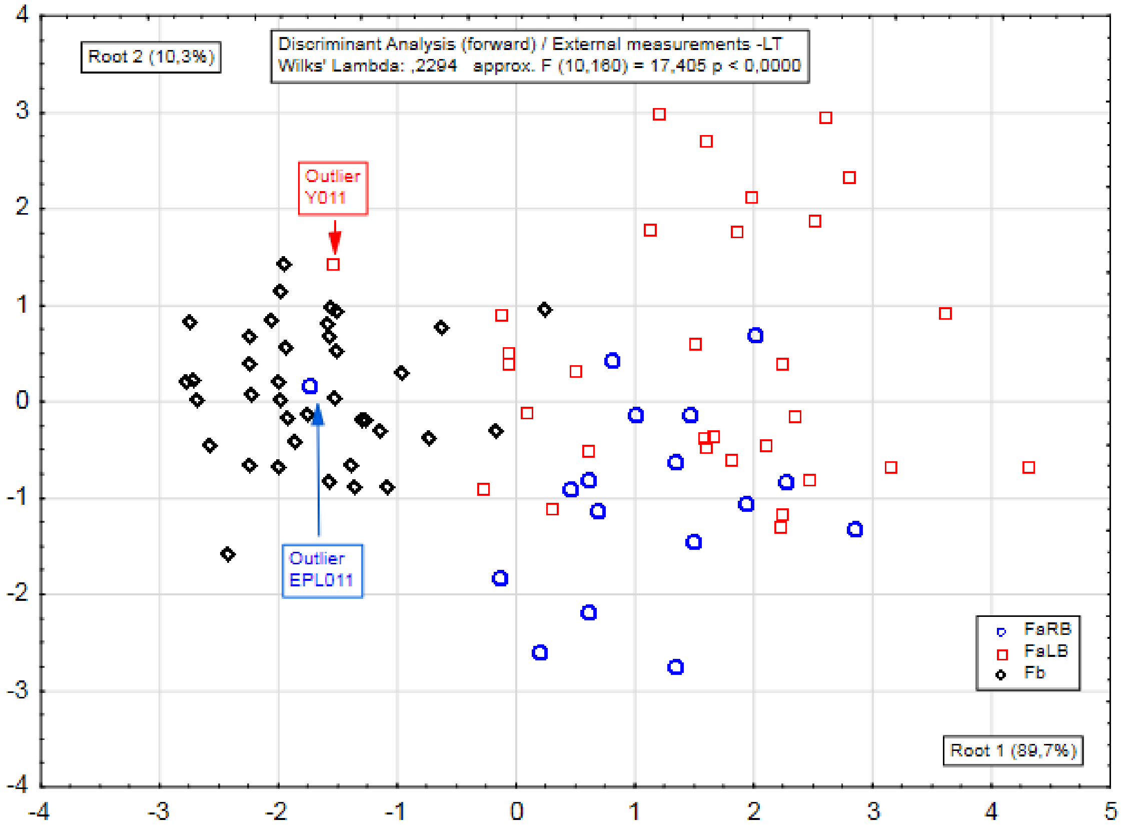


**Figure S3.** A discriminant analysis on the external measurements of FaRB, FaLB and FB. Since [Total length = Body length + Tail Length] Total length was not included in the analysis. FaRB = *F. anerythrus* Right Bank, FaLB = *F. anerythrus* Left Bank, FB = *F.cf bayonii*

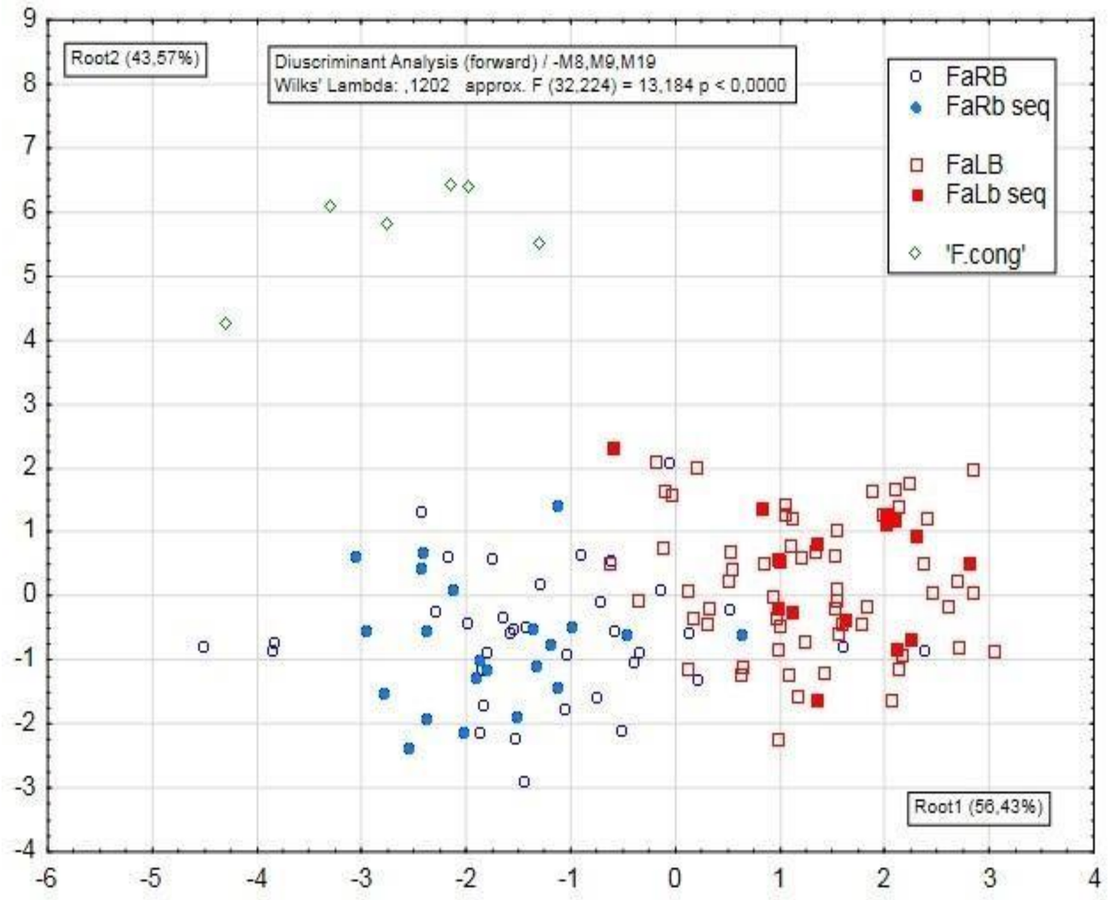


**Figure S4.** Discriminant analysis showing the OTUs FaLB, FaRB, and *F.* '*congicus*', using a partial craniometric dataset (FaRb : *F. anerythrus* Right Bank of Congo River; FaLb : *F. anerythrus* Left Bank; FbLb : *F.cf bayonii* Left Bank; F.cong : *Funisciurus cf congicus*; seq. : sequenced Most of the measured skulls of F.cong are damaged, so only six specimens are complete

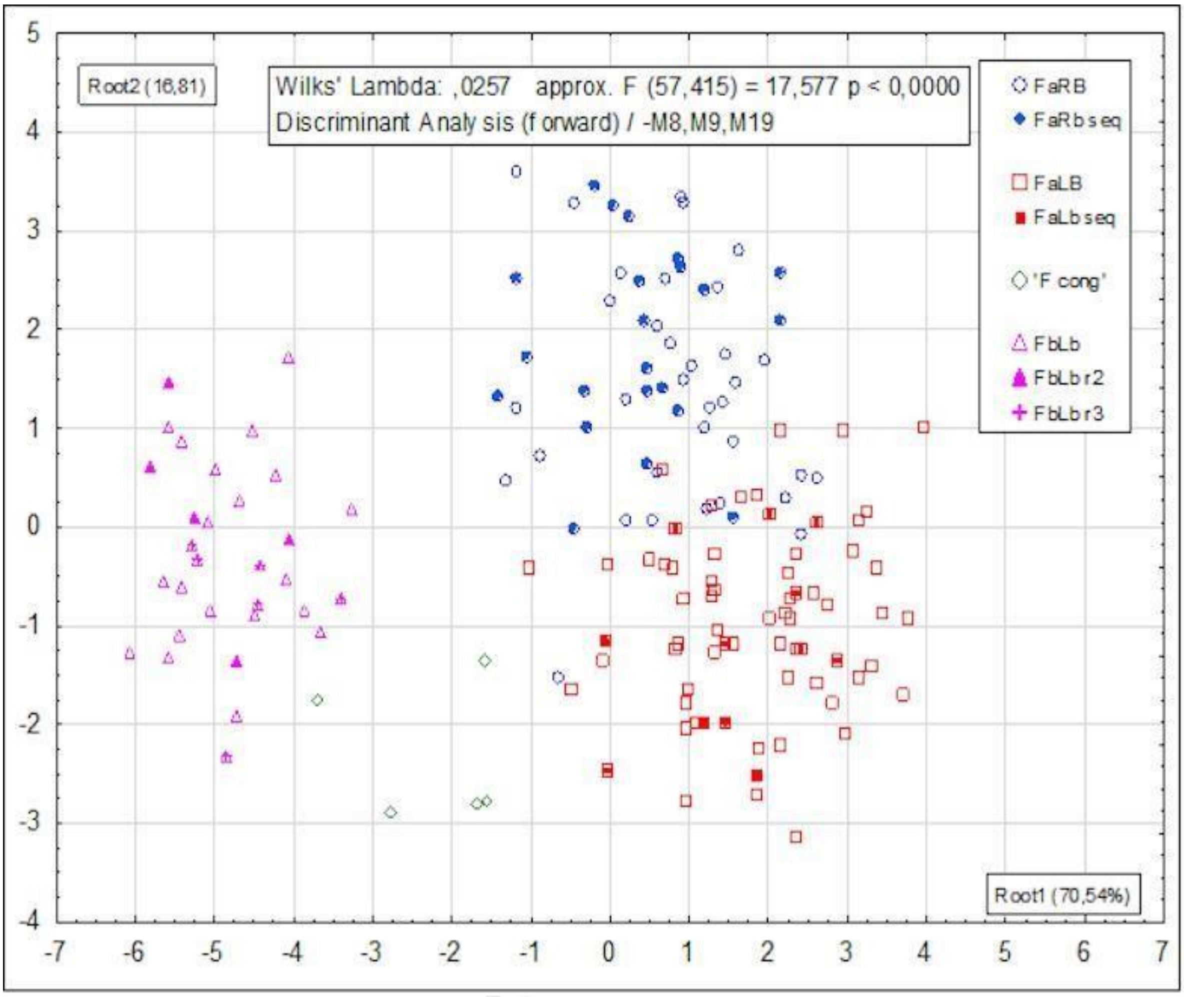


**Figure S5**. A forward discriminant analysis utilizing 20 variables (M8, M9 and M19 were omitted to maximize the number of observations of ‘F cong’) with sequenced specimens indicated by fully colored symbols. FaRb: *F. anerythrus* Right Bank of Congo River; FaLb: *F. anerythrus* Left Bank; FbLb: *F.cf bayonii* Left Bank; seq.: sequenced.

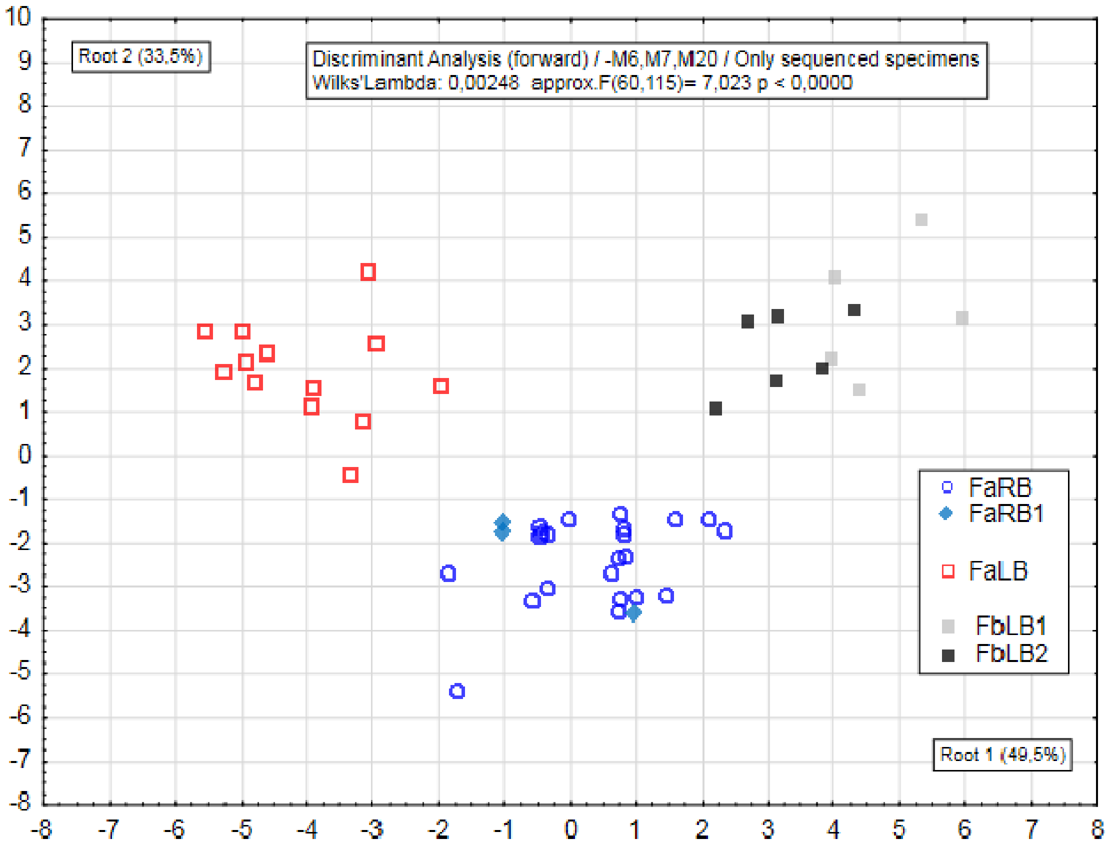


**Figure S6**. A discriminant analysis centered on the OTUs FaLb, FaRb, and Fb, utilizing a craniometric dataset limited to skulls from specimens with corresponding genetic sequences. This approach highlights subtle differences within the Fb group, though the limited availability of specimens with both sequence data and craniometric measurements has hindered the separation of these potentially distinct OTUs (FaRB : *F. anerythrus* right bank; FaRB; *F. anerythrus*: 3 specimens sitting apart in Cytb tree (= Fx in Figure 5); FaLB : *F. anerythrus* left bank; FbLB1 and FbLB2 : *F.cf bayonii*, differentiatied in Cytb tree as lineages 1 and 2).

Bueno et al., submitted

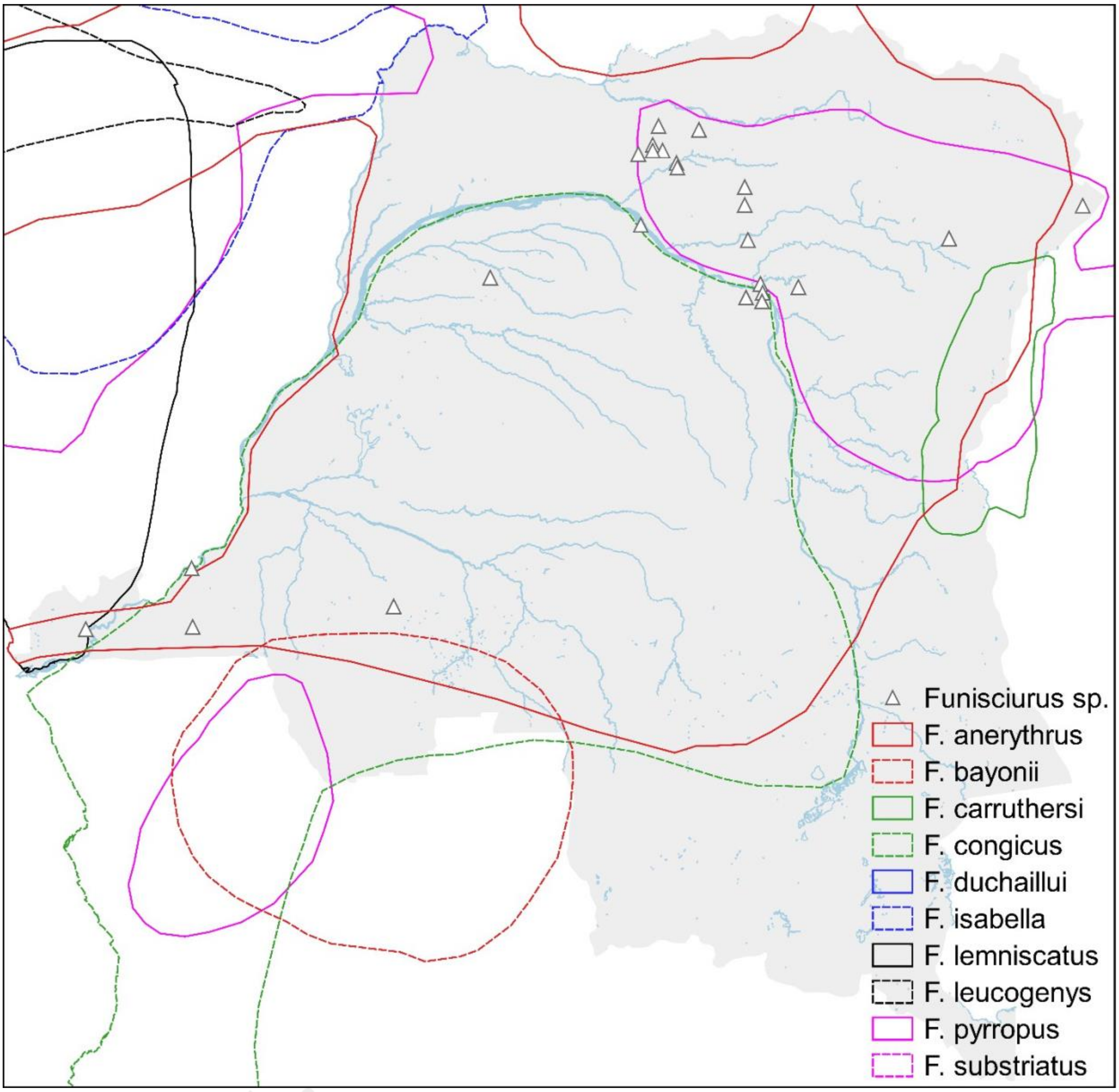


**Figure S7.** Polygon of extent occurrence of ten species of African rope squirrels. The data are available in the IUCN Red List of Threatened Species, v.2024-2; https://www.iucnredlist.org/

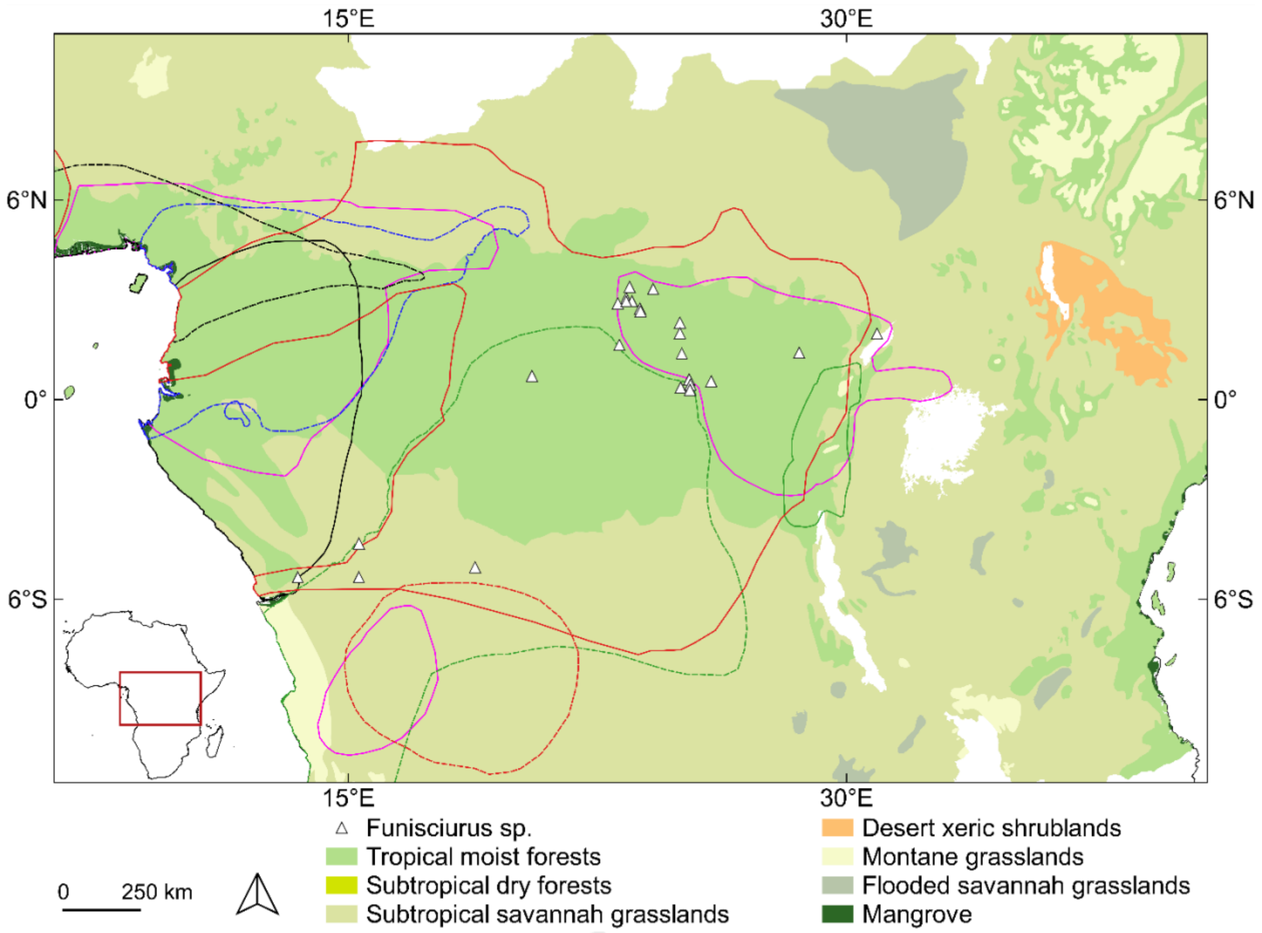


**Figure S8.** Geographic distribution of samples; extent of occurrence of ten African rope squirrel species (see Fig.2 for colour of polygon line); biotic zones from the Africa Biomes Dataset (https://opendata.rcmrd.org/).

Baelo et al., submitted
